# Gene syntax defines supercoiling-mediated transcriptional feedback

**DOI:** 10.1101/2025.01.19.633652

**Authors:** Christopher P. Johnstone, Kasey S. Love, Sneha R. Kabaria, Ross D. Jones, Albert Blanch-Asensio, Deon S. Ploessl, Emma L. Peterman, Rachel Lee, Jiyoung Yun, Conrad G. Oakes, Christine L. Mummery, Richard P. Davis, Brandon J. DeKosky, Peter W. Zandstra, Kate E. Galloway

**Affiliations:** Department of Chemical Engineering, MIT, 25 Ames St., Cambridge, MA, 02139, USA; Department of Biological Engineering, MIT, 25 Ames St., Cambridge, MA, 02139, USA; School of Biomedical Engineering, UBC, 6088 University Boulevard, Vancouver, BC, V6T 1Z3, Canada; Michael Smith Laboratories, UBC, 2185 East Mall, Vancouver, BC, V6T 1Z4, Canada; Department of Anatomy and Embryology, Leiden University Medical Center, 2300RC Leiden, the Netherlands; The Novo Nordisk Foundation Center for Stem Cell Medicine, reNEW, Leiden University Medical Center; Department of Bioengineering, California Institute of Technology, Pasadena, CA, 91125, USA; The Ragon Institute of Mass General, MIT, and Harvard, 600 Main St., Cambridge, MA, 02139, USA

## Abstract

Gene syntax—the order and arrangement of genes and their regulatory elements—shapes the dynamic coordination of both natural and synthetic gene circuits. Transcription at one locus profoundly impacts the transcription of nearby adjacent genes, but the molecular basis of this effect remains poorly understood. Here, using integrated reporter circuits in human cells, we show that the reciprocal effects of transcription and DNA supercoiling, which we term supercoiling-mediated feedback, regulates expression of adjacent genes in a syntax-specific manner. Using a suite of chromatin state assays, we measure syntax-and induction-dependent formation of chromatin structures in human induced pluripotent stem cells. Applying syntax as a design parameter and without altering sequence or copy number, we built compact gene circuits, tuning the expression mean, noise, and stoichiometry across diverse delivery methods and cell types. Integrating supercoiling-mediated feedback into models of gene regulation will expand our understanding of native systems and enhance the design of synthetic gene circuits.

## 1 Main

Native gene circuits coordinate transcriptional programs that orchestrate complex cellular processes across space and time. Circuits that require precise transcriptional regulation in development, such as *Hox* genes [1, 2] and segmentation clocks [3], co-localize multiple transcriptional units within tens of kilobases. By harnessing local chemical and physical forces, the organization of these neighboring genes may support their coordinated expression (fig. 1a) [4–6]. Gene syntax specifies the relative order and orientation of adjacent genes. Pairs of genes can adopt a tandem, divergent, or convergent syntax, which are predicted to generate distinct profiles of gene coupling and dynamics [7]. These predicted profiles differ in both mean expression and in levels of transcriptional noise. In native genomes, syntaxes are not equally represented across gene pairs (figs. 1b and S1). For instance, in the human genome, closely spaced gene pairs more commonly adopt a divergent syntax. Potentially, gene syntax may couple expression of adjacent genes and tune transcriptional noise.

**Figure 1:**
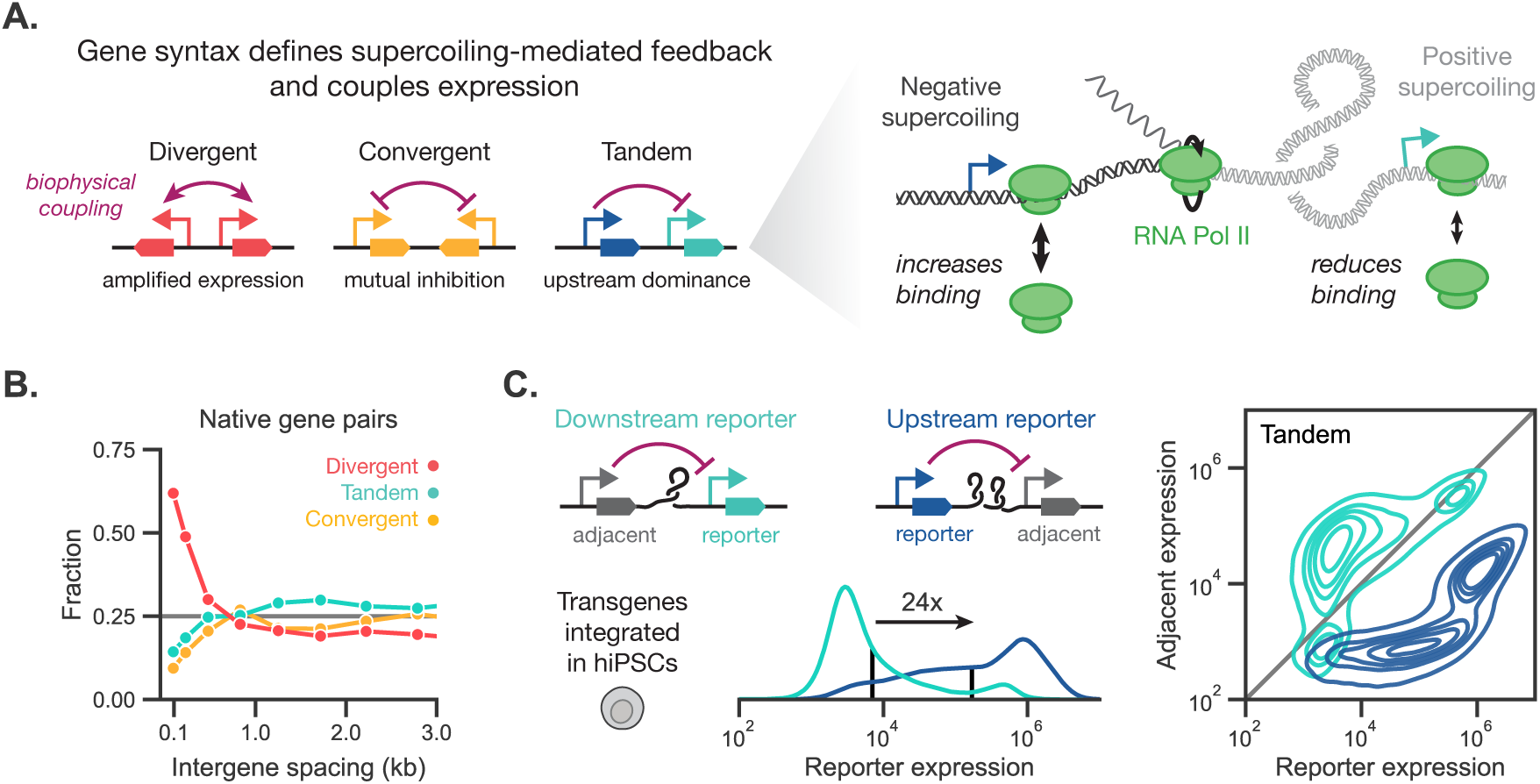
Supercoiling-mediated feedback couples expression of adjacent genes. a) Gene pairs can be arranged in divergent, convergent, or tandem syntax, which specifies the biophysical coupling (purple arrows) between genes through supercoiling-mediated feedback. DNA supercoiling modifies the energy required for polymerases to bind and locally melt DNA, leading to increased or decreased polymerase initiation. Transcribing polymerases generate negative supercoiling upstream and positive supercoiling downstream, which can diffuse to nearby genes. This supercoiling takes the form of *twist*, where supercoiling changes the average rotation per base pair, or *writhe*, where supercoiling forms large-scale twisted loops, freely interconverting between the two forms. Together, transcription affects and is affected by supercoiling, forming a biophysical feedback loop that depends on the direction of transcription and thus on gene syntax. b) Using Ensembl annotations for the human genome, gene extents were identified using the maximum extent of all annotated exons, then the gene syntax and intergene spacing were determined for each pair of adjacent genes. Gene pairs were split into equal-quantile bins based on intergene spacing, and the fraction of gene pairs with each syntax was computed for each bin. c) Due to accumulation of positive supercoiling at the downstream promoter, expression from an upstream gene is predicted to decrease expression of a downstream gene. Two-gene constructs expressing fluorescent proteins were integrated using PiggyBac in hiPSCs, where the adjacent gene is expressed from a weak PGK promoter and the reporter gene is expressed from a strong EF1α promoter. Left: For a representative biological replicate, the distribution of reporter expression is shown as a function of position in the circuit. Geometric means are shown as black vertical lines, and the fold change between the two circuits is annotated. Right: Distributions of reporter and adjacent gene expression for representative replicates of the same tandem circuits. The gray diagonal represents equal adjacent and reporter signal; the two circuits are approximately symmetric to each other across this line.

Gene syntax affects expression of colocalized genes through biophysical forces such as DNA supercoiling [7, 8]. DNA supercoiling, or the over-and under-twisting of double-stranded DNA, both influences and is generated by transcription in a directional manner. Mechanistically, transcribing RNA polymerases induce waves of supercoiling by melting the double helical DNA polymer to read the underlying base pairs. According to the twin domain model, polymerases generate negative supercoiling upstream and positive supercoiling downstream [9, 10]. This supercoiling can take the form of *twist*, where supercoiling changes the average rotation per base pair, or *writhe*, where supercoiling forms large-scale intertwined loops. In turn, supercoiling impacts gene expression by influencing transcriptional bursting [11–13], polymerase stalling [14, 15], topoisomerase activity [16–21], and chromatin folding [22–25]. In particular, supercoiling may alter RNA polymerase binding; negative supercoiling reduces the binding energy and facilitates polymerase loading, while positive supercoiling increases the energy barrier and reduces binding rates [26]. As supercoiling diffuses along the DNA polymer, the directionality of transcription sets the energy landscape for subsequent polymerase binding events (fig. 1a). Thus, models of supercoiling predict that transcription-induced changes in chromatin structure feed back into changes in transcriptional activity at adjacent genes [27–30], a phenomenon we define as supercoiling-mediated feedback [7]. Because polymerase directionality governs supercoiling generation, gene syntax defines supercoiling-mediated feedback, setting syntax-specific profiles of gene expression [7]. Patterns of supercoiling around actively transcribed genes have been observed and averaged genome-wide in bacteria [31, 32], in eukaryotes [11, 22, 24, 31, 33], and in human cells [17, 22, 25]. However, the impact of transcriptionally induced supercoiling on fine-scale chromatin structure and syntax-specific patterns of expression remain uncharacterized in human cells.

DNA supercoiling is often studied through broad perturbations including the loss or inhibition of topoisomerases and polymerases [11, 17, 25, 33, 34], which induce large changes in cellular physiology and limit observations to acute treatments and short timescales. Alternatively, synthetic two-gene systems offer a well-controlled testbed to independently vary gene syntax, while keeping gene identity, circuit copy number, genomic location, and other parameters constant. Modulating the transcriptional activity of a single transgene within this genetically uniform background offers the precise control required to examine the predictions of supercoiling-mediated feedback using chromatin structure techniques.

Beyond advancing our understanding of native gene regulation, defining syntax-specific expression profiles may enhance design of synthetic gene circuits. Synthetic circuits offer programmable control of therapeutic cargoes, genome editors, and cell fate [35–43]. However, predictable forward design of gene circuits remains challenging, requiring iterative “design-build-test” loops to achieve desired functions. Synthetic circuits on the length scale of ∼10 kb are regularly integrated into the genome and thus are subject to biophysical forces, including DNA supercoiling, that may rapidly couple expression of their components [7, 8]. Harnessing syntax as an explicit design parameter may improve the predictability and performance of genome-integrated synthetic circuits [7].

Here, we use synthetic two-gene circuits as a model system to examine how transcription of a single gene couples the expression and chromatin structure of adjacent genes in human cells. Leveraging the control and orthogonality of inducible systems, we demonstrate that transcription-induced coupling generates syntax-specific profiles of expression across a range of human cell types and integration methods. To investigate the corresponding changes in chromatin, we employ a suite of genomics techniques to characterize the structural and biochemical properties of the region surrounding the locus of circuit integration in human induced pluripotent stem cells (hiPSCs). Induction of transcription perturbs chromatin structure both at the circuit and hundreds of kilobases away, substantially altering the insulation, RNA Pol II binding profile, and histone marks across the locus. Applying these principles of supercoiling-mediated feedback, we design compact synthetic gene circuits for efficient delivery, tunable expression levels and noise, and robust induction across a variety of cells. These techniques allow us to optimize production of a therapeutic antibody without substitution of genetic parts. Overall, we demonstrate how supercoiling-mediated feedback influences expression of adjacent genes, providing insights into native gene regulation and informing the design of synthetic systems.

### Upstream dominance defines expression profiles of constitutive tandem transgenes

In native genes, upstream transcription can reduce expression of downstream genes [44–46]. Models of supercoiling-mediated feedback predict that positive supercoiling generated by transcription at the upstream gene reduces the rate of transcription and thus expression at the downstream locus, resulting in upstream dominance (fig. 1a) [7]. To examine upstream dominance in a synthetic system, we constructed two-gene systems in tandem. Each gene is paired with a promoter and polyadenylation signal (PAS) to form a transcriptional unit. The modularity of these synthetic systems allows us to independently switch gene positions and regulatory elements.

To measure expression, we integrated these tandem two-gene systems into two common human cell lines, HEK293Ts and hiPSCs, via PiggyBac transposase. We switch the positions of the tandem genes to isolate the effect of position on expression level. We quantified expression of the fluorescent reporter genes by flow cytometry, providing single-cell resolution needed to measure expression distributions. We found that gene position strongly influences the expression of the reporter. Even when pairing a strong promoter with a weak promoter in hiPSCs, we saw clear upstream dominance with a large, 24-fold shift in expression based on position (fig. 1c). In HEK293Ts, the gene in the upstream position expresses at levels nearly four-times higher than when placed downstream (fig. S2). Potentially, genetically encoded sequences—such as binding sites for the CCCTC-binding factor (CTCF) [47, 48] and the cHS4 insulator [49]—that restrict chromatin-mediated interactions may reduce upstream dominance. To examine this hypothesis, we inserted tandem-oriented CTCF binding sites, native motifs previously identified to reduce enhancer-promoter interactions [48, 50]. These sequences have been used to flank integrated synthetic circuits to minimize impact of the integration site on circuit expression. Flanking the upstream gene, the downstream gene, or the entire two-gene construct with these sites does not eliminate upstream dominance (fig. S3a). Other known insulator sequences placed between the two genes did not reduce upstream dominance relative to an inert spacer (fig. S3b). Instead, addition of these sites generally reduces expression of one or both genes.

The identity of regulatory elements such promoters and PAS may influence coupling between genes [51, 52]. In testing a panel of common constitutive promoters, we consistently observed upstream dominance in PiggyBac-integrated HEK293Ts (figs. S4a and S4b), lentivirally integrated HEK293Ts (fig. S4c), and PiggyBac-integrated hiPSCs (fig. S5). In exchanging the PAS, we found that the choice of PAS has a minimal effect on the shape of expression distributions, mildly tuning the levels of gene expression (fig. S6). Thus, trends in syntax-specific expression are robust across a range of genetic parts for the tandem syntax.

### Transcription of an adjacent gene induces syntax-specific coupling

Expression patterns of constitutively expressed tandem gene pairs suggest that syntax influences expression. However, these constitutive systems do not support dynamic control of transcription of a single adjacent gene, making it difficult to parse the transcription-driven mechanism of supercoiling-mediated feedback. To allow controlled induction of a single adjacent gene, we generated monoclonal HEK293T cell lines containing a doxycycline (dox)-inducible two-gene system in different syntaxes (fig. 2). To build these lines, we placed a constitutive reporter gene under the control of a strong constitutive promoter (EF1α) and an adjacent inducible gene under the control of the dox-inducible promoter TRE (fig. 2a). Using PiggyBac, we delivered the dox-inducible two-gene systems encoded in tandem, convergent, and divergent syntaxes, which are predicted to show different transcription-induced couplings [7]. The constitutively expressed dox-responsive activator, rtTA, was integrated from a separate PiggyBac donor. We sorted single cells to establish monoclonal lines of each syntax.

**Figure 2:**
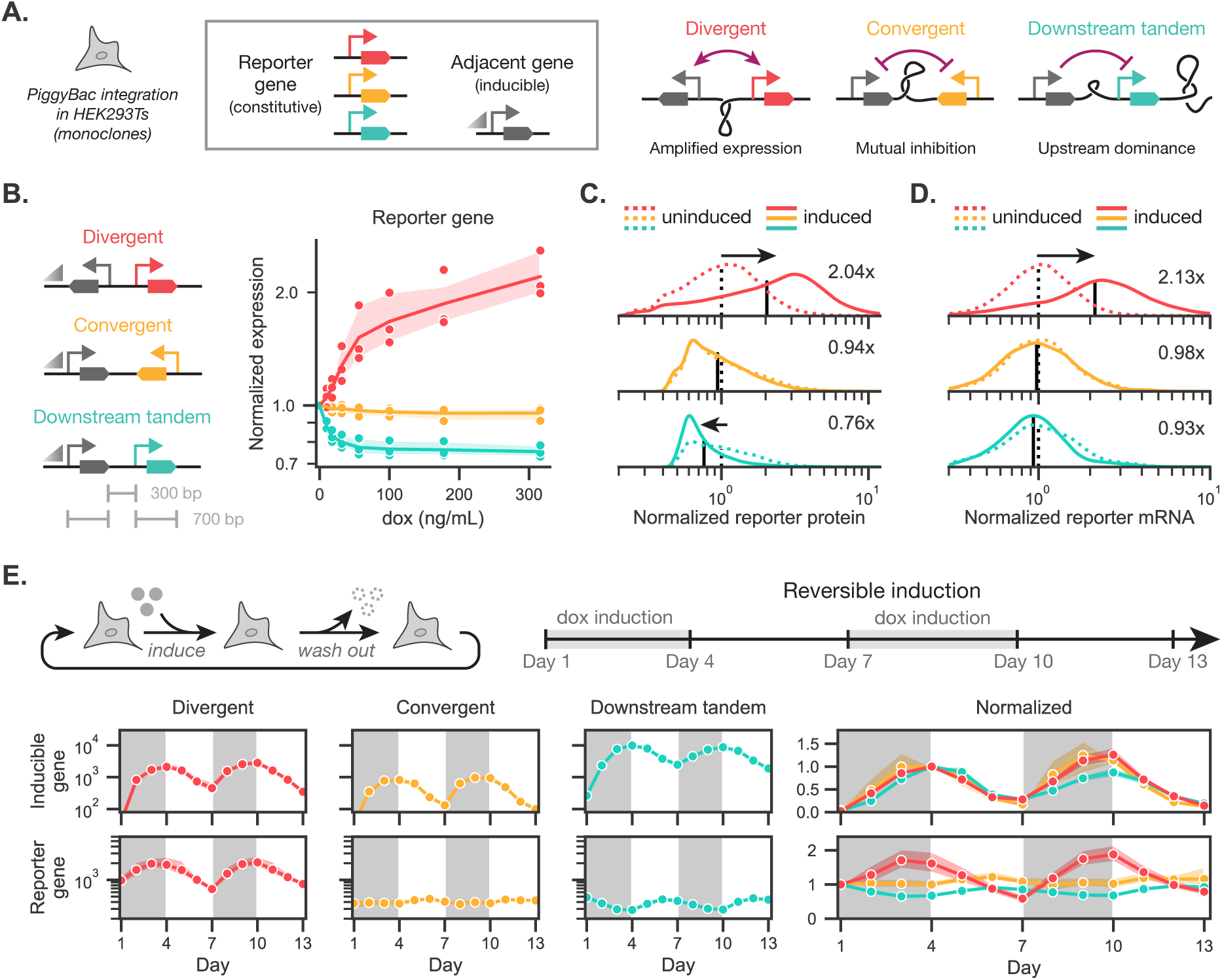
Transcription induces syntax-specific coupling of expression of adjacent genes. a) Two-gene systems consisting of an dox-inducible gene (∼700 bp, TRE promoter) and a constitutively expressed gene (∼700 bp, EF1α promoter) spaced ∼300 bp apart were integrated into HEK293Ts using PiggyBac. The resulting cell lines were flow-sorted to single cells and expanded as monoclonal populations. b) The geometric mean reporter expression, normalized to the uninduced condition, is shown as a function of dox concentration for the three different syntaxes. Geometric mean and associated 99% confidence interval are shown for three separately passaged and maintained replicates of the monoclonal cell lines. c)-d) Full reporter protein and mRNA distributions are shown in the uninduced case and the second-highest dox induction state. Geometric means are shown as black vertical lines, and the fold changes upon induction are annotated. e) Dox was sequentially introduced and removed in order to measure the turn-on and turn-off dynamics of the integrated systems. The systems respond reversibly to the presence of dox. Geometric mean and associated 99% confidence interval are shown for four separately passaged and maintained replicates of the monoclonal cell lines.

Upon dox addition, all syntaxes show strong induction of the TRE-driven inducible gene (figs. S7a and S7b). To quantify changes in expression of the constitutive reporter upon induction of the adjacent gene, we normalized reporter expression to the uninduced condition for each line. In the tandem syntax, induction of the upstream gene reduces expression of the downstream reporter gene (fig. 2b, figs. S7c and S7d). Conversely, induction of the divergent syntax strongly upregulates expression from the constitutive reporter, matching predictions of amplification in divergent syntax [7]. The convergent syntax shows a near-invariant profile of reporter expression. For the tandem and divergent syntaxes, induction of the adjacent gene results in a unimodal shift in the geometric mean (fig. 2c). Unimodal shifts indicate a general mechanism of regulation, such as changes in the transcription rate, that is not restricted to a subpopulation of cells.

As a transcription-based process, supercoiling-mediated feedback should manifest in the distributions of mRNAs. To measure the mRNA distributions, we used single-cell hybridization chain reaction RNA-FISH [52, 53] to quantify the transcriptional profiles of both the constitutive reporter gene and the dox-inducible gene (fig. 2d, figs. S7e, S7f and S8). As expected, mRNA profiles generally match the syntax-specific profiles of proteins (fig. 2c). Intriguingly, for the convergent syntax, the inducible gene exhibits bimodal mRNA expression (fig. S7), matching modeling predictions of bimodality that may be obscured by stable protein reporters [7]. Bimodality in mRNA expression for the constitutive reporter may not be visible due to the overall low expression of this gene in convergent syntax. Overall, the mRNA profiles align with the models of supercoiling-mediated feedback that predict that syntax influences transgene expression by altering rates of transcription. DNA supercoiling-mediated coupling is predicted to be rapid and reversible. To test the reversibility of syntax-specific coupling, we sequentially induced and removed dox for three-day periods over 13 days. We observed repeatable induction-and syntax-specific coupling of the inducible and reporter genes. Over 13 days we observed minimal hysteresis, indicating that the trends in coupling are reversible and are not due to permanent changes in the chromatin state (fig. 2e).

### Transcription induces syntax-specific structures in the neighborhood of synthetic circuits

Transcription and transcription-induced supercoiling drives genome folding [54]. Just as we used synthetic inducible circuits to probe how gene syntax affects gene expression (fig. 2), these circuits provide a facile mechanism to regulate transcriptional activity and measure transcriptionally induced chromatin structures.

Measuring both 1D and 3D chromatin profiles requires site-specific, homozygous integration of the synthetic circuit for downstream next-generation sequencing (NGS). Site-specific integration allows circuits to be measured at a fixed location, and homozygous integration ensures reads can be accurately interpreted without allelic heterogeneity. Using the STRAIGHT-IN Dual allele platform [55], we integrated dox-inducible circuits placed in all four possible two-gene syntaxes into both alleles of a genomic safe harbor region located in intron 2 of the citrate lyase beta-like (*CLYBL*) gene [56] in hiPSCs, generating homozygous cell lines (fig. S10a). These circuits include the activator within the constitutively expressed gene. We confirmed that all four syntaxes showed reversible, syntax-specific profiles of expression, including divergent-mediated amplification and tandem-mediated upstream dominance, similar to those observed in the dox-inducible two-gene systems integrated into HEK293T cells (figs. 3b and S10b). Examining the single-cell mRNA distributions (figs. S10c to S10e), we found that most transcript levels matched the observed protein level trends. However, the induced downstream tandem syntax showed high levels of RNA and low protein expression for the constitutive gene, potentially indicating a large fraction of unproductive transcripts (fig. S10e). To control for locus-specific effects, we integrated these constructs into another common safe harbor locus, *AAVS1*. *AAVS1* is also located within the intron of a transcriptionally active native gene (figs. S11a and S11b). Like at the *CLYBL* locus, we observe syntax-specific profiles of expression at *AAVS1* (fig. S11), including both divergent-mediated amplification of the constitutive gene and poor expression from the downstream inducible gene in the upstream tandem syntax (fig. S11c). We attribute the approximately two-fold drop in inducible gene expression at the *AAVS1* locus to interference from upstream native expression.

To understand how transcriptional activity and syntax affect the chromatin structure of our two-gene circuits, we used Region Capture Micro-C (RCMC) [57] to measure the contact probability between genomic locations within a targeted region of interest (fig. 3a). Notably, the *CLYBL* safe harbor integration site is located at the boundary of two topologically associating domains (TADs) (fig. 3a). By computing the fold-change in contact probability, we find that induction increases intra-TAD contacts and reduces inter-TAD contacts in the divergent and convergent syntaxes (figs. 3c and S14). Induction does not alter these contacts in the tandem syntaxes. Using a common sliding-window insulation score [58, 59] that becomes more negative at strong TAD boundaries, the overall change in contact probability in the divergent and convergent syntaxes occurs alongside a substantial weakening and strengthening, respectively, of the local TAD boundary (fig. 3d). No substantial change in this boundary is observed in the tandem syntaxes. This change in the local insulation may be caused by perturbation of a loop domain present at the integration site. Visible as a “corner dot”, the loop domain moves upon induction of the divergent and convergent syntaxes (fig. S13). In the divergent case, the loop domain splits to form a dual-loop domain. Putatively, this double-loop may be an averaged interaction between two stochastic loop domains influenced by polymerases traveling upstream from the divergent promoter.

**Figure 3:**
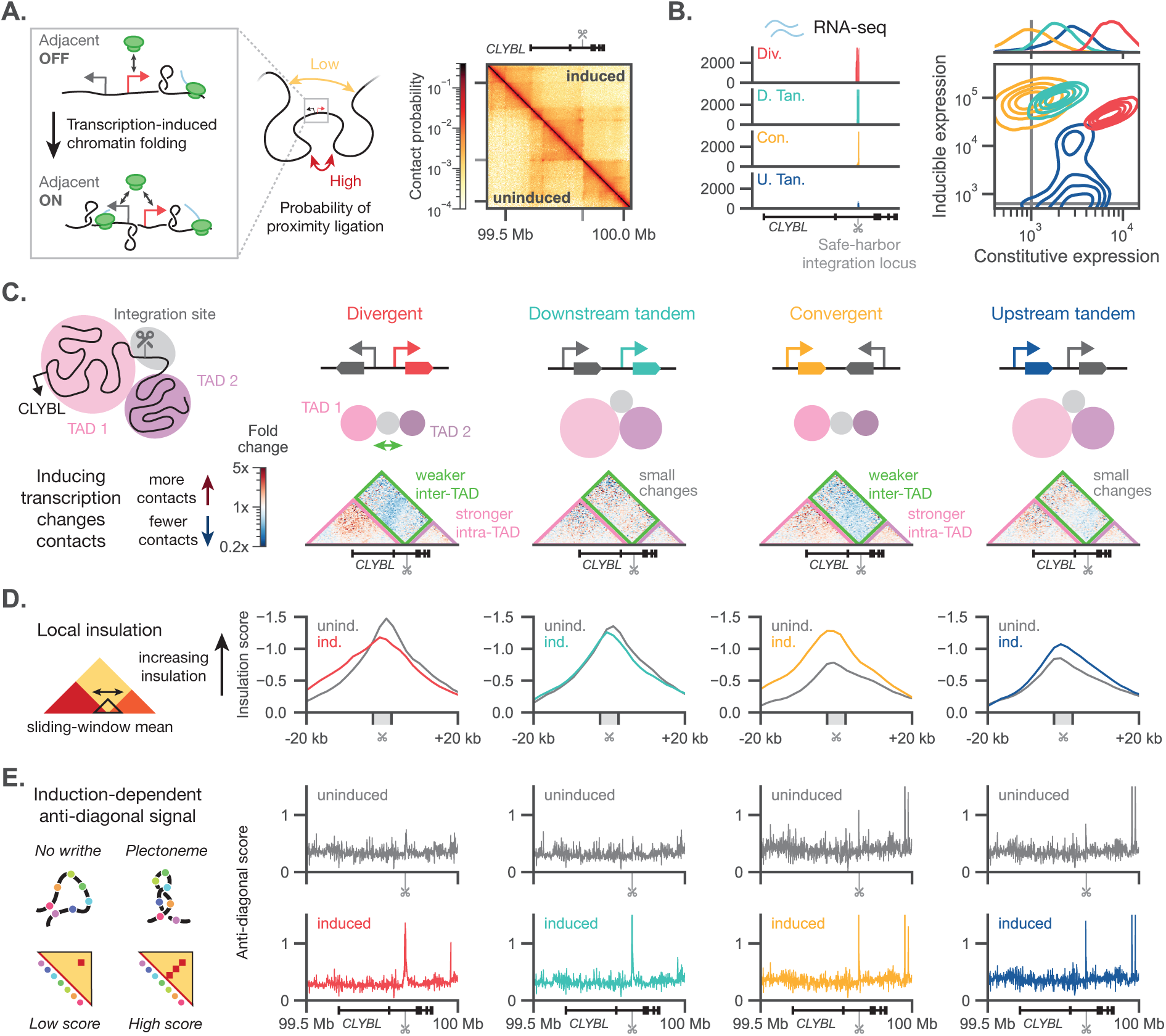
Transcription induces syntax-specific chromatin structures across synthetic gene circuits and the surrounding locus. a) Inducing transcription of an adjacent gene alters chromatin folding at the integration location and the surrounding region. 3D chromatin structure can be measured via Region Capture Micro-C (RCMC), which uses proximity ligation and downstream sequencing to quantify contact probability. We performed RCMC on uninduced and induced conditions for all four syntaxes of two-gene circuits. The circuits were homozygously integrated into a landing pad at the *CLYBL* locus in hiPSCs (fig. S10a). A representative contact map for the divergent uninduced and induced conditions is shown. Gray ticks indicate the integration location. b) Left: Bulk RNA-seq tracks are shown for the induced condition of the cell lines in a). Right: Joint distributions of protein expression for the constitutive and inducible genes are shown for the same conditions. Histograms for constitutive expression are also shown (top). Gray lines indicate expression thresholds set by the non-integrated, parental population. c) RCMC data was binned at 5-kb resolution and iteratively balanced within the capture region. For each syntax, the fold change in contact probability upon induction was computed. We observe that in the divergent and convergent syntaxes, inter-TAD contacts decrease while intra-TAD contacts increase. d) Using a sliding-window insulation score (window size of 6 kb), the local insulation around the integration site is computed. Stronger insulation gives a more negative insulation score. The divergent and convergent syntaxes show weaker and stronger local insulation, respectively. The tandem syntaxes only show small changes in local insulation. e) The anti-diagonal score, calculated as the sum of anti-diagonal contact probabilities out to a distance of 5 kb, is shown across the entire capture region. A high anti-diagonal score may indicate the presence of plectonemes, structures that form when DNA supercoiling twist converts to writhe. Strong anti-diagonal scores are present at the integration locus upon induction for all four gene syntaxes. No chromatin outside of the integration region shows a comparable induction-dependent anti-diagonal score.

Beyond changes in TAD strength, we looked for other induction-dependent signals in the contact probability dataset. Specifically, both overwound or underwound supercoiled DNA can buckle, forming “figure-eight” structures called plectonemes as DNA twist converts to writhe. Plectonemes facilitate the loading of chromatin loop extruders [60] and should appear as small-scale regions of high contact probability perpendicular to the diagonal, similar in shape to the large-scale “jets” generated by cohesion loop extrusion [61]. Quantifying these structures using an anti-diagonal score that sums contact probabilities along the anti-diagonal over 5 kb, we observe strong, induction-dependent plectonemic signals at the integrated locus in all four syntaxes (fig. 3e). Indeed, this signal correlates with the appearance of red anti-diagonal stripes in the contact matrix (fig. S19). The magnitude of these induction-dependent, putatively plectonemic signals is not seen elsewhere in the capture region (fig. S15), strongly suggesting that transcription reshapes chromatin folding around our synthetic circuit.

### Transcription drives syntax-specific profiles of supercoiling and changes in chromatin state

Supercoiling-mediated changes in gene expression coincide with changes in chromatin structure. In bacteria and in yeast, these 3D chromatin structures correlate with supercoiling density [34, 62]. Supercoiling-dependent structures should be accompanied by changes in both local supercoiling density and in epigenetic profiles demarcating active transcription. Our measurement of the anti-diagonal score (fig. 3e) is a suggestive but indirect measurement of supercoiling, as other structures such as chromatin jets also appear as anti-diagonal contacts. Thus, we used GapRUN to measure positive supercoiling density [17, 34], bulk RNA-seq to measure productive transcripts, and CUT&Tag to measure the concentration of both actively elongating RNA polymerases and of various transcription-linked histone marks.

Specifically, to complement our putative measurement of plectonemes (positive and negative writhe) via RCMC, we measured the amount of positive supercoiling twist. GapRUN uses the bacterial protein GapR to identify regions of positive supercoiling twist through an MNase-based NGS readout [17, 34]. Because supercoiling freely interconverts between twist and writhe in the regime where plectonemes form, the presence of both anti-diagonal RCMC contacts and GapR signal indicates overall positive supercoiling. We generally observe the formation of positive supercoils at the 3’ ends of active genes. In particular, positive supercoiling accumulates in the intergenic regions of tandem syntaxes (figs. 4 and S16), putatively the cause of upstream dominance. Measuring the polyadenylated transcripts via RNA-seq, we observe readthrough transcripts for the inducible gene, where the transcript extends past the polyadenylation signal. For the downstream tandem syntax, we observe unproductive constitutive transcripts (fig. S10e) that do not contribute to constitutive protein expression (fig. 3b). This readthrough effect, while not observed in our randomly integrated constructs (fig. 2), may explain some of the upstream dominance.

To quantify the transcriptional state of the locus, we measured the concentration of actively elongating RNA Pol II (serine 2 phosphorylation), active histone marks H3K27ac and H3K4me3, and H3K27me3, a mark of heterochromatin. As a control, these transcription-linked signals are present at low levels in the distal, unedited *AAVS1* locus (fig. S17), but appear at other highly transcribed loci (fig. S18). At our synthetically integrated locus, we find that the actively transcribed genes co-occur with elongating RNA Pol II and active histone marks (fig. 4). Induction strongly increases signs of activity across the locus for the divergent and downstream tandem syntaxes (fig. 4). The upstream tandem and convergent syntaxes show only minimal changes in these marks of active transcription upon induction (fig. S16), consistent with their poor expression of the circuit (fig. 3b). Across syntaxes, we observe small changes in the heterochromatin mark H3K27me3, indicating that epigenetic silencing does not explain the low expression in the upstream tandem and convergent syntaxes.

**Figure 4:**
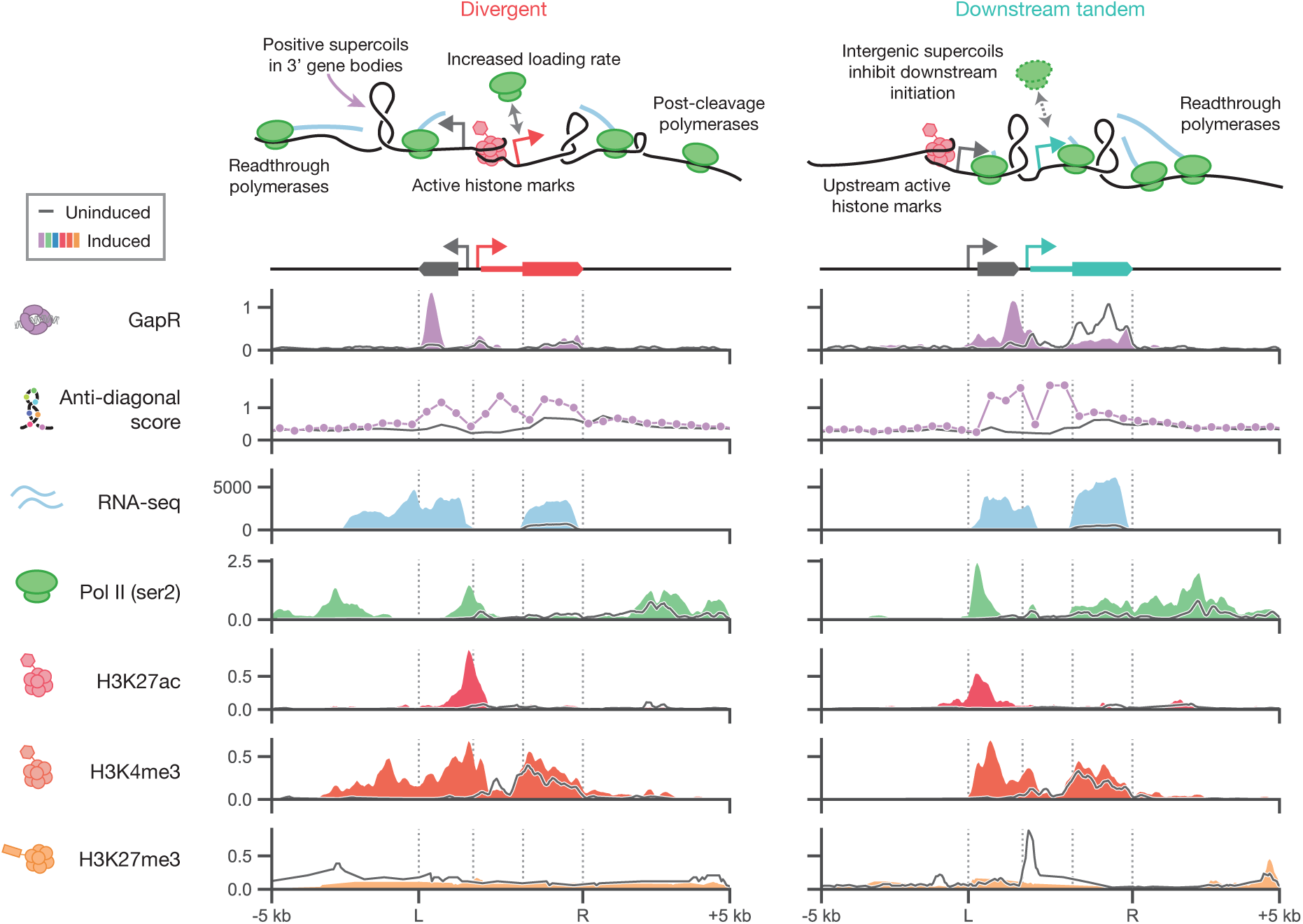
Local chromatin state shows transcription-linked supercoiling generation. For the divergent and downstream tandem two-gene syntaxes, integrated homozygously into the CLYBL locus in hiPSCs as described in fig. 3, various one dimensional genomics tracks. Tracks are shown for the integration region and the surrounding 5 kilobases to the left and right. Vertical lines show the promoter start locations, the start of the constitutive gene exon, and the polyadenylation signals. The GapR track is measured with GapRUN, a CUT&RUN derived technique, and is shown as reads-per-million. The anti-diagonal score track is computed as a one-dimensional track from the two-dimensional RCMC data and is a non-dimensional quantity that is directly proportional to contact probability. The RNA-seq track is spike-in normalized and shown as reads-per-million. The Pol II (ser2), H3K27ac, H3K4me3, and H3K27me3 tracks are measured with CUT&Tag, are spike-in normalized, and shown as reads-per-million. All tracks are shown for the uninduced (gray line) and induced (colored fills and lines) cases. Active histone marks, polymerase motion (including polymerases reading past the polyadenylation signal), and supercoiling combine to form a dynamic feedback loop that gives rise to the observed patterns of divergent amplification and tandem upstream dominance. Using the bacterial protein GapR, which binds to positive supercoiling (writhe), strong positive supercoiling is observed downstream of active transcription. Notably, positive supercoiling accumulates at either end of the divergent construct and in the intergenic region of the downstream tandem construct. These regions of positive supercoiling co-occur with strong anti-diagonal scores (RCMC) and in regions of active RNA Pol II elongation (CUT&Tag). Further comparing the positive supercoiling signal with three histone marks quantified by CUT&Tag, we observe that positive supercoiling accumulates downstream of active transcription start sites and co-occurs with active transcriptional histone marks. Upon induction, the level of the repressive mark H3K27ac decreases across the integration region.

Our direct and indirect supercoiling measurements (figs. 3e, 4 and S28) co-occur with induction-dependent changes in the transcriptional state of the edited locus. The combination of these measurements strongly suggests that the supercoiling mechanism drives syntax-specific profiles of expression.

### Syntax-based tuning optimizes circuit expression and biologic production without part substitution

Synthetic circuits and other transgenic systems such as biologic producer lines are often optimized through selection of transcription factors, promoters, stoichiometric ratios, copy number, and integration locus [63–65]. However, as each element may affect the dynamics of gene expression, forward design through simple exchange of parts remains iterative. Given that syntax modulates expression to a similar degree as genetic element selection [52], we propose using syntax to tune the relative levels of expression without changing the sequences of these elements or their relative copy number. To test this syntax-based tuning scheme, we explored optimization of two common biotechnological tools: a monoclonal antibody producer line and an inducible lentivirus system.

Low-cost production of antibodies, especially for those against infectious and tropical diseases [66], can improve worldwide access to these antibody drugs. Increasing antibody titers offers a simple way to reduce the cost of production and enhance affordable access. To demonstrate the promise of syntax-based optimization, we integrated two-gene constructs encoding the heavy and light chains of an anti-yellow-fever monoclonal antibody into a landing pad HEK293T cell line. Previous reports [65, 67–69] suggest that excess light chain translation can increase titers. Thus, based on the principle of upstream dominance, we would expect that by setting a high ratio of light chain to heavy chain, the downstream tandem syntax would outperform the upstream tandem syntax. In measuring total human IgG titer via both sandwich ELISA (fig. 5a) and a bead agglutination assay (fig. S20a), we found a nearly four-fold difference in antibody titer as a function of syntax, with the downstream tandem and divergent syntaxes providing the highest titers as expected.

**Figure 5:**
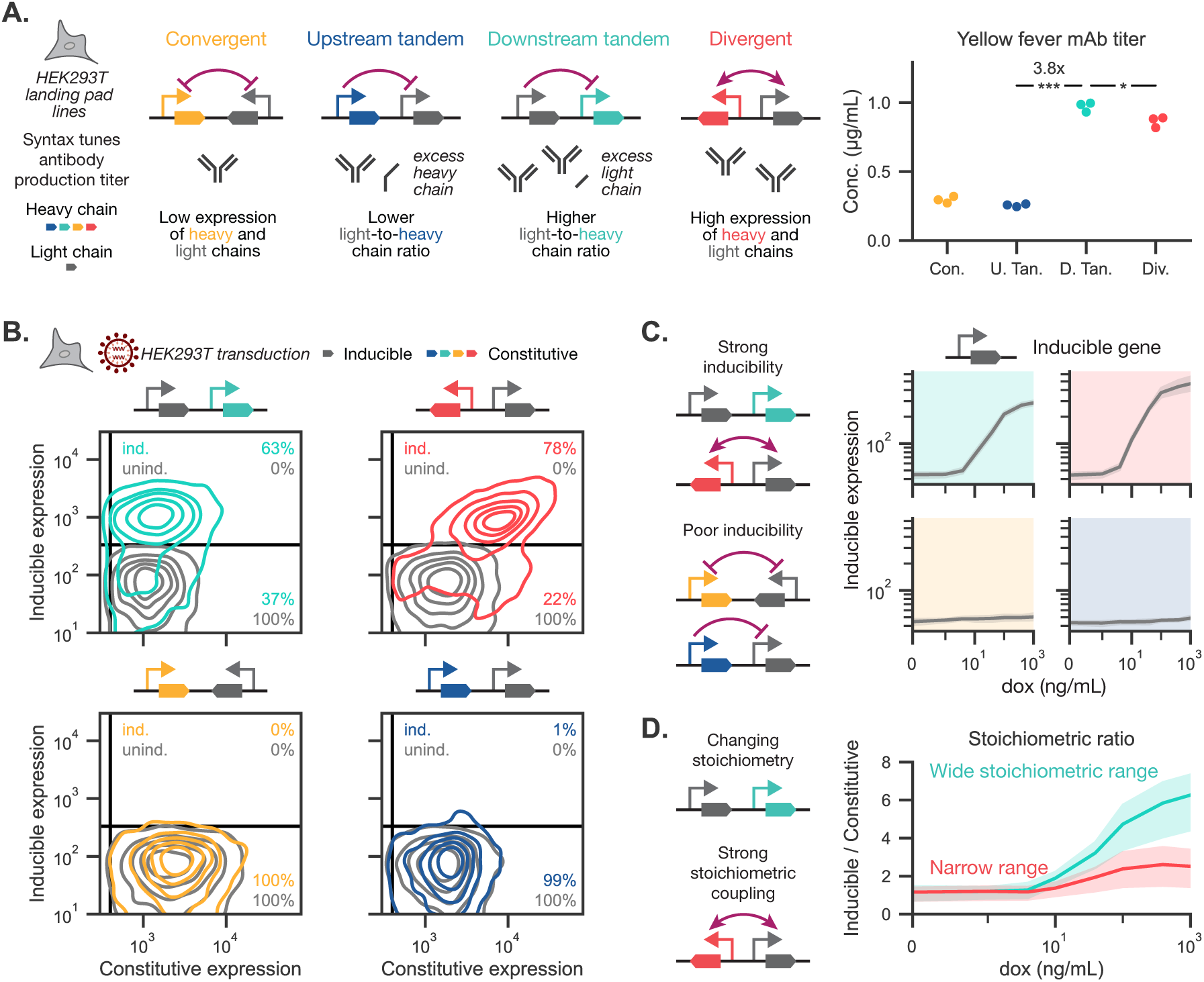
Syntax-based tuning optimizes circuit expression and biologic production without part substitution. a) The light and heavy chains of an anti-yellow-fever monoclonal antibody (mAb) are expressed from two-gene constructs (dual CMV promoters) integrated at a landing pad at the Rogi2 locus in HEK293T cells. Antibody titer, as measured via sandwich ELISA, differs across syntaxes. Points depict three biological replicates. Statistics are two-sided student t-tests. n.s.: *p >* 0.05, *: *p <* 0.05, **: *p <* 0.01, ***: *p <* 0.001, ****: *p <* 0.0001 b) Two-gene circuits, consisting of a constitutive gene (EF1α promoter) and an inducible gene (TRE promoter), were lentivirally transduced into HEK293T cells with a second vector expressing the activator rtTA (EF1α-short promoter). Joint distributions in the absence (uninduced, gray) or presence (induced, colored) of 1 µg/mL dox are shown. Percentages refer to the proportion of cells expressing (top) or not expressing (bottom) the inducible gene, as indicated by manual thresholds (black lines). c) The geometric mean of the inducible gene (gray) is shown as a function of dox concentration. Gray shading represents the 95% confidence interval across four biological replicates. d) The stoichiometric ratio between inducible and constitutive gene expression is shown as a function of dox concentration. Strong coupling of expression reduces the change in this ratio. Colored shading represents the 95% confidence interval across four biological replicates.

Lentiviruses offer efficient delivery of transgenes to diverse primary cells for therapeutic applications such as *ex vivo* engineering of CAR-T therapies and *ex vivo* immune cell reprogramming [70]. Gene circuits and inducible systems offer safe, clinically guided control and the ability to target specific cell states [35]. However, ensuring robust co-expression of multiple genes in these systems remains challenging. To explore syntax-based tuning of expression from an inducible lentivirus, we tested all four possible two-gene syntaxes transduced into HEK293T cells. Only the divergent and downstream tandem syntaxes display an appreciable double-positive population at maximum induction (figs. 5b, S21 and S22). Expression of the constitutive gene in the upstream tandem and convergent syntaxes strongly inhibits induction across all inducer concentrations (fig. 5c). In the two syntaxes with strong induction, syntax sets the stoichiometric ratio between the two genes (fig. 5d). Weak coupling between the two genes in the tandem syntax allows a wide range of stoichiometries upon induction, varying stoichiometry seven-fold (fig. 5d). Conversely, the strong positive coupling between genes in the divergent syntax maintains a narrower ratio of expression. Tuning the ratio of expression between elements can significantly shift the behavior of gene circuits, potentially supporting or impeding desired functions [71, 72]. Together, our results demonstrate that syntax can tune expression levels in diverse synthetic circuits without requiring part substitution.

### Syntax augments performance of compact gene circuits across cell types

Compact gene circuits support efficient delivery of therapeutic cargoes via size-restricted vectors such as lentiviruses and adeno-associated viruses. However, the close proximity of multiple genes in these vectors introduces the potential for physical coupling between transcriptional units. To harness supercoiling-mediated feedback for improved circuit performance, we focused on optimizing a compact, lentivirally delivered “all-in-one” inducible circuit.

Unlike the inducible circuits in figs. 2 and 5, all-in-one designs include the dox-responsive activator on the same construct, resulting in both biophysical and biochemical coupling (fig. 6a). In the divergent syntax, positive supercoiling-mediated feedback should generate high, correlated expression. For both tandem syntaxes, negative feedback should reduce the degree of correlation between genes. Despite negative feedback, we expect that the downstream tandem syntax will support induction provided that activator levels remain sufficient [55].

**Figure 6:**
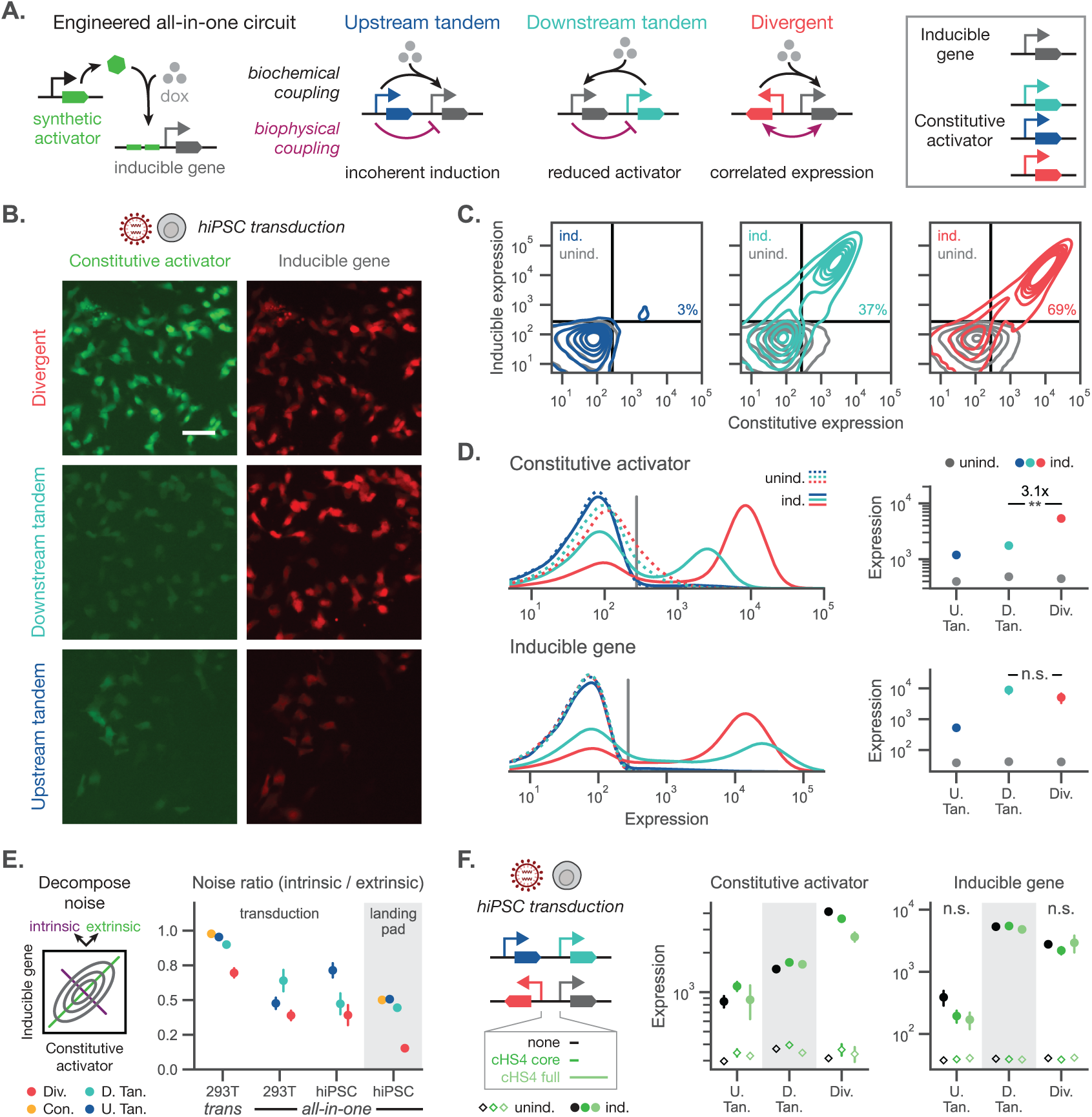
Syntax augments performance of compact gene circuits across cell types. a) A dox-inducible circuit relies on expression of a synthetic activator (rtTA) to activate the TRE promoter; an “all-in-one” circuit places both the activator—co-expressed with a fluorescent protein from the EF1α-short promoter—and the inducible gene of interest in the same cassette. The performance of the system is determined by the interplay between the biochemical coupling (black arrows) and the supercoiling-dependent biophysical coupling (purple arrows). b) Representative microscopy images show the expression of the constitutive activator (left) and the inducible gene (right) for the circuit in a) transduced into hiPSCs and induced with 300 ng/mL dox. Scale bar represents 50 microns. c) Joint distributions of activator and inducible gene expression are shown for each syntax when induced (colored) or uninduced (gray). Black lines depict expression thresholds set by the untransduced population. Percentages refer to the proportion of double-positive cells in the induced condition. d) Left: Expression distributions for the constitutive activator and inducible gene in the uninduced (dashed) and induced (solid) conditions. Vertical gray lines represent the expression thresholds from the untransduced population. Right: Geometric mean expression of the constitutive activator and inducible gene are shown for uninduced (gray) and induced (colored) populations. Fold change is annotated for the divergent syntax relative to the downstream tandem syntax when induced. e) Joint distributions can be decomposed into intrinsic (off-diagonal) and extrinsic (on-diagonal) noise. Noise is related to the variance in the log-transformed distributions of the two genes (see methods). Computed ratios for the *trans*-inducible or all-in-one circuit in HEK293Ts or hiPSCs, integrated via transduction or a landing pad. f) The cHS4 core (300 bp) or full (1300 bp) insulator sequence was placed in the intergenic region of the all-in-one circuit. The no insulator condition (none, 300 bp) is the same as in d). Geometric mean expression of each gene are shown for hiPSCs lentivirally transduced with these circuits and induced with 300 ng/mL dox (circles) or uninduced (diamonds). Points represent the mean ± standard error for three biological replicates transduced with separate batches of lentivirus. Statistics are two-sided student t-tests. n.s.: *p >* 0.05, *: *p <* 0.05, **: *p <* 0.01, ***: *p <* 0.001, ****: *p <* 0.0001

Transducing hiPSCs, we observe robust induction from the divergent and downstream tandem syntaxes (figs. 6b and 6c). These trends are mirrored for transduction in HEK293T cells and mouse embryonic fibroblasts (fig. S23). Expression of the synthetic activator in the downstream tandem syntax is more than an order of magnitude lower than in the divergent syntax. However, for all syntaxes, expression of the activator increases relative to the uninduced case (fig. 6d). This increase may reflect a local increase in transcriptional resources that affects circuits with the activator in *cis* but not those in *trans* (i.e., with a separately integrated activator, as in figs. 2 and 5) (fig. S22) [55].

Supercoiling-mediated feedback is predicted to couple the probabilities of transcriptional bursting [7]. Even for stable protein reporters, changes in correlated bursting may be visible in the variance of co-expression. Variance between two genes across a population of cells can be decomposed into two components: intrinsic noise, which quantifies the variability within individual single cells, and extrinsic noise, which reflects differences such as cell size [73, 74]. Conceptually, this decomposition of the variance is related to the “width” of the distribution along the main diagonal (extrinsic) and off-diagonal (intrinsic) when relating expression of the two genes. Dividing the intrinsic noise by the extrinsic noise gives a noise ratio that represents the relative contribution of intrinsic noise. Computing this metric, we observe that the divergent syntax consistently generates the lowest noise ratio (fig. 6e); this generalizes across integration method, cell type, and circuit type (all-in-one and *trans*-located activator). These observations align with our previous modeling predictions, where transcriptional co-bursting in the divergent syntax dramatically reduces intrinsic noise relative to extrinsic noise (fig. S24a) [7]. In all cases, we observe that syntax affects both the intrinsic and extrinsic noise (figs. S24b to S24f). Finally, placing the synthetic activator in *trans* to the main circuit increases the noise ratio relative to the all-in-one circuits (fig. 6e). However, the *trans* circuits have lower extrinsic noise (fig. S24d), suggesting that placing the synthetic activator distal to the two-gene circuit dramatically increases the intrinsic noise.

Insulator sequences are used in synthetic circuits to reduce coupling between integrated transgenes [75–77], potentially mitigating the effects of supercoiling-mediated feedback. To test this hypothesis, we added cHS4 insulator sequences to the intergenic region in the all-in-one circuit. For all syntaxes, addition of the cHS4 core or full insulator sequence did not substantially change expression levels or noise profiles (figs. 6f and S25). Similarly, non-insulator spacer sequences ranging from 300 bp to 1.8 kb did not change the observed syntax-dependent trends (fig. S26).

In contrast to intergene spacer length, both gene length and average transcription rate are expected to influence the amount of generated supercoiling according to prior modeling. Longer genes generally generate more supercoiling for the same transcription rate [7]. Integrating two-gene circuits into hiPSCs at the *CLYBL* locus with differing inducible gene lengths (fig. S27), we observe that protein expression of the inducible gene decreases monotonically with length. The expression of the constitutive gene remains largely the same as a function of inducible gene length, potentially indicating that the enhanced supercoiling from longer genes compensates for the reduction in expression. Together, these data indicate that syntax offers a powerful design parameter for coupling and tuning profiles of expression that can be harnessed to dampen or amplify noise [78, 79].

## Discussion

Transcription forms a dynamic feedback loop mediated by DNA supercoiling. Here we use two-gene synthetic gene circuits as model systems to probe the biophysical influence of adjacent transcription (fig. 1). Using inducible synthetic circuits, we isolate the effects of reversible biophysical coupling and examine the impact of transcriptionally induced coupling on gene expression (fig. 2) and chromatin structure. Applying fine-scale mapping of a genomic safe harbor using RCMC, we observe both transcriptionally induced anti-diagonal stripes at the 5–10-kb length scale and polymerase-dependent perturbations of ∼200-kb TADs and associated loop domains (figs. 3 and S13). Importantly, we attribute the strong anti-diagonal stripes to the formation of transcriptionally induced plectonemes, as these stripes are colocalized with positive supercoiling as measured by GapR, with actively elongating RNA Pol II, and with histone marks characteristic of active transcription. Integrating the predictions of supercoiling-mediated feedback into circuit design, we use syntax-based optimization to increase antibody production, to tune expression from lentiviruses, and to explore performance regimes of inducible circuits in diverse cell types (fig. 5). We find that, in agreement with our model of supercoiling-dependent biophysical feedback [7], syntax-based optimization is mostly unaffected by intergene spacers (figs. 6f, S3, S25 and S26). Overall, we find that syntax defines the profiles and performance of gene circuits across a range of integration methods, cargoes, and cell types, offering a design parameter for tuning the performance and predictability of circuits (fig. 6).

Syntax offers an orthogonal design variable that can be combined with library-based approaches. Growing compendiums of parts expand the potential for library-based approaches for circuit tuning [64, 80, 81], but syntax-based tuning can be used to optimize circuits even when parts are constrained [55]. When changing design features including promoters, polyadenylation sequences, cell type, integration locus, and integration method, we consistently observed reproducible syntax-specific coupling of gene expression. Syntax-specific profiles persist even in the presence of putative insulator sequences commonly used in synthetic biology such cHS4 and CTCF-binding motifs [47–49], suggesting these features do not serve as barriers to supercoiling-induced coupling. However, exploration of additional intergene sequences, genetic elements, and locations within the genome may reveal elements that mitigate or amplify syntax-specific coupling.

In both constitutive and inducible systems, we find that upstream dominance strongly affects closely spaced tandem genes, resulting in reduced expression of the downstream gene (fig. 1). Previous work suggests that upstream dominance affects integrated transgenes [82], that choice of gene orientation and direction affects expression from adenovirus vectors [83], and that the divergent syntax offers an efficient “all-in-one” Cas9 editing circuit [84]. Increasing numbers of synthetic circuits employ divergent syntax for delivery to primary cells [35, 36, 85, 86]. As these applications require high rates of co-delivery, choice of divergent syntax may reflect the selection of functional circuits during the design process.

Supercoiling-mediated feedback provides an extremely rapid mechanism of transcriptional coupling [7]. In alignment with our predictions of supercoiling-mediated feedback, we find that transcription of an adjacent gene induces reversible, syntax-specific profiles of expression, tuning both the mean and variance (figs. 2, 6e and S24). Remarkably, the effect of syntax on the ratio of intrinsic and extrinsic noise generalizes over both integration method and cell type. Parallel work in yeast showed increased burst coupling for *cis* divergent genes compared to those in *trans* at the *Gal1-Gal10* locus [11]. Potentially, harnessing the fast-timescale feedback of supercoiling may improve the performance of dynamic circuits such a pulse generators, toggle switches, and oscillators, which require coordinated expression of multiple genes. Supercoiling-mediated feedback may be especially useful to buffer noise in RNA-based control systems, supporting perfect adaptation and dosage control [36, 85, 87]. As noise and small changes in expression can direct cell fate [44, 78, 88, 89], syntax-based tuning offers a simple method to explore the stability of cell fate by perturbing native networks with levels of transgenes that vary in their mean and variability.

Our findings are consistent with both biophysical predictions of DNA-supercoiling-mediated feedback [7] and *in vitro* studies [26, 90–92], which used GapR to measure positive supercoiling [17, 34] or small-molecule intercalators like psoralen to measure negative supercoiling [22, 93–96]. Our work links strong anti-diagonal scores measured with RCMC—an indirect measurement of negative or positive supercoiling—with the presence of positive supercoiling measured directly with GapR. In our work, we measure how induction of transcriptional activity at a single locus induces perturbations in supercoiling and chromatin state at that engineered locus. Future work could use library screening approaches to observe the effects of targeted perturbations at loci across the genome to fully explore the hypothesis of DNA supercoiling-mediated feedback.

Engineered synthetic circuits are much more compact than native mammalian genomes and therefore are putatively more strongly affected by supercoiling. However, regulation of some native genes depend on adjacent gene expression. Tandem arrays of *Hox* genes show temporal activation during development that proceeds from downstream to upstream genes [97], a process named “posterior dominance”. Potentially, supercoiling-mediated upstream dominance may stabilize this phenomenon, which restricts the reactivation of downstream *Hox* genes. Transcription of non-coding RNAs tunes expression of native adjacent genes, both amplifying and attenuating expression of co-localized genes [44, 98]. Loss of transcription from an adjacent gene can induce significant developmental defects, suggesting adjacent transcription plays an essential role in tuning expression to control and coordinate cell fate [3, 44].

While supercoiling-mediated feedback offers a compelling explanation for the coupling of gene expression and for syntax-specific profiles of intrinsic noise, there remain limits to the model of supercoiling-mediated feedback that may be explained by alternative modes of regulation. For instance, collisions of RNA polymerases undergoing transcriptional readthrough may reduce expression in the convergent and tandem syntaxes [99]. We also observe non-monotonic changes in reporter expression with increasing adjacent gene length, a variable predicted to chiefly increase the amount of generated supercoiling (fig. S27). However, these predictions assume the basal rate of transcription remains constant while gene length alone varies. It remains challenging to develop a matched experimental system where gene length can be changed without modifying transcription rates, mRNA stability, or translation efficiency. Modifying circuit sequences can lead to RNA species with dramatically different processing rates [52]. The local transcriptional neighborhood is also known to affect transcript isoforms, stability, and overall expression levels [100]. Syntax-mediated changes in RNA isoforms, modifications, and processing may explain the differences in the levels of RNA and protein observed in the downstream tandem syntax in hiPSCs that were not observed in HEK293Ts. Overall, we have shown that syntax-specific behaviors generalize across hiPSCs, MEFs, and HEK293T cells. However, we expect future work to uncover how cell-type– and species–dependent changes in topoisomerase levels, transcriptional resources, and other biochemical factors will affect the level of biophysical coupling. For instance, the field lacks a definitive model of how topoisomerase recruitment occurs and how topoisomerases differentially target positive and negative supercoiling in human cells. Existing studies from other organisms suggest that supercoiling perturbations non-monotonically affect transcription rates [101, 102], making it difficult to predict syntax-specific effects of topoisomerase inhibition. While genome-wide perturbations of topoisomerase activity offer some mechanistic insight [22], synthetic circuits integrated at defined loci in single copy could precisely measure how targeted recruitment of topoisomerases or other locus-specific perturbations interact with biophysical coupling. Alternatively, methods of defining polymerase positions on single DNA fibers may resolve questions on readthrough and facilitated recruitment of RNA Pol II via supercoiling-mediated feedback [103].

Despite the clean, abstract way that synthetic circuits are often drawn, integration into the genome wraps synthetic circuitry in layers of native regulation. Biochemical interactions *and* biophysical forces combine to shape our genomes. By harnessing both layers of control, leveraging gene syntax and supercoiling-mediated feedback can enhance the predictability, performance, and functional range of engineered gene circuits.

## Acknowledgments

We thank Matheus Oliveira de Souza for providing the antibody-encoding plasmids used for the antibody work. We thank the MIT BioMicro Center (supported in part by the Koch Institute Support (core) Grant P30-CA14051 from the National Cancer Institute and by the National Institute of Environmental Health Sciences of the NIH under award P30-ES002109) for their support with the sequencing-based assays in this study. We thank Mike Laub and Anders Hansen for their feedback. We thank Mary Ehmann, Maria Castellanos, and Nat Wang for reviewing the manuscript. Research reported in this manuscript was supported by the NIH NIGMS under award number R35-GM143033, by the National Science Foundation under the NSF-CAREER under award number 2339986, and with funding from Institute for Collaborative Biotechnologies (W911NF-19-2-0026), and by the Air Force Research Laboratory MURI (FA9550-22-1-03t16). K.S.L. is supported by the National Science Foundation Graduate Research Fellowship Program under grant No. 1745302. R.D.J. is supported by a Michael Smith Health Research BC trainee award (RT-2021-1946) and an MSL Pathway to Independence Award. Work in P.W.Z.’s lab was supported by the Stem Cell Network, the Canadian Institute for Health Research, and the Wellcome Leap Human Organs, Physiology, and Engineering (HOPE) program. P.W.Z. is the Canada Research Chair in Stem Cell Bioengineering. C.L.M., R.D. and A.B.A. were supported by Novo Nordisk Foundation grant (NNF21CC0073729; reNEW). B.J.D. is supported by the NIH (R01AI181684 and R21AI166396).

## Author contributions

Following the contributor role taxonomy, C.P.J. contributed to project conceptualization, methodology, validation, investigation/data acquisition, statistical analysis, writing (original draft, review, editing), and visualization. K.S.L. contributed to conceptualization and methodology (all-in-one circuits), validation, investigation/data acquisition, statistical analysis, writing (review, editing), and visualization. S.R.K. contributed to investigation/data acquisition (two-gene lentiviruses) and writing (review, editing). R.D.J. contributed to conceptualization and methodology (constitutive hiPSC circuits), data acquisition, and visualization.

A.B.A. contributed to conceptualization and methodology (STRAIGHT-IN circuits), investigation/data acquisition, and writing (review). D.S.P. contributed to conceptualization and methodology (STRAIGHT-IN circuits), investigation/data acquisition, and writing (review). E.L.P contributed to investigation/data acquisition (RNA-HCR-FISH) and writing (review, editing). R.L. contributed to investigation/data acquisition (constitutive PiggyBac cell lines) and writing (review). J.Y. contributed to investigation/data acquisition (constitutive hiPSC circuits). C.G.O. contributed to methodology (two-gene PiggyBac constructs) and writing (review). C.L.M. contributed to resources, writing (review), supervision, amd funding acquisition. R.P.D. contributed to resources, writing (review), supervision, and funding acquisition. B.J.D. conceptualization and methodology (antibody experiments), resources, and writing (review). P.W.Z. contributed to writing (review), resources, supervision, project administration, and funding acquisition. K.E.G. contributed to conceptualization, methodology, validation, resources, writing (review, editing), supervision, project administration, and funding acquisition.

## Supplementary Materials

- Materials and Methods
- Tables S1 to S5.
- Figs. S1 to S29.

## References

1. Darbellay, F. et al. The Constrained Architecture of Mammalian Hox Gene Clusters. PNAS 116, 13424–13433. issn: 0027-8424, 1091-6490. PMID: 31209053 (July 2, 2019) (cit. on p. 1).

2. Murphy, S. E. & Boettiger, A. N. Polycomb Repression of Hox Genes Involves Spatial Feedback but Not Domain Compaction or Phase Transition. Nat Genet 56, 493–504. issn: 1546-1718 (Mar. 2024) (cit. on p. 1).

3. Zinani, O. Q. H., Keseroğlu, K., Ay, A. & Özbudak, E. M. Pairing of Segmentation Clock Genes Drives Robust Pattern Formation. Nature, 1–6. issn: 1476-4687 (Dec. 23, 2020) (cit. on pp. 1, 15).

4. Gherman, A., Wang, R. & Avramopoulos, D. Orientation, Distance, Regulation and Function of Neighbouring Genes. Hum Genomics 3, 143–156. issn: 1473-9542. PMID: 19164091 (Jan. 1, 2009) (cit. on p. 1).

5. Trinklein, N. D. et al. An Abundance of Bidirectional Promoters in the Human Genome. Genome Res. 14, 62–66. issn: 1088-9051, 1549-5469. PMID: 14707170 (Jan. 1, 2004) (cit. on p. 1).

6. Korbel, J. O., Jensen, L. J., von Mering, C. & Bork, P. Analysis of Genomic Context: Prediction of Functional Associations from Conserved Bidirectionally Transcribed Gene Pairs. Nat Biotechnol 22, 911–917. issn: 1546-1696 (July 2004) (cit. on p. 1).

7. Johnstone, C. P. & Galloway, K. E. Supercoiling-Mediated Feedback Rapidly Couples and Tunes Transcription. Cell Reports 41. issn: 2211-1247 (Oct. 18, 2022) (cit. on pp. 1–4, 12, 14, 53).

8. Brahmachari, S., Tripathi, S., Onuchic, J. N. & Levine, H. Nucleosomes Play a Dual Role in Regulating Transcription Dynamics. Proceedings of the National Academy of Sciences 121, e2319772121 (July 9, 2024) (cit. on pp. 1, 2).

9. Tsao, Y.-P., Wu, H.-Y. & Liu, L. F. Transcription-Driven Supercoiling of DNA: Direct Biochemical Evidence from in Vitro Studies. Cell 56, 111–118. issn: 0092-8674 (Jan. 13, 1989) (cit. on p. 2).

10. Janissen, R., Barth, R., Polinder, M., van der Torre, J. & Dekker, C. Single-Molecule Visualization of Twin-Supercoiled Domains Generated during Transcription. Nucleic Acids Research 52, 1677–1687. issn: 0305-1048 (Feb. 28, 2024) (cit. on p. 2).

11. Patel, H. P. et al. DNA Supercoiling Restricts the Transcriptional Bursting of Neighboring Eukaryotic Genes. Molecular Cell 83, 1573–1587.e8. issn: 1097-2765. PMID: 37207624 (May 18, 2023) (cit. on pp. 2, 14).

12. Ancona, M., Bentivoglio, A., Brackley, C. A., Gonnella, G. & Marenduzzo, D. Transcriptional Bursts in a Nonequilibrium Model for Gene Regulation by Supercoiling. Biophysical Journal 117, 369–376. issn: 00063495 (July 2019) (cit. on p. 2).

13. Geng, Y. et al. A Spatially Resolved Stochastic Model Reveals the Role of Supercoiling in Transcription Regulation. PLOS Computational Biology 18, e1009788. issn: 1553-7358 (Sept. 19, 2022) (cit. on p. 2).

14. Ma, J. et al. Transcription Factor Regulation of RNA Polymerase’s Torque Generation Capacity. Proceedings of the National Academy of Sciences 116, 2583–2588 (Feb. 12, 2019) (cit. on p. 2).

15. Qian, J. et al. Chromatin Buffers Torsional Stress During Transcription. bioRxiv, 2024.10.15.618270. issn: 2692-8205. PMID: 39464147 (Oct. 18, 2024) (cit. on p. 2).

16. Bhola, M. et al. RNA Interacts with Topoisomerase I to Adjust DNA Topology. Molecular Cell 84, 3192–3208.e11. issn: 1097-2765. PMID: 39173639 (Sept. 5, 2024) (cit. on p. 2).

17. Longo, G. M. C. et al. Type II Topoisomerases Shape Multi-Scale 3D Chromatin Folding in Regions of Positive Supercoils. Molecular Cell. issn: 1097-2765 (Oct. 31, 2024) (cit. on pp. 2, 8, 14, 28, 30).

18. Lee, J. et al. Chromatinization Modulates Topoisomerase II Processivity https://www.biorxiv.org/content/10.1101/2023.10.03.560726v1 (2023). Pre-published (cit. on p. 2).

19. Saha, S. & Pommier, Y. R-Loops, Type I Topoisomerases and Cancer. NAR Cancer 5, zcad013. issn: 2632-8674 (Mar. 1, 2023) (cit. on p. 2).

20. Sutormin, D. et al. Interaction between Transcribing RNA Polymerase and Topoisomerase I Prevents R-loop Formation in E. Coli. Nat Commun 13, 4524. issn: 2041-1723. PMID: 35927234 (Aug. 4, 2022) (cit. on p. 2).

21. Le, T. T. et al. Synergistic Coordination of Chromatin Torsional Mechanics and Topoisomerase Activity. Cell 179, 619–631.e15. issn: 0092-8674 (Oct. 17, 2019) (cit. on p. 2).

22. Yao, Q., Zhu, L., Shi, Z., Banerjee, S. & Chen, C. Topoisomerase-Modulated Genome-Wide DNA Supercoiling Domains Colocalize with Nuclear Compartments and Regulate Human Gene Expression. Nat Struct Mol Biol, 1–14. issn: 1545-9985 (Aug. 16, 2024) (cit. on pp. 2, 14, 15).

23. Corless, S. & Gilbert, N. Effects of DNA Supercoiling on Chromatin Architecture. Biophys Rev 8, 245–258. issn: 1867-2469 (Sept. 1, 2016) (cit. on p. 2).

24. Achar, Y. J., Adhil, M., Choudhary, R., Gilbert, N. & Foiani, M. Negative Supercoil at Gene Boundaries Modulates Gene Topology. Nature 577, 701–705. issn: 1476-4687 (7792 Jan. 2020) (cit. on p. 2).

25. Naughton, C. et al. Transcription Forms and Remodels Supercoiling Domains Unfolding Large-Scale Chromatin Structures. Nat Struct Mol Biol 20, 387–395. issn: 1545-9985. PMID: 23416946 (3 Mar. 2013) (cit. on p. 2).

26. Revyakin, A., Ebright, R. H. & Strick, T. R. Promoter Unwinding and Promoter Clearance by RNA Polymerase: Detection by Single-Molecule DNA Nanomanipulation. PNAS 101, 4776–4780. issn: 0027-8424, 1091-6490. PMID: 15037753 (Apr. 6, 2004) (cit. on pp. 2, 14).

27. Boulas, I. et al. Assessing in Vivo the Impact of Gene Context on Transcription through DNA Super-coiling. Nucleic Acids Research 51, 9509–9521. issn: 0305-1048 (Oct. 13, 2023) (cit. on p. 2).

28. Marko, J. F. Torque and Dynamics of Linking Number Relaxation in Stretched Supercoiled DNA. *Phys*. Rev. E 76, 021926. issn: 1539-3755, 1550-2376 (Aug. 29, 2007) (cit. on p. 2).

29. Sevier, S. A. & Levine, H. Mechanical Properties of Transcription. Phys. Rev. Lett. 118, 268101 (June 27, 2017) (cit. on p. 2).

30. Hacker, W. C. & Elcock, A. H. Spotter: A Single-Nucleotide Resolution Stochastic Simulation Model of Supercoiling-Mediated Transcription and Translation in Prokaryotes. *Nucleic Acids Research*, gkad682. issn: 0305-1048 (Aug. 21, 2023) (cit. on p. 2).

31. Chong, S., Chen, C., Ge, H. & Xie, X. S. Mechanism of Transcriptional Bursting in Bacteria. Cell 158, 314–326. issn: 0092-8674 (July 17, 2014) (cit. on p. 2).

32. Yeung, E. et al. Biophysical Constraints Arising from Compositional Context in Synthetic Gene Networks. Cell Systems 5, 11–24.e12. issn: 2405-4712 (July 26, 2017) (cit. on p. 2).

33. Teves, S. S. & Henikoff, S. Transcription-Generated Torsional Stress Destabilizes Nucleosomes. Nat Struct Mol Biol 21, 88–94. issn: 1545-9993. PMID: 24317489 (1 Jan. 2014) (cit. on p. 2).

34. Guo, M. S., Kawamura, R., Littlehale, M. L., Marko, J. F. & Laub, M. T. High-Resolution, Genome-Wide Mapping of Positive Supercoiling in Chromosomes. eLife 10 (eds Berger, J. M. & Barkai, N.) e67236. issn: 2050-084X (July 19, 2021) (cit. on pp. 2, 8, 14).

35. Li, H.-S. et al. Multidimensional Control of Therapeutic Human Cell Function with Synthetic Gene Circuits. Science 378, 1227–1234 (Dec. 16, 2022) (cit. on pp. 2, 10, 14).

36. Love, K. S., Johnstone, C. P., Peterman, E. L., Gaglione, S. & Galloway, K. E. Model-Guided Design of microRNA-based Gene Circuits Supports Precise Dosage of Transgenic Cargoes into Diverse Primary Cells https://www.biorxiv.org/content/10.1101/2024.06.25.600629v2 (2024). Pre-published (cit. on pp. 2, 14).

37. Cho, J. H. et al. Engineering Advanced Logic and Distributed Computing in Human CAR Immune Cells. Nat Commun 12, 792. issn: 2041-1723 (Feb. 4, 2021) (cit. on p. 2).

38. Anzalone, A. V. et al. Search-and-Replace Genome Editing without Double-Strand Breaks or Donor DNA. Nature 576, 149–157. issn: 1476-4687 (Dec. 2019) (cit. on p. 2).

39. Mali, P. et al. RNA-Guided Human Genome Engineering via Cas9. Science 339, 823–826 (Feb. 15, 2013) (cit. on p. 2).

40. Cong, L. et al. Multiplex Genome Engineering Using CRISPR/Cas Systems. Science 339, 819–823 (Feb. 15, 2013) (cit. on p. 2).

41. Tousley, A. M. et al. Co-Opting Signalling Molecules Enables Logic-Gated Control of CAR T Cells. Nature 615, 507–516. issn: 1476-4687 (Mar. 2023) (cit. on p. 2).

42. Allen, G. M. et al. Synthetic Cytokine Circuits That Drive T Cells into Immune-Excluded Tumors. Science 378, eaba1624 (Dec. 16, 2022) (cit. on p. 2).

43. Gao, W. et al. Engineered T Cell Therapy for Central Nervous System Injury. Nature 634, 693–701. issn: 1476-4687 (Oct. 2024) (cit. on p. 2).

44. Ganesh, V. S., et al. Novel Syndromic Neurodevelopmental Disorder Caused by de Novo Deletion of CHASERR, a Long Noncoding RNA https://www.medrxiv.org/content/10.1101/2024.01. 31.24301497v1 (2024). Pre-published (cit. on pp. 3, 14, 15).

45. Rom, A. et al. Regulation of CHD2 Expression by the Chaserr Long Noncoding RNA Gene Is Essential for Viability. Nat Commun 10, 5092. issn: 2041-1723 (Nov. 8, 2019) (cit. on p. 3).

46. Martens, J. A., Laprade, L. & Winston, F. Intergenic Transcription Is Required to Repress the Saccharomyces Cerevisiae SER3 Gene. Nature 429, 571–574. issn: 1476-4687 (June 2004) (cit. on p. 3).

47. Tang, Z. et al. CTCF-Mediated Human 3D Genome Architecture Reveals Chromatin Topology for Transcription. Cell 163, 1611–1627. issn: 0092-8674, 1097-4172. PMID: 26686651 (Dec. 17, 2015) (cit. on pp. 3, 14).

48. Liu, M. et al. Genomic Discovery of Potent Chromatin Insulators for Human Gene Therapy. Nat Biotechnol 33, 198–203. issn: 1546-1696 (Feb. 2015) (cit. on pp. 3, 4, 14).

49. Lu, X.-B., Guo, Y.-H. & Huang, W. Characterization of the cHS4 Insulator in Mouse Embryonic Stem Cells. FEBS Open Bio 10, 644–656. issn: 2211-5463. PMID: 32087050 (Apr. 2020) (cit. on pp. 3, 14).

50. Zhang, S., Übelmesser, N., Barbieri, M. & Papantonis, A. Enhancer–Promoter Contact Formation Requires RNAPII and Antagonizes Loop Extrusion. Nat Genet, 1–9. issn: 1546-1718 (Apr. 3, 2023) (cit. on p. 4).

51. Calvo-Roitberg, E. et al. mRNA Initiation and Termination Are Spatially Coordinated https://www.biorxiv.org/content/10.1101/2024.01.05.574404v1 (2024). Pre-published (cit. on p. 4).

52. Peterman, E. L., et al. High-Resolution Profiling Reveals Coupled Transcriptional and Translational Regulation of Transgenes https://www.biorxiv.org/content/10.1101/2024.11.26.625483v2 (2024). Pre-published (cit. on pp. 4, 10, 15, 25, 32).

53. Choi, H. M. T., Beck, V. A. & Pierce, N. A. Next-Generation in Situ Hybridization Chain Reaction: Higher Gain, Lower Cost, Greater Durability. ACS Nano 8, 4284–4294. issn: 1936-0851 (May 27, 2014) (cit. on p. 4).

54. Björkegren, C. & Baranello, L. DNA Supercoiling, Topoisomerases, and Cohesin: Partners in Regulating Chromatin Architecture? International Journal of Molecular Sciences 19, 884. issn: 1422-0067 (3 Mar. 2018) (cit. on p. 6).

55. Blanch-Asensio, A., et al. STRAIGHT-IN Dual: A Platform for Dual, Single-Copy Integrations of DNA Payloads and Gene Circuits into Human Induced Pluripotent Stem Cells https://www.biorxiv.org/content/10.1101/2024.10.17.616637v1 (2024). Pre-published (cit. on pp. 6, 12, 14, 26, 32).

56. Cerbini, T. et al. Transcription Activator-like Effector Nuclease (TALEN)-Mediated CLYBL Targeting Enables Enhanced Transgene Expression and One-Step Generation of Dual Reporter Human Induced Pluripotent Stem Cell (iPSC) and Neural Stem Cell (NSC) Lines. PLoS ONE 10, e0116032. issn: 1932-6203. PMID: 25587899 (2015) (cit. on p. 6).

57. Goel, V. Y., Huseyin, M. K. & Hansen, A. S. Region Capture Micro-C Reveals Coalescence of Enhancers and Promoters into Nested Microcompartments. Nat Genet 55, 1048–1056. issn: 1546-1718 (6 June 2023) (cit. on pp. 6, 27, 30).

58. Open2C et al. Cooltools: Enabling High-Resolution Hi-C Analysis in Python. PLOS Computational Biology 20, e1012067. issn: 1553-7358 (May 6, 2024) (cit. on p. 6).

59. Zufferey, M., Tavernari, D., Oricchio, E. & Ciriello, G. Comparison of Computational Methods for the Identification of Topologically Associating Domains. Genome Biology 19, 217. issn: 1474-760X (Dec. 10, 2018) (cit. on p. 6).

60. Narducci, D. N. & Hansen, A. S. Reeling It in: How DNA Topology Drives Loop Extrusion by Condensin. Nat Struct Mol Biol 29, 623–625. issn: 1545-9985 (July 2022) (cit. on p. 6).

61. Guo, Y. et al. Chromatin Jets Define the Properties of Cohesin-Driven in Vivo Loop Extrusion. Molecular Cell 82, 3769–3780.e5. issn: 1097-2765. PMID: 36182691 (Oct. 20, 2022) (cit. on p. 6).

62. Fu, Z., Guo, M. S., Zhou, W. & Xiao, J. Differential Roles of Positive and Negative Supercoiling in Organizing the E. Coli Genome. Nucleic Acids Research 52, 724–737. issn: 0305-1048 (Jan. 25, 2024) (cit. on p. 8).

63. Gam, J. J., DiAndreth, B., Jones, R. D., Huh, J. & Weiss, R. A ‘Poly-Transfection’ Method for Rapid, One-Pot Characterization and Optimization of Genetic Systems. Nucleic Acids Res 47, e106. issn: 0305-1048. PMID: 31372658 (Oct. 10, 2019) (cit. on p. 10).

64. O’Connell, R. W., et al. Ultra-High Throughput Mapping of Genetic Design Space https://www.biorxiv.org/content/10.1101/2023.03.16.532704v1 (2023). Pre-published (cit. on pp. 10, 14).

65. Schlatter, S. et al. On the Optimal Ratio of Heavy to Light Chain Genes for Efficient Recombinant Antibody Production by CHO Cells. Biotechnology Progress 21, 122–133. issn: 1520-6033 (2005) (cit. on p. 10).

66. Hooft van Huijsduijnen, R., et al. Reassessing Therapeutic Antibodies for Neglected and Tropical Diseases. PLoS Negl Trop Dis 14, e0007860. issn: 1935-2727. PMID: 31999695 (Jan. 30, 2020) (cit. on p. 10).

67. Carver, J. et al. Maximizing Antibody Production in a Targeted Integration Host by Optimization of Subunit Gene Dosage and Position. Biotechnology Progress 36, e2967. issn: 1520-6033 (2020) (cit. on p. 10).

68. Lee, Z., Wan, J., Shen, A. & Barnard, G. Gene Copy Number, Gene Configuration and LC/HC mRNA Ratio Impact on Antibody Productivity and Product Quality in Targeted Integration CHO Cell Lines. Biotechnology Progress 40, e3433. issn: 1520-6033 (2024) (cit. on p. 10).

69. Ho, S. C. L. et al. Control of IgG LC:HC Ratio in Stably Transfected CHO Cells and Study of the Impact on Expression, Aggregation, Glycosylation and Conformational Stability. Journal of Biotechnology 165, 157–166. issn: 0168-1656 (June 10, 2013) (cit. on p. 10).

70. Ascic, E. et al. In Vivo Dendritic Cell Reprogramming for Cancer Immunotherapy. Science 386, eadn9083 (Sept. 5, 2024) (cit. on p. 10).

71. Chen, Z. et al. A Synthetic Protein-Level Neural Network in Mammalian Cells. Science 386, 1243–1250 (Dec. 13, 2024) (cit. on p. 10).

72. Galloway, K. & Johnstone, C. Bringing Neural Networks to Life. Science 386, 1225–1226 (Dec. 13, 2024) (cit. on p. 10).

73. Elowitz, M. B., Levine, A. J., Siggia, E. D. & Swain, P. S. Stochastic Gene Expression in a Single Cell. Science 297, 1183–1186. issn: 0036-8075, 1095-9203. PMID: 12183631 (Aug. 16, 2002) (cit. on p. 12).

74. Swaffer, M. P. et al. RNA Polymerase II Dynamics and mRNA Stability Feedback Scale mRNA Amounts with Cell Size. Cell. issn: 0092-8674 (Nov. 8, 2023) (cit. on p. 12).

75. Cabrera, A. et al. The Sound of Silence: Transgene Silencing in Mammalian Cell Engineering. Cell Systems 13, 950–973. issn: 2405-4712 (Dec. 21, 2022) (cit. on p. 12).

76. Yahata, K. et al. cHS4 Insulator-mediated Alleviation of Promoter Interference during Cell-based Expression of Tandemly Associated Transgenes. Journal of Molecular Biology 374, 580–590. issn: 0022-2836 (Nov. 30, 2007) (cit. on p. 12).

77. Guye, P., Li, Y., Wroblewska, L., Duportet, X. & Weiss, R. Rapid, Modular and Reliable Construction of Complex Mammalian Gene Circuits. Nucleic Acids Res 41, e156. issn: 0305-1048 (Sept. 1, 2013) (cit. on p. 12).

78. Desai, R. V. et al. A DNA-repair Pathway Can Affect Transcriptional Noise to Promote Cell Fate Transitions. Science. issn: 0036-8075, 1095-9203. PMID: 34301855 (July 22, 2021) (cit. on pp. 12, 14).

79. Johnstone, C. P. & Galloway, K. E. Engineering Cellular Symphonies out of Transcriptional Noise. Nature Reviews Molecular Cell Biology. issn: 1471-0080 (Mar. 15, 2021) (cit. on p. 12).

80. Tycko, J. et al. High-Throughput Discovery and Characterization of Human Transcriptional Effectors. Cell 183, 2020–2035.e16. issn: 0092-8674 (Dec. 23, 2020) (cit. on p. 14).

81. Oesinghaus, L., Castillo-Hair, S., Ludwig, N., Keller, A. & Seelig, G. Quantitative Design of Cell Type-Specific mRNA Stability from microRNA Expression Data https://www.biorxiv.org/content/10.1101/2024.10.28.620728v1 (2025). Pre-published (cit. on p. 14).

82. Beal, K. M. et al. The Impact of Expression Vector Position on Transgene Transcription Allows for Rational Expression Vector Design in a Targeted Integration System. Biotechnology Journal 18, 2300038. issn: 1860-7314 (2023) (cit. on p. 14).

83. Fry, L. E. et al. Promoter Orientation within an AAV-CRISPR Vector Affects Cas9 Expression and Gene Editing Efficiency. CRISPR J 3, 276–283. issn: 2573-1599. PMID: 32833533 (Aug. 1, 2020) (cit. on p. 14).

84. Castel, B., Tomlinson, L., Locci, F., Yang, Y. & Jones, J. D. G. Optimization of T-DNA Architecture for Cas9-mediated Mutagenesis in Arabidopsis. PLOS ONE 14, e0204778. issn: 1932-6203 (Jan. 9, 2019) (cit. on p. 14).

85. Du, R., Flynn, M. J., Honsa, M., Jungmann, R. & Elowitz, M. B. miRNA Circuit Modules for Precise, Tunable Control of Gene Expression https://www.biorxiv.org/content/10.1101/2024.03.12.583048v1 (2024). Pre-published (cit. on p. 14).

86. Kabaria, S. R., et al. Programmable Promoter Editing for Precise Control of Transgene Expression https://www.biorxiv.org/content/10.1101/2024.06.19.599813v2 (2025). Pre-published (cit. on p. 14).

87. Aoki, S. K. et al. A Universal Biomolecular Integral Feedback Controller for Robust Perfect Adaptation. Nature 570, 533–537. issn: 1476-4687 (June 2019) (cit. on p. 14).

88. Wang, N. B., et al. Proliferation History and Transcription Factor Levels Drive Direct Conversion https://www.biorxiv.org/content/10.1101/2023.11.26.568736v1 (2023). Pre-published (cit. on pp. 14, 33).

89. Hansen, M. M. K. et al. A Post-Transcriptional Feedback Mechanism for Noise Suppression and Fate Stabilization. Cell 173, 1609–1621.e15. issn: 0092-8674 (June 14, 2018) (cit. on p. 14).

90. Vanderlinden, W., Skoruppa, E., Kolbeck, P. J., Carlon, E. & Lipfert, J. DNA Fluctuations Reveal the Size and Dynamics of Topological Domains. PNAS Nexus 1, pgac268. issn: 2752-6542 (Nov. 1, 2022) (cit. on p. 14).

91. Kim, E., Gonzalez, A. M., Pradhan, B., van der Torre, J. & Dekker, C. Condensin-Driven Loop Extrusion on Supercoiled DNA. Nat Struct Mol Biol. issn: 1545-9985. PMID: 35835864 (July 14, 2022) (cit. on p. 14).

92. Gao, X., Hong, Y., Ye, F., Inman, J. T. & Wang, M. D. Torsional Stiffness of Extended and Plectonemic DNA. Phys. Rev. Lett. 127, 028101 (July 7, 2021) (cit. on p. 14).

93. Corless, S., Naughton, C. & Gilbert, N. Profiling DNA Supercoiling Domains in Vivo. Genomics Data 2, 264–267. issn: 2213-5960 (Dec. 1, 2014) (cit. on p. 14).

94. Corless, S. & Gilbert, N. Investigating DNA Supercoiling in Eukaryotic Genomes. Briefings in Functional Genomics 16, 379–389. issn: 2041-2649 (Nov. 1, 2017) (cit. on p. 14).

95. Visser, B. J. & Bates, D. in Bacterial Chromatin: Methods and Protocols (ed Dame, R. T.) 147–156 (Springer US, New York, NY, 2024). isbn: 978-1-0716-3930-6 (cit. on p. 14).

96. Kouzine, F. et al. Transcription-Dependent Dynamic Supercoiling Is a Short-Range Genomic Force. Nat Struct Mol Biol 20, 396–403. issn: 1545-9985 (3 Mar. 2013) (cit. on p. 14).

97. Mark, M., Rijli, F. M. & Chambon, P. Homeobox Genes in Embryogenesis and Pathogenesis. Pediatr Res 42, 421–429. issn: 1530-0447 (Oct. 1997) (cit. on p. 15).

98. Engreitz, J. M. et al. Local Regulation of Gene Expression by lncRNA Promoters, Transcription and Splicing. Nature 539, 452–455. issn: 1476-4687 (7629 Nov. 2016) (cit. on p. 15).

99. Shearwin, K. E., Callen, B. P. & Egan, J. B. Transcriptional Interference – a Crash Course. Trends in Genetics 21, 339–345. issn: 0168-9525 (June 1, 2005) (cit. on p. 15).

100. Brooks, A. N. et al. Transcriptional Neighborhoods Regulate Transcript Isoform Lengths and Expression Levels. Science 375, 1000–1005 (Mar. 4, 2022) (cit. on p. 15).

101. Goldberg, B., Yehya, N., Xiao, J. & Meyer, S. Differential Effect of Supercoiling on Bacterial Transcription in Topological Domains. PLOS Computational Biology 21, e1012764. issn: 1553-7358 (Nov. 11, 2025) (cit. on p. 15).

102. Morao, A. K., Chervova, A., Zhao, Y., Ercan, S. & Cecere, G. DNA Supercoiling Modulates Eukaryotic Transcription in a Gene-Orientation Dependent Manner https://www.biorxiv.org/content/10.1101/2025.01.03.631213v1 (2025). Pre-published (cit. on p. 15).

103. Swanson, E. G., et al. Deaminase-Assisted Single-Molecule and Single-Cell Chromatin Fiber Sequencing https://www.biorxiv.org/content/10.1101/2024.11.06.622310v1 (2025). Pre-published (cit. on p. 15).

104. Yusa, K., Zhou, L., Li, M. A., Bradley, A. & Craig, N. L. A Hyperactive piggyBac Transposase for Mammalian Applications. Proc Natl Acad Sci U S A 108, 1531–1536. issn: 0027-8424. PMID: 21205896 (Jan. 25, 2011) (cit. on p. 24).

105. Aznauryan, E. et al. Discovery and Validation of Human Genomic Safe Harbor Sites for Gene and Cell Therapies. Cell Reports Methods 2, 100154. issn: 2667-2375 (Jan. 24, 2022) (cit. on p. 27).

106. Yang, T. et al. HiCRep: Assessing the Reproducibility of Hi-C Data Using a Stratum-Adjusted Correlation Coefficient. Genome Res 27, 1939–1949. issn: 1549-5469. PMID: 28855260 (Nov. 2017) (cit. on p. 28).

107. Koidl, S. & Timmers, H. T. M. greenCUT&RUN: Efficient Genomic Profiling of GFP-Tagged Transcription Factors and Chromatin Regulators. Current Protocols 1, e266. issn: 2691-1299 (2021) (cit. on p. 28).

108. Guo, M. S., Haakonsen, D. L., Zeng, W., Schumacher, M. A. & Laub, M. T. A Bacterial Chromosome Structuring Protein Binds Overtwisted DNA to Stimulate Type II Topoisomerases and Enable DNA Replication. Cell 175, 583–597.e23. issn: 00928674 (Oct. 2018) (cit. on p. 30).

