## Supplementary Figures for "Gene syntax defines supercoiling-mediated transcriptional feedback"

### Supplemental information

#### Materials and Methods

##### Bioinformatic analysis of intergene spacing across organisms

Genome annotations for *Saccharomyces cerevisiae*, *Mus musculus*, *Homo sapiens*, *Drosophila melanogaster*, *Caenorhabditis elegans*, and *Danio rerio* were downloaded from Ensembl, Release 110 (July 2023). Annotations with the type `gene`, `ncRNA_gene`, and `pseudogene` were selected and gene pair information (intergene spacing and orientation) was calculated for all gene pairs and gene trios. Gene pairs were split into equal-quantile bins based on intergene spacing, and the fraction of gene pairs with each orientation (divergent, tandem, or convergent) was computed for each bin. The frequency of the tandem orientation was normalized by dividing by two to account for the two possible syntaxes (downstream tandem and upstream tandem). For fig. S1d, a null hypothesis distribution for gene trios was generated by randomly selecting gene pairs with intergene spacings and two-gene orientations $(o_l, x_l)$ and $(o_r, x_r)$ and treating this as a gene trio with the given left and right spacings and orientations.

##### Cloning of genetic constructs

All plasmids were constructed using a combination of scarless NEB HiFi assembly (NEB E2621X) and an in-house Golden Gate cloning scheme that allows for facile multi-level assembly of gene cassettes and multi-cassette plasmids. The fluorescent proteins used for each construct are listed in table S1. All construct designs and source plasmid identifiers for every figure panel are listed in table S2. Plasmid descriptions are found in tables S3 to S5. Plasmids maps for all plasmids are included as supplemental files.

##### Human iPSC transfection and PiggyBac integration

iPS11 cells (Alstem, episomal HFF-derived) were maintained in mTeSR1, mTeSR Plus, or eTeSR media (StemCell Technologies) supplemented with 0.5% penicillin/streptomycin (GIBCO). Cells were grown in normoxia conditions (20% O2, 37°C, 5% CO2) on tissue culture-treated plastic plates coated for 30-60 min with Geltrex (Gibco) or Cultrex (Bio-Techne). For routine passaging, hiPSCs were dissociated to small clusters by (1) incubating in TrypLE Express (Gibco) for 2-4 minutes at 37°C, followed by quenching with mTeSR or eTeSR and light pipetting, or (2) incubating in ReLeSR (STEMCELL Tech) according to manufacturer's instructions. On thaw or passage with TrypLE, cells were treated with 5 µM ROCK inhibitor (ROCKi) Y-27632 (STEMCELL Tech).

For transfections, hiPSCs were collected and seeded at 75-100 x $10^3$ cells/mL in mTeSR1, mTeSR Plus, or eTeSR supplemented with 5 µM ROCKi Y-27632. The following day, cells were fed with fresh media and transfected with 80 ng (96w), 480 ng (24w), or 960 ng (12w) DNA complexed at a 3:1 ratio of µg DNA:µL Lipofectamine Stem Reagent (ThermoFisher STEM00015). The DNA mixtures typically comprised 6:1:1:1 ratio (by mass) of plasmids respectively encoding the PiggyBac transposon, hyperactive PiggyBac transposase (hyPBase [104]), Puromycin resistance gene (`pac`), and constitutive transfection marker (`mKO2`). On days 1 and 2 after transfection, cells were fed with fresh media supplemented with 0.5 µg/mL puromycin (Sigma P8833) to select for transfected cells transiently expressing `pac` from a non-integrating plasmid, thereby enriching for cells with PiggyBac integrations up to 90% purity. On subsequent days, cells were fed and passaged normally without puromycin, and subsampled during passages for flow cytometry.

##### Derivation of cell lines with matrix of constitutive promoters

Freshly passaged HEK293T cells maintained in DMEM + 10% FBS were seeded at 160k cells per well onto 0.1% gelatin-coated 24-well plates. The following day (0 dpt), cells were transfected with a 4:1 mass ratio of donor plasmid to PiggyBac supertranspoase, with a total transfected plasmid mass of 450 ng. Donor plasmids express two genes: one `mRuby2` transcript, and one `PuroR-T2A-mGreenLantern` transcript. At 1 day post transfection (dpt), cells were media changed into fresh media containing 1 µg / mL puromycin. Selection was maintained for two days (until 3 dpt), and surviving cells were passaged onto 6-well plates. After around a week of outgrowth (until each cell line was ∼30% confluent), 1 µg / mL puromycin was maintained for around two days, until no visible surviving cells in an untransfected control well were observed.

One day prior to sorting, single color control plasmids were transfected into fresh HEK293T cells for use in compensation. A Sony MA-900 was used to polyclonally sort cells. The majority of cell lines were double-positive sorted, using gates that exclude an untransfected control. Cell lines with extremely low expression (PGK-PGK) were sorted in a single-positive manner, where a cell was included if it was positive for mRuby2 or mGreenLantern. Cells were sorted into Pen-Strep supplemented conditioned media, containing 50% 0.2 micron-filtered media from a confluent plate of HEK293T cells and 50% fresh DMEM + 10% FBS. The resulting cells were outgrown and tested negative for mycoplasma.

**Lentivirus production**

To produce all lentiviral vectors, HEK293T Lenti-X cells (Takara 632180) were seeded at 7.0 to 7.5 million cells per 10cm dish coated with 0.1% gelatin. The following day, each plate was transfected with 6 µg transfer plasmid, 6 µg packaging plasmid (psPAX2, Addgene #12260), and 12 µg envelope plasmid (VSVG, Addgene #12259). Six hours later, each 10cm dish was media changed into 6.5 mL of HEPES-buffered (25 mM HEPES at pH 7.0) DMEM + 10% FBS. The following day, this media was collected, and fresh buffered media was added. The following day, these two aliquots were combined, filtered through a 0.45 µm PES filter, and combined with Lenti-X Concentrator (Takara 631232) overnight and concentrated following manufacturer instructions.

**Transduction of tandem syntax lentivirus**

Freshly passaged HEK293T cells were seeded at 15k cells per well onto 0.1% gelatin coated 96-well plates. The following day, cells were media changed into media containing 5 µg/mL polybrene (hexadimethrine bromide, Sigma-Aldrich, H9268-5G) and a serial dilution over 3.5 orders of magnitude (highest concentration: 5.0 µL concentrated virus per well). After two days, cells were dissociated and flowed. From the titration, representative concentrations were chosen with an MOI of approximately 0.5 for plotting in fig. S4c.

**Generation of monoclonal HEK293T cell lines**

Freshly passaged HEK293T cells were seeded at 20k cells per well onto 0.1% gelatin-coated 96-well plates. The following day, cells were transfected with a total of 340 ng of DNA, with a 1:1:1 mass ratio of PiggyBac supertransposase : two-gene tandem rtTA and PuroR donor : two-gene donor. On 1 dpt, 2 dpt, and 3 dpt, cells were media changed into fresh media containing 1 µg / mL puromycin. On 4 dpt, cells were passaged to 24-well scale and left to outgrow in media without puromycin. On 14 dpt, cells were passaged onto a 6-well plate. On 17 dpt, cells were passaged at high dilution to a new 6-well plate and maintained for the next ten days in dox-containing media.

One day prior to sorting, single color control plasmids were transfected for compensation. A Sony MA-900 was used to monoclonally sort double-positive cells, using gates that exclude an untransfected control. Specifically, individual cells were sorted into wells of a fresh 96-well plate with Pen-Strep supplemented conditioned media as described in the previous sections. Cells were allowed to outgrow over a period of two weeks. The resulting monoclonal cell lines tested negative for mycoplasma.

**Characterization of fluorescent protein expression in monoclonal lines**

30k cells per well of each monoclonal cell line were seeded onto 0.1% gelatin coated 96-well plates. The next day, media was changed into DMEM + 10% FBS containing dox concentrations from zero to 316 ng / mL dox. Two days later, cells were prepared for flow cytometry.

**Characterization of mRNA profile in monoclonal lines**

150k cells per well of each monoclonal line were seeded onto 0.1% gelatin coated 12-well plates. The next day, media was changed into media, with half the wells supplemented with 316 ng/mL dox. Two days later, cells were resuspended in PBS for HCR RNA-FISH.

Here, we use Molecular Instruments probe sets for TagBFP (compatible with B1 amplifiers conjugated to Alexa Fluor™ 647) and mRuby2 (compatible with B2 amplifiers conjugated to Alexa Fluor™ 514). We used the optimized FISH protocol as described in Peterman *et al.* [52]. Briefly, after suspension in PBS, cells were transferred to 96-well v-bottom plate for HCR Flow-FISH. After each resuspension, spins were performed at 500 rcf for 5 minutes with default settings, unless otherwise noted. Cells were first fixed through incubation in 4% PFA for 15 minutes at room temperature. After spinning, cells were then permeabilized using 0.5% Tween-20 for 15 minutes at room temperature. Next, cells were spun and resuspended in hybridization buffer for 30 minutes at 37°C. During this incubation, probe set stock solution was diluted in hybridization buffer to a concentration of 14 nM for transfected cells or 28 nM for integrated cell lines. Cells were spun and resuspended in this probe solution for incubation overnight at 37°C. Due to the viscosity of the hybridization buffer, these spins were performed with reduced deceleration speed to minimize cell loss.

Following hybridization, cells were spun and resuspended in wash buffer for 15 minutes at 37°C. Then, cells were spun and resuspended in 5X SSCT for five minutes at room temperature. After these washes, cells were spun and resuspended in amplification buffer for 30 minutes at room temperature. Amplifier solution was prepared by combining separately snap-cooled hairpins h1 and h2 at a concentration of 130 nM in amplification buffer. Cells were spun and then incubated with this amplifier solution overnight at room temperature.

Following amplification, cells were spun and resuspended in 5X SSCT for one 30 minute incubation and one five minute incubation at room temperature. Finally, cells were spun and resuspended in PBS for flow cytometry.

**Dox titration series and time course induction**

The monoclonal cell lines as described above were seeded at low confluency, 20k cells per well, onto 0.1% gelatin coated 24-well plates, in DMEM + 10% FBS containing 316 ng/mL dox. Independent wells were seeded for every experimental day. On each subsequent day from day 1 to day 13, cells were dissociated with trypsin and flowed as described in the Flow Cytometry section. Cells were media changed into media without dox at day 4. At day 7, the remaining relatively confluent cells were passaged onto fresh 24-well plates, at 12.5k cells per well, in media containing dox. At day 10, cells were media changed into media without dox.

**Derivation of integrated rtTA cell lines**

Freshly passaged HEK293T cells were seeded at 180k cells per well onto 0.1% gelatin coated 24-well plates. The following day (0 dpt), cells were transfected with a total of 90 ng of PiggyBac supertransposase and 225 ng of plasmid 41, a plasmid encoding for rtTA, SNAP-tag, and a zeocin resistance gene. The following day (1 dpt), cells were split onto a 6-well plate and media changed into media containing 1000 µg / mL zeocin. The following day (2 dpt), the selection media was replaced. On 3 dpt, cells were media changed into fresh media not containing zeocin. On 4 dpt, cells were passaged to a fresh 6-well plate and allowed to outgrow, then were frozen down. Later, the selected lines were unfrozen, allowed a passage to recover and were SNAP-stained immediately prior to sort (at 500:1 dilution). A Sony MA-900 was used to monoclonally sort SNAP-positive cells into individual wells of a 96-well plate containing Pen-Strep supplemented conditioned media. Six monoclones were outgrown and tested negative for mycoplasma.

For each of these lines, a test plasmid containing a TRE-inducible gene was transfected into each line, and doxycycline was added. Across the six monoclones, we selected the monoclone with a medium level of tight expression, as measured from the inducible gene. This monoclonal became our integrated rtTA line.

**Generation of hiPSC lines with circuits integrated at *CLYBL* or *AAVS1***

The STRAIGHT-IN Dual platform was used to integrate the all-in-one inducible circuits at both alleles of *CLYBL* [55]. A single copy of the all-in-one inducible circuit with each syntax was separately integrated into a "GT" STRIAGHT-IN landing pad located within intron 2 of one allele of *CLYBL*. Following the STRAIGHT-IN Dual protocol, 600 ng of the donor plasmid ("GT" plasmids 9-16 in table S3, including Addgene #229794) were transfected along with 400 ng of Bxb1-expressing plasmid (Addgene #51271) into 100,000 cells of the parental landing pad hiPSCs using Lipofectamine Stem Reagent (ThermoFisher STEM00015). The selection and excision of the lines was performed using the STRAIGHT-IN Dual protocol [55]. These cell lines were used for the timecourse induction experiment in fig. S10b.

Next, a second copy of the all-in one inducible circuits was integrated into a "GA" STRAIGHT-IN landing pad at the other allele of *CLYBL*. For this, donor plasmids ("GA" plasmids 9-16 in table S3, Addgene #229791 and #229792) were used, following the same protocol. These homozygously integrated lines were then used for the HCR RNA Flow-FISH, RNA-seq, RCMC, GapRUN, and CUT&Tag assays.

The same donor plasmids and protocol were used to integrate the circuits at a single allele of *AAVS1*, beginning with a parental hiPSC line containing the STRAIGHT-IN landing pad at this locus. Lastly, the circuits with varying gene lengths (donor plasmids 60-71 in table S3) in were integrated at a single allele of *CLYBL* following the same protocol.

These lines were maintained in StemFlex™ Medium (Thermo Scientific A3349401) with 100 U/mL Penicillin-Streptomycin (Gibco 15140122) on plates coated with 5 µg/mL Laminin-521 (STEMCELL Technologies 200-0117). Cells were passaged using Gentle Cell Dissociation Reagent (STEMCELL Technologies 100-1077) and cultured with media containing a 1:200 dilution of RevitaCell™ Supplement (Thermo Scientific A2644501) for 24 hours after plating. Media was replaced with fresh StemFlex + Pen/Strep every 2-3 days, and cells were grown at 37°C at 5% $CO_2$.

**Characterization of mRNA profile in homozygous hiPSC lines**

The four hiPSC lines with two-gene circuits homozygously integrated at *CLYBL* were cultured with or without 1 µg/mL dox for three days. Then, the cells were non-enzymatically dissociated using Gentle Cell Dissociation Reagent (STEMCELL Technologies 100-1077) and resuspended in PBS for HCR RNA-FISH. Molecular Instruments probe sets for mTagBFP2 (compatible with B1 amplifiers conjugated to Alexa Fluor™ 647) and mScarlet (compatible with B2 amplifiers conjugated to Alexa Fluor™ 514) were used, and the same protocol as for the monoclonal HEK293T lines was followed. The STRAIGHT-IN landing pad line without a circuit integrated was used as the parental condition to control for background RNA-FISH signal.

**RNA-seq**

The four hiPSC lines with two-gene circuits homozygously integrated at *CLYBL* were cultured with or without 1 µg/mL dox for three days. Then, the cells were non-enzymatically dissociated using Gentle Cell Dissociation Reagent (STEMCELL Technologies 100-1077), washed in PBS, and counted. 308,000 cells were collected per condition, and RNA was extracted using the Monarch Total RNA purification kit (NEB T2010) according to manufacturer's instructions. For each sample, 2 µL of a 1:100 dilution of ERCC spike-in RNA (Mix 1, Thermo Fisher 4456740) was added during the lysis step. RNA samples were sent to the MIT BioMicro Center for processing, using the NEB Ultra II Directional RNA with Poly(A) selection followed by library prep with unique dual indexes.

The pooled libraries were sequenced using 2x75bp paired reads on a single lane of an AVITI Cloudbreak Freestyle High chip (500M reads).

Paired-end reads were trimmed using version 0.6.10 of `trim-galore`. Trimmed paired-end reads were aligned to these genomes using `bowtie2` (version 2.5.4) using the `-very-sensitive-local` preset. Reads were deduplicated using `gatk`'s `MarkDuplicatesSpark` (version 4.6.2.0) and were cell-number normalized based on the number of ERCC-mapped reads. Normalized bigwig files were generated using `deeptools` (version 3.5.6).

**Region Capture Micro-C**

Three capture regions of interest, each roughly 0.5 Mb around three common landing pad integration sites were selected: *CLYBL* (chr13, 99,422,000-100,031,000), *AAVS1* (chr19, 54,810,000-55,715,000), and Rogi2 (chr3, 22,542,000-22,964,000) [105]. A custom probe panel of 80-mer probes targeting these regions and all synthetic parts inserted into these loci was designed and ordered from Twist Bioscience. Probes were selected based on a medium-stringency filter to reduce off-target pulldown, with some key areas near native TAD boundaries included using a low-stringency filter.

The homozygous hiPSC lines were expanded and outgrown at 6-well scale, with one plate each of the eight conditions (two syntaxes, two induction conditions, and two bioreplicates). Cells were dissociated with Gentle Cell Dissociation Reagent (STEMCELL Technologies 100-1077) according to the manufacturer's instructions and counted. From each of the eight conditions, 15M cells were collected and treated as described in Goel *et al.* [57]. An additional 25M cells were collected and for the MNase titration. Briefly, cells were washed in PBS and crosslinked using DSG (ThermoFisher 20593) and formaldehyde. After quenching, cells were washed, resuspended in PBS and counted, then resuspended in Micro-C buffer #1. Cell counts per condition varied between 6M and 13M cells.

A key variable in Micro-C is the ratio of MNase to cells. To identify ideal digestion concentrations, an MNase (Worthington Biochem LS004798) titration was performed on the separated aliquot. Increasing amounts of MNase were added and samples were purified and gel-separated. The optimal amount of MNase digests to primarily mononucleosomal fragments, with few but visible dinucleosomal and trinucleosomal bands. Using the optimal ratio, samples were digested with MNase, end-repaired, blunted, and labeled with biotinylated nucleotides. The biotin-labeled chromatin was then proximity ligated overnight. After enzymatic cleanup steps, the chromatin was then reverse crosslinked overnight. Dinucleosomes were selected using a gel extraction and T1 Streptavidin beads (Invitrogen 65601) were used to purify labeled fragments.

In order to determine the minimum number of cycles required to reach the required library concentration while minimizing PCR duplicates, a test amplification was performed with a small aliquot of each sample for a range of cycle numbers. The resulting PCRs were run on an agarose gel and quantified. Using this quantifiation, each sample was amplified to reach a target mass of 200 ng. NEB Multiplex Oligos for Illumina Primer Set 1 (NEB E7335) and NEBNext® High-Fidelity 2X PCR Master Mix (NEB M0541) were used for all PCRs, and sample barcodes were selected following the NEB recommendations. The resulting libraries were purified using AmPure XP beads (Beckman Coulter A63880) quantified via Bioanalyzer and qPCR using the NEBNext Library Quant kit for Illumina (NEB E7630).

The resulting libraries were mixed in equimass proportions and region-captured following Twist Bioscience's Standard Hybridization Target Enrichment Protocol, with the modification that a test PCR was used using the same reagents as above to identify an appropriate number of amplification cycles. The resulting region-capture libraries were purified and quantified via Bioanalyzer and qPCR. The pooled library was sequenced either via paired-end 2x75 cycle sequencing using Element's AVITI Cloudbreak Freestyle High flowcell (direct Illumina anchor sequencing without index conversion) for the divergent and downstream tandem conditions or the 2x150 cycle sequencing using a single lane of a NovaSeq X 10B flowcell for the convergent and upstream tandem conditions. Both methods gave about 500M reads per condition.

**RCMC bioinformatics**

For each integrated syntax, custom human genomes were generated by editing the GRCh38 genome build to contain the inserted synthetic construct at the *CLYBL* landing pad. Paired-end reads were trimmed using version 0.6.10 of `trim-galore`. Trimmed paired-end reads were aligned to these genomes using `bowtie2` using the `-very-sensitive-local` preset. Then, `pairtools` was used to identify Hi-C pairs (with `-walks-policy mask` and `-min-mapq 2`) and deduplicate them (with `-max-mismatch 1`). The resulting reads were converted to `.mcool` format using `cooltools`. The resulting matrices were ICE balanced (within each capture region for the majority of this work, and globally for fig. S12b) and binned into bins at both the 500 bp and 2000 bp resolution.

To evaluate the reproducibility of this method, the stratum-corrected correlation coefficient as implemented in Yang *et al.* [106] was used to evaluate the similarity of all samples, both within the capture region of interest fig. S12a and in distal regions fig. S12c. Based on the high reproduciblity observed, reads from the two bioreplicates were merged for the rest of this work.

Two distal, unmodified capture regions did not show induction-dependent changes (fig. S12c).

Using the balanced contact matrices, fold-changes in contact probability upon induction are calculated by dividing these matrices by each other. The sliding-window insulation score in fig. 3d is calculated using the `insulation` function in `cooltools` and uses a sliding window of 50 kb. The anti-diagonal score presented in fig. 3e sums the contact probability along the anti-diagonal (i.e., perpendicular to the main diagonal) up to a given distance, centered at a given genomic coordinate. Specifically, we use the balanced contact matrices, binned at 500 bp resolution, and sum bins within 5 kb of the target location. For a given bin at location $x$, this means summing the probability of the $(x - 500\,\text{bp})$ bin contacting the $(x + 500\,\text{bp})$ bin to the probability of the $(x - 1000\,\text{bp})$ bin contacting the $(x + 1000\,\text{bp})$ bin.

**Producing nanobody-MNase fusion protein**

The following protocol is modified and scaled down from the protein production protocol in Koidl & Timmers [107] and the GapRUN protocol in Longo *et al.* [17]. GapRUN relies on an MNase-nanobody fusion protein that is not commercially available. However, the protein can be reliably produced at a scale sufficient for roughly 24 reactions without needing a FPLC or sonicator. At a high-level, protein production proceeds by inducing bacterial cells overnight at 18°C, lysing the cells, clarifying the lysate, and binding the fusion protein to glutathione spin columns. After successive washes, the fusion protein is cleaved off the column using biotin-tagged thrombin protease. The protease is removed from the eluate using streptavidin beads, and the resulting fusion protein is concentrated and buffer is exchanged using spin columns.

The following buffers were prepared. First, sample buffer is 10% glycerol, 2% SDS, 50 mM Tris, 20 mM EDTA, and 1% $\beta$-mercaptoethanol (for 10 mL, this is 2 mL of 50% glycerol, 500 µL of 1M Tris pH 6.8, 2 mL 10% SDS, 400 µL 0.5M EDTA, and 100 µL of $\beta$-mercaptoethanol, with Milli-Q grade water added to 10 mL). The $\beta$-mercaptoethanol should be added shortly before use. Bromophenol blue dye can optionally be added to give the buffer color, which is useful when loading protein gels. Second, the protein purification wash buffer is 125 mM Tris, 150 mM NaCl at pH 8.0.

An IPTG-inducible protein production plasmid encoding for GST-LaG16-MNase (Addgene 170978) was transformed into BL21 (DE3) competent cells. The resulting bacteria were streaked to single colonies, and a single colony was used to start a 10 mL overnight culture at 37°C in LB-Amp media. The following day, 800 mL of LB-Amp (split into 200 mL volumes distributed across 4 baffled 1 liter flasks) was inoculated with 8 mL of the overnight culture (1:100 dilution). The cells were grown to an OD600 of 0.5, which takes about two hours. Once the flasks reached an OD of 0.5 mL, a 1.5 mL "pre-induction" sample was taken, spun down, and resuspended in 300 µL of sample buffer, then stored at -20°C for later analysis. Each flask containing 200 mL of media was induced with 200 µL of 1M IPTG (sterile filtered with 0.2 micron filters) and the flasks were placed back in the shake incubator at a temperature of 18°C overnight. The lower temperature is required to maintain stability of the produced protein. The following morning, the OD600 was measured on a 5-fold diluted sample, and found to be be around 6.0. A "post-induction" 500 µL sample was also taken, spun down, and resuspended in 300 µL of sample buffer, and stored at -20°C for later analysis.

Cells were pelleted at maximum speed in a fixed-angle rotor. After combining pellets together, the cells were resuspended in complete B-Per buffer (Thermo Fisher 78248) plus EDTA-free protease inhibitor (Thermo Fisher A32965). Specifically, for every gram of cell pellet, we resuspended it in 4 mL of B-Per plus 2 µL of lysosome and DNAse I (both included in the B-Per kit) plus 40 µL of 100x protease inhibitor cocktail (1 protease tablet in 500 µL of Milli-Q grade water). After incubation at room temperature for 15 minutes, the slurry was subjected to two freeze thaw cycles to -80°C to further lyse the cells. The resulting lysate was spun at 14k xg in a fixed-angle rotor for 30 minutes at 4°C. A 50 µL sample of the clarified lysate was diluted to 300 µL in sample buffer and stored at -20°C for later analysis.

The resulting clarified lysate was diluted 2:1 with protein purification wash buffer (for every 2 mL of lysate, add 1 mL of wash buffer). Two 0.2 mL glutathione spin columns (Thermo Fisher 16106) were washed with wash buffer as directed by the manufacturer instructions, then the diluted clarified lysate was loaded onto the columns. This takes many sequential spins or a strong vacuum manifold! After loading, the columns were washed with 2 column volumes (400 µL) of wash buffer until the A280 absorbance (measured with a NanoDrop) dropped to background levels, about 20 column volumes total. To cleave the fusion protein from the column, 100 units of biotin-tagged thrombin protease (Sigma-Aldrich SAE0147-5KU) diluted to 200 µL in wash buffer was added to each capped spin column and incubated overnight at 4°C.

The following day, the columns were spun, collecting "elution 1". 1 column volume (200 µL) of wash buffer was added and the columns were again spun, for "elution 2", and so on for a third elution. 10 µL samples of the elutions were combined with 10 µL of sample buffer, for later analysis. The A280 absorbance of these elutions were checked, and the first two elutions were combined. To remove the thrombin protease, the combined elution (400 µL) was combined in low-binding 1.7mL tubes with 60 µL of T1 streptavidin beads (Thermo Fisher 65601) that had been washed three times with PBS (each wash consisting of magnetically separating the beads, removing the supernatant, and resuspending in 200 µL of PBS). The resulting bead slurry was rotated at 4°C for 30 minutes. The beads were magnetically removed, and the elution fraction was transferred to fresh low-binding tubes. A 10 µL sample of the combined elution was combined with 10 µL of sample buffer for later analysis.

Fresh 1M dithiothreitol (DTT) was prepared and used to make Buffer A: 20 mM K2PO4, 10% glycerol, 0.5 mM EDTA, 1 mM DTT (for 5 mL, this is 1 mL 0.1M K2PO4, pH 7.0, 0.5 mL glycerol, 5 µL 0.5M EDTA, and 5 µL 1M DTT). After 0.22 micron filter sterilization, 100x protease inhibitor cocktail was spiked into the buffer.

To concentrate and buffer exchange the protein, the elution fraction was added to 10 kDa MWCO protein concentrator spin columns (Thermo Fisher 88513) and concentrated to a volume of approximately 100 µL. The concentrated protein was buffer exchanged into Buffer A by adding prepared Buffer A on top and concentrating four times (e.g., if concentrated to 100 µL, add 100 µL, concentrate back to 100 µL, add 100 µL, concentrate to 100 µL, and so on). Finally, the protein was concentrated to a volume of 75 µL, removed from the concentrator, and mixed with 75 µL of 100% glycerol to reach 150 µL of purified protein, to be stored at -20C.

To confirm proper protein production, an acrylamide protein gel was cast. 10 µL aliquots of each sample saved from above were briefly heated to 95°C prior to cooling on the bench, and these samples were loaded, ran on the protein gel, and stained with Coomassie to confirm protein purification. A single band was visible in the elution samples.

**Titrating nanobody-MNase activity**

An MNase activity analysis was performed to estimate the activity of the purified fusion protein. hiPSC cells were collected as in the above section for RCMC and as described in Goel *et al.* [57], crosslinked only with formaldehyde (DSG is not necessary for a simple activity assay), quenched, washed, and flash frozen in aliquots of 5M cells. Cell pellets were resuspended in Micro-C Buffer #1, split into 1M aliquots, and were then incubated with a range of both 1) commercial MNase and 2) the purified LaG16-MNase fusion protein. The cells were incubated, quenched, reverse-crosslinked, and had DNA purified as described in the MNase titration step in Goel *et al.* [57]. Benchmarking the LaG16-MNase activity based on the commercial MNase, we determined around 8 units of LaG16-MNase was required per million cells (in our case, about 2 of the purified protein sample per million cells).

**Measuring positive supercoiling with GapRUN**

Lentiviral vectors containing the GapR-GFP fusion protein (pTA571, EF1α-GapR-FLAGx3-NLS-EGFP) were cloned. Plasmids for the constructs in Longo *et al.* [17] were not available on Addgene, so a DNA fragment (Twist Biosciences) was ordered containing a human-codon-optimized sequence encoding for GapR-3xFLAG-NLS-EGFP. The GapR sequence was derived from the cDNA for CCNA_03428, a.k.a. GapR as identified in Guo *et al.* [108]. This fragment was cloned into a Golden-Gate compatible donor plasmid via HiFi assembly, which was then assembled with Golden Gate assembly alongside the EF1α promoter and the bGH polyadenylation signal into a second-level donor plasmid. This expression unit was Golden Gate assembled into a LentiX1 backbone. Lentivirus was produced as described above, then titer was computed by performing a serial dilution, transducing Lenti-X1 cells seeded at 40k cells per well of a 96-well plate. These cells were flowed three days later, gated on positive GFP signal, and the dilution series was fit to a Poisson distribution to calculate the infectious units per µL.

To prepare cells for GapRUN, hiPSCs homozygously integrated with the two-gene circuits at *CLYBL* were cultured with or without 1µg/mL dox for three days. On the second day of induction, cells were spinfected with the the GapR virus. Namely, fresh media containing virus at an MOI of 0.8 plus 5 µg/mL polybrene (hexadimethrine bromide, Sigma-Aldrich H9268), with or without dox, was added to the cells, and the plate was spun at 1,500xg at 32°C for 90 minutes. Conditions without virus and without dox were included, giving 12 conditions total (virus and dox, virus only, or neither for each syntax). The day after spinfection, cells were changed to fresh media with or without dox. The following day, cells were non-enzymatically dissociated using Gentle Cell Dissociation Reagent (STEMCELL Technologies 100-1077), washed in PBS, and counted.

In advance, stock solutions of 1M HEPES-NaOH at pH 7.5, 500 mM spermidine, 5M NaCl, 2.5M CaCl2, 0.5M EDTA, and 0.5M EGTA were prepared. A 100x solution of protease inhibitor cocktail (PIC, Thermo Fisher A32965) was prepared by dissolving one tablet in 500 µL of 20 mM HEPES-NaOH and 150 mM NaCl.

On the first day, 700 µL of Wash Buffer per sample was prepared: Wash Buffer is 20 mM HEPES-NaOH, 150 mM NaCL, and 0.5 mM spermidine (for 5 mL, this is 100 µL 1M HEPES-NaOH, 150 µL 5M NaCl, and 5 µL 500 mM spermidine). Digitonin (Cell Signaling Technologies 16359) was warmed to room temperature, pipetted to mix, then heated briefly to 95°C (e.g., in PCR tubes) before cooled on ice. When mentioned below, Wash Buffer + PIC (100 µL Wash Buffer plus 1 µL PIC) or Complete Wash Buffer (100 µL Wash Buffer plus 1 µL PIC plus 2.5 µL digitonin) was prepared immediately before use.

10 µL Concanavalin A beads (Cell Signaling Technology 93569) per reaction were activated by magnetically separating the beads, removing the supernatant, and resuspending each in 100 µL of activation buffer (included with CST 93569). After 10 minutes at room temperature, the activation step was repeated, then the beads were separated and resuspended in 10 µL of activation buffer. Aliquots of 500k cells (dissociated and counted above) were washed twice with 100 µL of Wash Buffer + PIC, then resuspended in 100 µL of Wash Buffer + PIC. Each aliquot was combined with the 10 µL of activated ConA beads, mixed via trituration, and placed on a Nutator at room temperature for ten minutes.

The beads were then separated and resuspended in 50 µL of Complete Wash Buffer. Around 8 units of LaG16-MNase was spiked into the mixture, mixed well via trituration, and nutated overnight at 4°C.

The following day, 500 µL of Wash Buffer was prepared per sample. The beads were separated and washed twice with 200 µL of Complete Wash Buffer, transferring the beads to new PCR tubes after the first wash to reduce background signal. The beads were resuspended in 50 µL of Complete Wash Buffer, mixed gently, and placed on ice for at least two minutes. 1 µL of 150 mM $CaCl_2$ was spiked into each reaction while still on ice, and the samples were nutated at 4°C for two hours.

Before the 2 hours were up, 55 µL of Stop Buffer was prepared per sample. Stop Buffer is 340 mM NaCl, 20 mM EDTA, 10 mM EGTA, 2.5% digitonin, 50 µg/mL glycogen, and 100 µg/mL RNase A (for 500 µL, this is 17 µL 5M NaCL, 20 µL 500 mM EDTA, 10 µL 500 mM EGTA, 12.5 µL digitonin, 1.25 µL 20 mg/mL glycogen, and 2.5 µL 20 mg/mL RNase A). The glycogen and RNase A was added last, shortly before use.

Samples were removed from the Nutator and placed on ice. 50 µL of Stop Buffer was added to each sample, mixed well via trituration, and incubated on a thermocycler for 10 minutes at 37°C. After transferring the samples to new low-binding 1.7 mL tubes, DNA was extracted following the Monarch Spin PCR & DNA Cleanup kit instructions for small-fragment retention (NEB T1130). DNA was eluted in 50 µL of 1x TE.

The resulting libraries were prepared following the NEBNext Ultra II DNA library kit instructions (NEB E7645), not performing size selection by following step 3B instead of 3A. Test PCRs (PCRs on 1 µL samples of the pre-amplification library) were performed on the resulting pre-amplification library to determine the number of PCR cycles required to reach sequencing submission requirements, around 13 cycles. The resulting libraries were uniquely dual-indexed and pooled for sequencing.

The pooled libraries were sequenced using 2x75bp paired reads on an entire AVITI Cloudbreak Freestyle High chip (1B reads).

Paired-end reads were trimmed using version 0.6.10 of `trim-galore`. Trimmed paired-end reads were aligned to these genomes using `bowtie2` (version 2.5.4) using the `-very-sensitive-local` preset. Reads were deduplicated using `gatk`'s `MarkDuplicatesSpark` (version 4.6.2.0). Bigwig files were generated using `deeptools` (version 3.5.6).

Genomic tracks were compared to control conditions that were not transduced with the GapR lentivirus. Signal in these control conditions was significantly lower than in the experimental conditions (fig. S29).

**Measuring polymerase occupancy and histone marks with CUT&Tag**

hiPSCs homozygously integrated with the two-gene circuits at *CLYBL* were cultured with or without 1 µg/mL dox for three days. Cells were then non-enzymatically dissociated using Gentle Cell Dissociation Reagent (STEMCELL Technologies 100-1077), washed in PBS, and counted. Four rabbit antibodies were used, targeting the following: Pol II (ser2) (Cell Signaling Technologies 13499S), H3K27ac (Active Motif 39133), H3K27me3 (Active Motif 91167), and H3K4me3 (Active Motif 39915). Aliquots of 500k cells per condition and per antibody were processed following manufacturer instructions for the CUT&Tag-IT Express kit (Active Motif 53177), with the following exceptions. First, because we had four antibody conditions per cell condition, "pre-aliquots" of 2M cells were made with spike-in *Drosophila* nuclei (Active Motif 53168) added prior to splitting into 500k aliquots, in order to reduce spike-in noise across antibody conditions. Second, instead of using Q5 polymerase and a fixed number of cycles, we used NEBNext Ultra II Q5 (NEB M0544) and did an initial test PCR (PCRs on 1 µL samples of the pre-amplification library) to determine the number of PCR cycles. Finally, Nextera-compatible combinatorial dual-index primers were used to index the libraries (Active Motif 53155).

The pooled libraries were sequenced using 2x75bp paired reads on an entire AVITI Cloudbreak Freestyle High chip (1B reads).

Paired-end reads were trimmed using version 0.6.10 of `trim-galore`. Trimmed paired-end reads were aligned to these genomes using `bowtie2` (version 2.5.4) using the `-very-sensitive-local` preset. Reads were deduplicated using `gatk`'s `MarkDuplicatesSpark` (version 4.6.2.0) and were cell-number normalized based on the number of *Drosophila*-mapped reads from the spike-in nuclei. Normalized bigwig files were generated using `deeptools` (version 3.5.6).

**Generation of antibody producer cell lines**

To integrate two-gene antibody production cassettes, we used a Bxb1-mediated landing pad line with attP receptor site at Rogi2 in HEK293Ts for parallel integration of each cassette Peterman *et al.* [52]. The landing pad functions analogously to the STRAIGHT-IN landing pad Blanch-Asensio *et al.* [55]. The landing pad contains a truncated puromycin resistance gene missing the promoter and start codon. Upon Bxb1-mediated recombination between the attB site on the donor plasmid and attP site in the landing pad, an EF1α promoter and start codon is placed in-frame of the resistance gene, conferring recombinant cells resistance to puromycin.

To integrate the donor plasmids, the landing pad HEK293T line was seeded at 100k cells per well onto 0.1% gelatin coated 24-well plates. The following day, the cells were transfected using 450 ng of two-gene donor plasmid and 300 ng of CAG-Bxb1 (gift from the Wong Lab at Boston University). At 1dpt, the cells were media changed. At 2 dpt, the cells were passaged to 0.1% gelatin coated 6-well plates. At 3 dpt, the cells were media changed into media containing 1 µg / mL puryomycin. Selection was maintained until an untransfected well was fully selected against (around 1.5 weeks). Once confluent (five to six days of puromycin administration), cells were passaged at a split ratio of 1:10 to dilute out residual donor plasmid, at which point cells were ready for use in downstream analyses.

**Titering of antibody production yields**

After outgrowth and passaging of the antibody producer lines, triplicates were seeded on 6-well plates at 500k cells per 6-well. The day after, cells were media changed into 2 mL of fresh DMEM + 10% FBS. After six days of outgrowth, 1.1 mL of supernatant was collected from every condition and centrifuged at 10k g for 10 minutes to remove cell debris. The top 1.0 mL of clarified supernatant was transferred to Pierce 35 kDa PES protein concentrator columns (Thermo Scientific 88502). Each sample was centrifuged to a final volume less than 75 µL. Each concentrated sample was diluted to 75 µL with pH 7.4 PBS.

The resulting concentrated samples were processed following manufacturer instructions using a Human IgG (Total) ELISA kit (Thermo Scientific BMS2091) and a Easy-Titer Human IgG (H+L) kit (Thermo Scientific 23310). Absorbance was measured on a Tecan Infinite M1000 Pro at 450nm (ELISA) and 340nm (Easy-Titer).

**Lentiviral transduction of two-gene cassettes**

Lentiviruses were produced as described above. To calculate lentiviral titer, a two-fold serial dilution of the concentrated lentivirus was combined with 5 µg/mL polybrene (hexadimethrine bromide, Sigma-Aldrich H9268) and 20,000 HEK293T cells per well in a 0.1% gelatin-coated 96-well plate. The following day, cells were changed into fresh DMEM + 10% FBS. Two days later, cells were flowed. The fraction of expressing cells in each dilution condition was used to compute viral titer, assuming a Poisson process for infection. Viral titers were then used to calculate the volume of concentrated virus needed to infect cells at the desired MOI.

On the day of transduction, 5k HEK293T cells in suspension were co-infected with virus expressing the two-gene cassette of different syntax at MOI of 0.8, and 3 µL of lentivirus expressing rtTA-P2A-mGreenLantern at high MOI. Cells were plated into a 96-well with DMEM + 10% FBS containing 5 µg/mL polybrene (hexadimethrine bromide, Sigma-Aldrich H9268). The following day, media was replaced with fresh DMEM + 10% FBS. At three days post infection, cells were passaged into 6-well plates. Once confluent, cells were flow-sorted on the Sony-MA900 for double-positive constitutive reporter gene and rtTA-mGreenLantern. The polyclonal sorted lines where then re-plated post-sorting and passaged consistently to maintain cells at 80% confluence or lower.

For evaluating effect of doxycycline addition, the post-sorted, integrated HEK293T cell lines were re-plated into a 96-well plate at 39k cells/well with DMEM + 10% FBS. The day after plating, media was replaced with DMEM + 10% FBS contained various concentration of doxycycline from 1 µg/mL to 0 µg/mL. Cells were flowed at three days post doxycycline treatment. Bioreplicates represent different passages of the post-sorted, integrated cell lines.

**Lentiviral transduction of the engineered all-in-one inducible circuits**

Lentiviruses were produced as described above. To calculate lentiviral titer, a two-fold serial dilution of the concentrated lentivirus was combined with 5 µg/mL polybrene (hexadimethrine bromide, Sigma-Aldrich H9268) and 20k HEK293T cells per well in a 0.1% gelatin-coated 96-well plate. The following day, cells were changed into fresh DMEM + 10% FBS containing 1 µg/mL doxycycline. Two days later, cells were flowed. The fraction of expressing cells in each dilution condition was used to compute viral titer, assuming a Poisson process for infection. Viral titers for iPS11 cells and mouse embryonic fibroblasts (MEFs) were estimated as two-fold greater or four-fold lower, respectively, than for HEK293T cells. Viral titers were then used to calculate the volume of concentrated virus needed to infect cells at an MOI of 0.3.

To transduce HEK293T cells, 20k cells were combined with 5 µg/mL polybrene and the calculated amount of concentrated virus. The next day, cells were changed into fresh DMEM + 10% FBS with or without 300 ng/mL dox. Two days later, cells were flowed.

To transduce iPS11 cells, cells were dissociated with Gentle Cell Dissociation Reagent according to manufacturer's instructions, then seeded at 15k cells/well in a Geltrex-coated 96-well plate with mTeSR-Plus and 5 µM ROCKi. The next day, the media was replaced with fresh mTeSR-Plus containing the calculated amount of concentrated virus and 5 µg/mL polybrene. Plates were then spun at 1500 x g for 90 minutes. The following day, media was changed to fresh mTeSR-Plus with or without 300 ng/mL dox. Two days later, cells were imaged and flowed.

To transduce MEFs, passage 1 primary MEFs—isolated as described in Wang *et al.* [88]—were thawed and allowed to recover for 1-2 days in DMEM + 10% FBS. Cells were then dissociated with Trypsin-EDTA and seeded at 10k cells per well in a 0.1% gelatin-coated 96-well plate. The following day, the media was replaced with fresh DMEM + 10% FBS containing the calculated amount of concentrated virus and 5 µg/mL polybrene. Plates were then spun at 1500 x g for 90 minutes. The following day, media was changed to fresh DMEM + 10% FBS with or without 300 ng/mL dox. Two days later, cells were flowed. Biological replicates include cells from multiple independent isolations.

**Flow cytometry**

Hanks' Balanced Salt Solution (HBSS) supplemented with 2% FBS (PiggyBac-integrated iPS11 cells) or PBS (all others) was used as Flow buffer. Cells were washed once with PBS, then dissociated with Trypsin-EDTA (HEK293T, MEFs), TrypLE Express (iPS11), or Gentle Cell Dissociation Reagent (iPS11). After dissociation, cells were quenched with Flow buffer or media, then spun down for 5 min at 300-500 x g. Cells were resuspended in Flow buffer and acquired on CytoFLEX LX N3-V5-B3-Y5-R3-I0 (Beckman Coulter) or Attune NxT Flow Cytometer (Thermo Fisher). Data were gated for single cells on the FlowJo software (v10.X; BD Biosciences) or CytExpert (v2.6; Beckman Coulter), then exported for plotting and additional analysis.

**Intrinsic and extrinsic noise analysis**

To compute the intrinsic and extrinsic noise, we first log-transform each of the gene expression levels. This log-transform ensures that changes in the values correspond to fold-change in protein concentration independent of fluorescent protein quantum yield and PMT values for cytometry channels. We calculate the variance

$$\mathrm{var}(\vec{x}) = \frac{1}{n}\sum_{i=1}^{N}(x_i - \overline{x})^2$$

of the sum and difference of the values for each gene. The intrinsic noise is the variance of the difference of the log-transformed expression levels. The extrinsic noise is the variance of the sum of the log-transformed expression levels. The intrinsic and extrinsic noise between genes $a$ and $b$ is thus:

$$n_{\text{intrinsic}} = \mathrm{var}(\log(\vec{a}) - \log(\vec{b}))$$
$$n_{\text{extrinsic}} = \mathrm{var}(\log(\vec{a}) + \log(\vec{b}))$$

Dividing the intrinsic noise by the extrinsic noise gives the intrinsic-to-extrinsic noise ratio.

**Copy number quantification with droplet digital PCR**

The copy number of the PiggyBac-integrated fluorescent reporter lines (fig. S9) and the Rogi2-integrated antibody production lines (fig. S20b) was evaluated using Bio-Rad's QX200 Droplet Digital PCR system.

Briefly, two sets of fluorescent PCR probes were ordered. First, probes targeting the mRuby2 coding sequence with the FAM fluorophore, and probes targeting RPP30, a native gene on chromosome 10, with the HEX fluorophore. Genomic DNA is isolated from the cell lines of interest and diluted to an appropriate concentration for PCR. Using the Bio-Rad ddPCR system, the resulting PCR mix is split into many nanoliter-sized droplets, PCR is performed, and each droplet is readout. Then, the fraction of the double-positive and single-positive populations is fit to a Poisson distribution. Because the RPP30 has a known copy number, the Poisson fits give an estimate of the mean and variance in copy number of the mRuby2 gene. As the mRuby2 gene is present once in each of the integrated constructs, this allows us to calculate the total number of integrations.

For the PiggyBac integrated lines (fig. S9), we observe copy numbers between 4 and 18, with the divergent syntax having the lowest copy number despite having the highest expression. For the Rogi2 antibody production lines (fig. S20b), we expect a single integration and observe copy numbers between 1 and 1.4. The fractional copy number comes from the relative genomic instability of the HEK293Ts. As the HEK293Ts divide, chromosomal rearrangements and duplications can occur, making the population heterogeneous. Still, the antibody production lines with the highest titer have the lowest copy number.

### A Supplemental figures and tables

| Figure | Reporter / Constitutive | Adjacent / Inducible |
|---|---|---|
| figs. 1c, S3 and S6 | TagBFP | mNeonGreen |
| figs. 2, 5b to 5d, S7, S21 and S22 | mRuby2 | TagBFP |
| figs. 3c, S10, S11, S24e and S24f | mTagBFP2 | mScarlet |
| figs. 6, S2, S4a, S4b, S23, S24b to S24d, S25 and S26 | mGreenLantern (mGL) | mRuby2 |
| fig. S27 | mGL | mCherry |
| **Figure** | **Upstream** | **Downstream** |
| fig. S4c | none | mRuby2 |
| fig. S5 | TagBFP | mNeonGreen |

**Table S1:** Mapping between described genes and fluorophores

| Figure | Cell type | Plasmids | Integration | Selection information |
|---|---|---|---|---|
| fig. 1c | iPS11 | 3 (DT), 4 (UT) | PiggyBac | Transient puromycin |
| figs. 2 and S7 to S9 | HEK293T | 5 + {6 (DT), 7 (C), 8 (D)} | PiggyBac | puromycin, monoclonal sort |
| figs. 3, 4, 6e, S10c to S10e, S11 to S19, S24e, S24f, S28 and S29 | STRAIGHT-IN hiPSC | 9, 10 (DT); 11, 12 (D); 13, 14 (UT); 15, 16 (C) | Bxb1 landing pad | STRAIGHT-IN |
| figs. 5a and S20 | Rogi2 landing pad HEK293T | 17 (UT), 18 (DT), 19 (C), 20 (D) | Bxb1 landing pad | puromycin |
| figs. 5b to 5d, 6e, S21 and S22 | HEK293T | 21/22 + 23 (UT), 24 (DT), 25 (C), 26 (D) | Lentivirus | Polyclonal sort |
| figs. 6b to 6f, S24b and S25c | iPS11 | 28 (DT), 29 (Div), 27 (UT) | Lentivirus | None |
| figs. 6f and S25c, cHS4 core | iPS11 | 42 (D), 43 (DT), 44 (UT) | Lentivirus | None |
| figs. 6f and S25c, cHS4 full | iPS11 | 45 (D), 46 (DT), 47 (UT) | Lentivirus | None |
| figs. S2 and S4b, PGK-PGK | HEK293T | 1 (UT), 2 (DT) | PiggyBac | puromycin, polyclonal sort |
| fig. S3a, downstream insulation | iPS11 | 72 (UT), 73 (DT) | PiggyBac | Transient puromycin |
| fig. S3a, upstream insulation | iPS11 | 74 (UT), 75 (DT) | PiggyBac | Transient puromycin |
| fig. S3a, full insulation | iPS11 | 76 (UT), 77 (DT) | PiggyBac | Transient puromycin |
| fig. S3b | iPS11 | 108 - 115 | PiggyBac | Transient puromycin |
| figs. S4a and S4b, EFS-EFS | HEK293T | 33 (UT), 34 (DT) | PiggyBac | puromycin, polyclonal sort |
| figs. S4a and S4b, EF1α-EF1α | HEK293T | 35 (UT), 36 (DT) | PiggyBac | puromycin, polyclonal sort |
| figs. S4a and S4b, EF1α-PGK | HEK293T | 37 (UT), 38 (DT) | PiggyBac | puromycin, polyclonal sort |
| figs. S4a and S4b, PGK-EF1α | HEK293T | 39 (UT), 40 (DT) | PiggyBac | puromycin, polyclonal sort |
| fig. S4c | HEK293T | 30, 31, 32 | Lentivirus | None |
| fig. S5 | iPS11 | 78 - 101 | PiggyBac | Transient puromycin |
| fig. S6 | iPS11 | 78, 102, 103 (D); 83, 104, 105 (UT); 3, 106, 107 (DT) | PiggyBac | Transient puromycin |
| fig. S10b | STRAIGHT-IN hiPSC | 10 (DT), 12 (D), 14 (UT), 16 (C) | Bxb1 landing pad | STRAIGHT-IN |
| figs. 6e, S23, S24c, S24d, S25a, S25b, S25d and S25e | HEK293T, MEF | 28 (DT), 29 (Div), 27 (UT) | Lentivirus | None |
| fig. S25, cHS4 core | iPS11, HEK293T, MEF | 42 (D), 43 (DT), 44 (UT) | Lentivirus | None |
| fig. S25, cHS4 full | iPS11, HEK293T, MEF | 45 (D), 46 (DT), 47 (UT) | Lentivirus | None |
| fig. S26, original 0.5 kb | HEK293T | 55 (D), 54 (DT), 56 (UT) | Lentivirus | None |
| fig. S26, original 1.0 kb | HEK293T | 58 (D), 57 (DT), 59 (UT) | Lentivirus | None |
| fig. S26, TF inert 0.3 kb | HEK293T | 49 (D), 48 (DT), 50 (UT) | Lentivirus | None |
| fig. S26, TF inert 1.8 kb | HEK293T | 52 (D), 51 (DT), 53 (UT) | Lentivirus | None |
| fig. S27, 0.7 kb | STRAIGHT-IN hiPSC | 60 (DT), 61 (D), 62 (UT), 63 (C) | Bxb1 landing pad | STRAIGHT-IN |
| fig. S27, 1.4 kb | STRAIGHT-IN hiPSC | 64 (DT), 65 (D), 66 (UT), 67 (C) | Bxb1 landing pad | STRAIGHT-IN |
| fig. S27, 5.2 kb | STRAIGHT-IN hiPSC | 68 (DT), 69 (D), 70 (UT), 71 (C) | Bxb1 landing pad | STRAIGHT-IN |

**Table S2:** Constructs, cell type, and integration method is listed for every data panel.

| Ref | Identifier | Syntax | Construct |
|---|---|---|---|
| 1 | pTA292 | U Tan | PGK-Puro-T2A-mGL-SV40 PGK-mRuby2-bGH |
| 2 | pTA289 | D Tan | PGK-mRuby2-bGH PGK-Puro-T2A-mGL-SV40 |
| 3 | aRJ134 | D Tan | PGK-mNeonGreen-syn_pA EF1α-TagBFP-syn_pa |
| 4 | aRJ135 | U Tan | EF1α-TagBFP-syn_pa PGK-mNeonGreen-syn_pa |
| 5 | pTA187 | Tan | CMV-rtTA-βglobin_pa CMV-PuroR-bGH |
| 6 | pTA129 | D Tan | TRE-TagBFP-bGH EF1α-mRuby2-SV40 |
| 7 | pTA130 | Con | TRE-TagBFP-bGH (EF1α-mRuby2-SV40) |
| 8 | pTA131 | Div | (TRE-TagBFP-bGH) EF1α-mRuby2-SV40 |
| 9 | pABA03 | D Tan | GA allele, TRE3G-mScarlet-WPRE CAG-rtTA-T2A-mTagBFP2-bGH |
| 10 | pABA04 | D Tan | GT allele, TRE3G-mScarlet-WPRE CAG-rtTA-T2A-mTagBFP2-bGH |
| 11 | pABA01 | Div | GA allele, (TRE3G-mScarlet-WPRE) CAG-rtTA-T2A-mTagBFP2-bGH |
| 12 | pABA02 | Div | GT allele, (TRE3G-mScarlet-WPRE) CAG-rtTA-T2A-mTagBFP2-bGH |
| 13 | pKG3040 | U Tan | GA allele, CAG-rtTA-T2A-mTagBFP2-bGH TRE3G-mScarlet-WPRE |
| 14 | pKG4157 | U Tan | GT allele, CAG-rtTA-T2A-mTagBFP2-bGH TRE3G-mScarlet-WPRE |
| 15 | pKG3038 | Con | GA allele, CAG-rtTA-T2A-mTagBFP2-bGH (TRE3G-mScarlet-WPRE) |
| 16 | pKG4155 | Con | GT allele, CAG-rtTA-T2A-mTagBFP2-bGH (TRE3G-mScarlet-WPRE) |
| 17 | pTA515 | U Tan | CMV-mAb17HC-bGH CMV-mAb17LC-P2A-mRuby2-bGH |
| 18 | pTA516 | D Tan | CMV-mAb17LC-P2A-mRuby2-bGH CMV-mAb17HC |
| 19 | pTA517 | Con | CMV-mAb17HC-bGH (CMV-mAb17LC-P2A-mRuby2-bGH) |
| 20 | pTA518 | Div | (CMV-mAb17HC-bGH) CMV-mAb17LC-P2A-mRuby2-bGH |
| 21 | pKG2752 | n/a | LX1, EFS-rtTA-P2A-mGL-WPRE |
| 22 | pTA169 | n/a | LX1, CMV-rtTA-bGH |
| 23 | pTA444 | U Tan | LX1, EF1α-mRuby2-bGH TRE-TagBFP-bGH |
| 24 | pTA443 | D Tan | LX1, TRE-TagBFP-bGH EF1α-mRuby2-bGH |
| 25 | pTA445 | Con | LX1, EF1α-mRuby2-bGH (TRE-TagBFP-bGH) |
| 26 | pTA446 | Div | LX1, (TRE-TagBFP-bGH) EF1α-mRuby2-bGH |
| 27 | pKG3549 | U Tan | EFS-rtTA-P2A-mGL-WPRE original_0.3kb TRE-mRuby2-bGH |
| 28 | pKG1484 | D Tan | TRE-mRuby2-bGH original_0.3kb EFS-rtTA-P2A-mGL-WPRE |
| 29 | pKG1487 | Div | (TRE-mRuby2-bGH) original_0.3kb EFS-rtTA-P2A-mGL-WPRE |

**Table S3:** Plasmids used in main-text figures. Text in parentheses indicates a construct placed in the antisense direction.

| Ref | Identifier | Syntax | Construct |
|---|---|---|---|
| 30 | pGEEC270 | D Tan | LX1, PGK-SNAPtag-bGH EF1α-mRuby2-WPRE |
| 31 | pGEEC272 | D Tan | LX1, UbC-SNAPtag-bGH EF1α-mRuby2-WPRE |
| 32 | pGEEC276 | n/a | LX1, EF1α-mRuby2-WPRE |
| 33 | pTA291 | U Tan | EFS-Puro-T2A-mGL-SV40 EFS-mRuby2-bGH |
| 34 | pTA288 | D Tan | EFS-mRuby2-bGH EFS-Puro-T2A-mGL-SV40 |
| 35 | pTA293 | U Tan | EF1α-Puro-T2A-mGL-SV40 EF1α-mRuby2-bGH |
| 36 | pTA290 | D Tan | EF1α-mRuby2-bGH EF1α-Puro-T2A-mGL-SV40 |
| 37 | pTA345 | U Tan | PGK-Puro-T2A-mGL-SV40 EF1α-mRuby2-bGH |
| 38 | pTA342 | D Tan | EF1α-mRuby2-bGH PGK-Puro-T2A-mGL-SV40 |
| 39 | pTA344 | U Tan | EF1α-Puro-T2A-mGL-SV40 PGK-mRuby2-bGH |
| 40 | pTA343 | D Tan | PGK-mRuby2-bGH EF1α-Puro-T2A-mGL-SV40 |
| 41 | pTA309 | n/a | UbC-rtTA-P2A-SNAPtag-T2A-ZeoR-bGH |
| 42 | pKG3572 | Div | (TRE-mRuby2-bGH) EFS-rtTA-P2A-mGL-WPRE |
| 43 | pKG3571 | D Tan | TRE-mRuby2-bGH cHS4.core EFS-rtTA-P2A-mGL-WPRE |
| 44 | pKG3574 | U Tan | EFS-rtTA-P2A-mGL-WPRE cHS4.core TRE-mRuby2-bGH |
| 45 | pKG3576 | Div | (TRE-mRuby2-bGH) cHS4.full EFS-rtTA-P2A-mGL-WPRE |
| 46 | pKG3575 | D Tan | TRE-mRuby2-bGH cHS4.full EFS-rtTA-P2A-mGL-WPRE |
| 47 | pKG3578 | U Tan | EFS-rtTA-P2A-mGL-WPRE cHS4.full TRE-mRuby2-bGH |
| 48 | pAIO203 | D Tan | TRE-mRuby2-bGH TF.inert_0.3kb EFS-rtTA-P2A-mGL-WPRE |
| 49 | pAIO204 | Div | (TRE-mRuby2-bGH) TF.inert_0.3kb EFS-rtTA-P2A-mGL-WPRE |
| 50 | pAIO205 | U Tan | EFS-rtTA-P2A-mGL-WPRE TF.inert_0.3kb TRE-mRuby2-bGH |
| 51 | pAIO206 | D Tan | TRE-mRuby2-bGH TF.inert_1.8kb EFS-rtTA-P2A-mGL-WPRE |
| 52 | pAIO207 | Div | (TRE-mRuby2-bGH) TF.inert_1.8kb EFS-rtTA-P2A-mGL-WPRE |
| 53 | pAIO208 | U Tan | EFS-rtTA-P2A-mGL-WPRE TF.inert_1.8kb TRE-mRuby2-bGH |
| 54 | pAIO212 | D Tan | TRE-mRuby2-bGH original_0.5kb EFS-rtTA-P2A-mGL-WPRE |
| 55 | pAIO213 | Div | (TRE-mRuby2-bGH) original_0.5kb EFS-rtTA-P2A-mGL-WPRE |
| 56 | pAIO214 | U Tan | EFS-rtTA-P2A-mGL-WPRE original_0.5kb TRE-mRuby2-bGH |
| 57 | pAIO215 | D Tan | TRE-mRuby2-bGH original_1.0kb EFS-rtTA-P2A-mGL-WPRE |
| 58 | pAIO216 | Div | (TRE-mRuby2-bGH) original_1.0kb EFS-rtTA-P2A-mGL-WPRE |
| 59 | pAIO217 | U Tan | EFS-rtTA-P2A-mGL-WPRE original_1.0kb TRE-mRuby2-bGH |
| 60 | pAIO254 | D Tan | TRE-mCherry-WPRE CAG-rtTA-P2A-mGL-bGH |
| 61 | pAIO255 | Div | (TRE-mCherry-WPRE) CAG-rtTA-P2A-mGL-bGH |
| 62 | pAIO256 | U Tan | CAG-rtTA-P2A-mGL-bGH TRE-mCherry-WPRE |
| 63 | pAIO257 | Con | TRE-mCherry-WPRE (CAG-rtTA-P2A-mGL-bGH) |
| 64 | pAIO258 | D Tan | TRE-VP16-ZF43-P2A-mCherry-WPRE CAG-rtTA-P2A-mGL-bGH |
| 65 | pAIO259 | Div | (TRE-VP16-ZF43-P2A-mCherry-WPRE) CAG-rtTA-P2A-mGL-bGH |
| 66 | pAIO260 | U Tan | CAG-rtTA-P2A-mGL-bGH TRE-VP16-ZF43-P2A-mCherry-WPRE |
| 67 | pAIO261 | Con | TRE-VP16-ZF43-P2A-mCherry-WPRE (CAG-rtTA-P2A-mGL-bGH) |
| 68 | pAIO262 | D Tan | TRE-BEmax-dCas12a-P2A-mCherry-WPRE CAG-rtTA-P2A-mGL-bGH |
| 69 | pAIO263 | Div | (TRE-BEmax-dCas12a-P2A-mCherry-WPRE) CAG-rtTA-P2A-mGL-bGH |
| 70 | pAIO264 | U Tan | CAG-rtTA-P2A-mGL-bGH TRE-BEmax-dCas12a-P2A-mCherry-WPRE |
| 71 | pAIO265 | Con | TRE-BEmax-dCas12a-P2A-mCherry-WPRE (CAG-rtTA-P2A-mGL-bGH) |

**Table S4:** Plasmids used in supplemental figures. Text in parentheses indicates a segment placed in the antisense direction.

| Ref | Identifier | Syntax | Construct |
|---|---|---|---|
| 72 | aRJ137 | U Tan | EF1α-TagBFP-Syn_pA D1-insulator PGK-mNG-Syn_pA E1-insulator |
| 73 | aRJ136 | D Tan | PGK-mNG-Syn_pA D1-insulator EF1α-TagBFP-Syn_pA E1-insulator |
| 74 | aRJ133 | U Tan | D1-insulator EF1α-TagBFP-Syn_pA E1-Insulator PGK-mNG-Syn_pA |
| 75 | aRJ132 | D Tan | D1-insulator PGK-mNG-Syn_pA E1-insulator EF1α-TagBFP-Syn_pA |
| 76 | aRJ139 | U Tan | D1-insulator EF1α-TagBFP-Syn_pA PGK-mNG-Syn_pA E1-insulator |
| 77 | aRJ138 | D Tan | D1-insulator PGK-mNG-Syn_pA EF1α-TagBFP-Syn_pA E1-insulator |
| 78 | aRJ210 | Div | (PGK-mNeonGreen-Syn_pA) EF1α-TagBFP-bGH |
| 79 | aRJ211 | Tan | PGK-mNeonGreen-Syn_pA PGK-TagBFP-Syn_pA |
| 80 | aRJ212 | Tan | PGK-mNeonGreen-Syn_pA UbC-TagBFP-Syn_pA |
| 81 | aRJ213 | Tan | PGK-mNeonGreen-Syn_pA CAG-TagBFP-Syn_pA |
| 82 | aRJ214 | Tan | EF1α-mNeonGreen-Syn_pA EF1α-TagBFP-Syn_pA |
| 83 | aRJ215 | Tan | EF1α-mNeonGreen-Syn_pA PGK-TagBFP-Syn_pA |
| 84 | aRJ216 | Tan | EF1α-mNeonGreen-Syn_pA UbC-TagBFP-Syn_pA |
| 85 | aRJ217 | Tan | EF1α-mNeonGreen-Syn_pA CAG-TagBFP-Syn_pA |
| 86 | aRJ218 | Tan | UbC-mNeonGreen-Syn_pA EF1α-TagBFP-Syn_pA |
| 87 | aRJ219 | Tan | UbC-mNeonGreen-Syn_pA PGK-TagBFP-Syn_pA |
| 88 | aRJ220 | Tan | UbC-mNeonGreen-Syn_pA UbC-TagBFP-Syn_pA |
| 89 | aRJ221 | Tan | UbC-mNeonGreen-Syn_pA CAG-TagBFP-Syn_pA |
| 90 | aRJ222 | Tan | CAG-mNeonGreen-Syn_pA EF1α-TagBFP-Syn_pA |
| 91 | aRJ223 | Tan | CAG-mNeonGreen-Syn_pA PGK-TagBFP-Syn_pA |
| 92 | aRJ224 | Tan | CAG-mNeonGreen-Syn_pA UbC-TagBFP-Syn_pA |
| 93 | aRJ225 | Tan | CAG-mNeonGreen-Syn_pA CAG-TagBFP-Syn_pA |
| 94 | aRJ242 | Tan | PGK-mNeonGreen-Syn_pA Inert-TagBFP-Syn_pA |
| 95 | aRJ243 | Tan | EF1α-mNeonGreen-Syn_pA Inert-TagBFP-Syn_pA |
| 96 | aRJ244 | Tan | UbC-mNeonGreen-Syn_pA Inert-TagBFP-Syn_pA |
| 97 | aRJ245 | Tan | CAG-mNeonGreen-Syn_pA Inert-TagBFP-Syn_pA |
| 98 | aRJ246 | Tan | Inert-mNeonGreen-Syn_pA PGK-TagBFP-Syn_pA |
| 99 | aRJ247 | Tan | Inert-mNeonGreen-Syn_pA EF1α-TagBFP-Syn_pA |
| 100 | aRJ248 | Tan | Inert-mNeonGreen-Syn_pA UbC-TagBFP-Syn_pA |
| 101 | aRJ249 | Tan | Inert-mNeonGreen-Syn_pA CAG-TagBFP-Syn_pA |
| 102 | aRJ228 | Div | (PGK-mNeonGreen-Syn_pA) EF1α-TagBFP-bGH |
| 103 | aRJ229 | Div | (PGK-mNeonGreen-bGH) EF1α-TagBFP-Syn_pA |
| 104 | aRJ230 | Tan | EF1α-mNeonGreen-Syn_pA PGK-TagBFP-bGH |
| 105 | aRJ231 | Tan | EF1α-mNeonGreen-bGH PGK-TagBFP-Syn_pA |
| 106 | aRJ232 | Tan | PGK-mNeonGreen-Syn_pA EF1α-TagBFP-bGH |
| 107 | aRJ233 | Tan | PGK-mNeonGreen-bGH EF1α-TagBFP-Syn_pA |
| 108 | aRJ260 | Tan | PGK-mNeonGreen D1-CTCF EF1α-TagBFP |
| 109 | aRJ261 | Tan | PGK-mNeonGreen D1-mutant-CTCF EF1α-TagBFP |
| 110 | aRJ262 | Tan | PGK-mNeonGreen (E1-CTCF) EF1α-TagBFP |
| 111 | aRJ263 | Tan | PGK-mNeonGreen (E1-scramble-CTCF) EF1α-TagBFP |
| 112 | aRJ264 | Tan | PGK-mNeonGreen D1-CTCF (E1-CTCF) EF1α-TagBFP |
| 113 | aRJ265 | Tan | PGK-mNeonGreen D1-scramble-CTCF (E1-scramble-CTCF) EF1α-TagBFP |
| 114 | aRJ266 | Tan | PGK-mNeonGreen (E1-CTCF) D1-CTCF EF1α-TagBFP |
| 115 | aRJ267 | Tan | PGK-mNeonGreen (E1-scramble-CTCF) D1-scramble-CTCF EF1α-TagBFP |

**Table S5:** Plasmids used in PiggyBac-integrated hiPSC supplemental figures. Text in parentheses indicates a segment placed in the antisense direction.

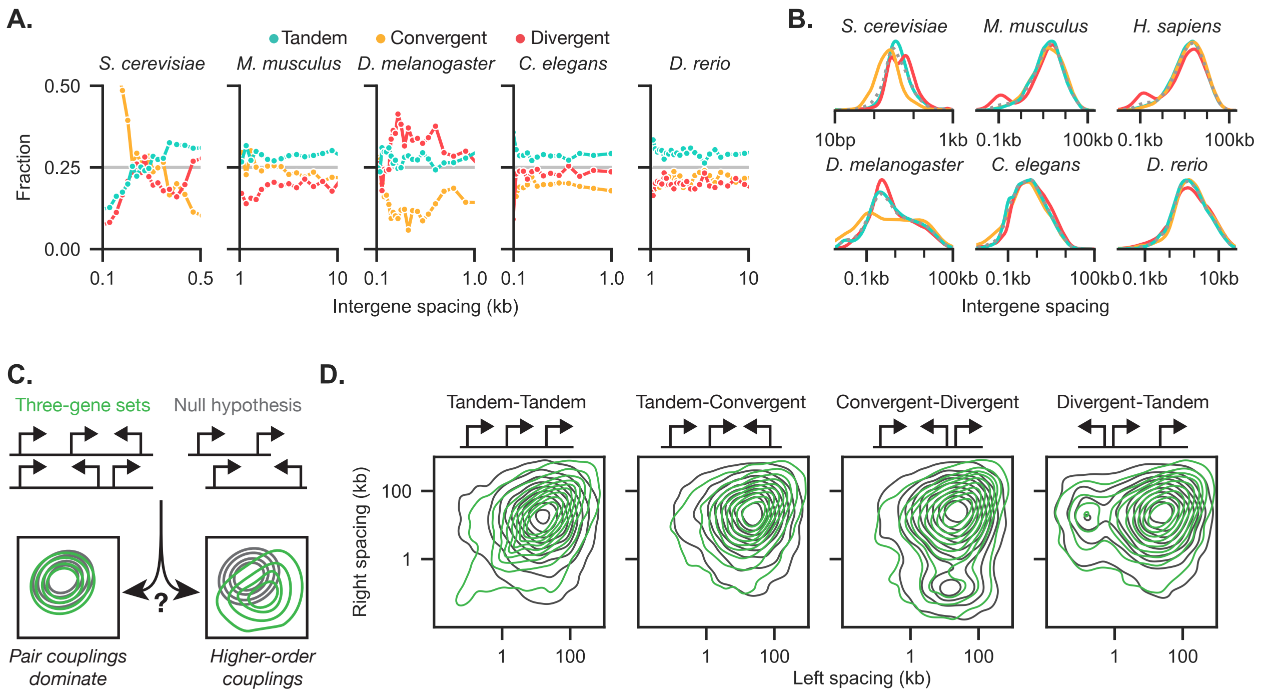


**Figure S1:** Bioinformatic analysis of gene pairs and trios in eukaryotic genomes.
a) Syntax distributions for gene pairs binned by intergene spacing are shown for *Saccharomyces cerevisiae*, *Mus musculus*, *Drosophila melanogaster*, *Caenorhabditis elegans*, and *Danio rerio*.
b) Instead of binning all gene pairs by intergene spacing as in fig. 1b, gene pairs were separated by syntax and the intergene spacing distributions were compared to the overall intergene distribution (gray, dashed).
c) Higher order couplings between trios of genes can be evaluated by comparing the joint distribution of the left and right intergene spacings for all pairs of genes, compared to a null hypothesis where gene trios are synthetically created by combining gene pairs.
d) For the human genome, three-gene pairs can be split into four syntaxes. The distribution of intergene spacings are compared to the null hypothesis where gene pairs are randomly combined to form three-gene trios. The overlap of the distributions indicates that trios were largely explained by the pairs.

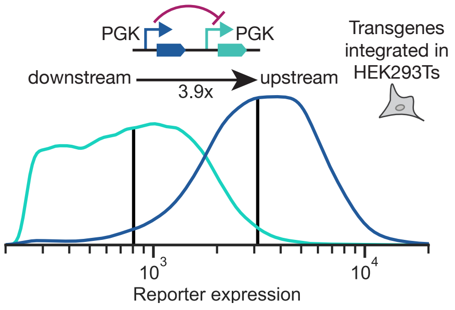


**Figure S2:** Transgenes randomly integrated using PiggyBac into HEK293Ts show upstream dominance. The expression of the upstream gene is around four-fold higher than the downstream gene when the same fluorescent protein is placed in successive positions. Geometric means are shown as black vertical lines.

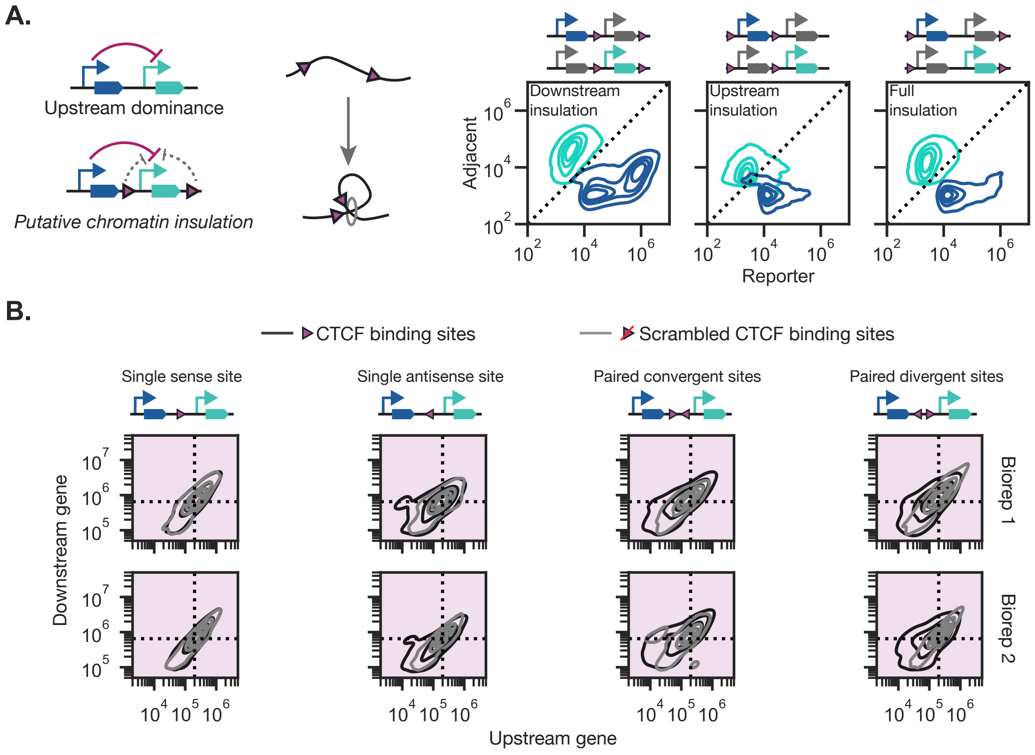


**Figure S3:** a) Flanking CTCF sites may insulate a chromatin region from its surroundings. Tandem-oriented CTCF binding sites were introduced flanking the upstream gene, the downstream gene, or the entire construct. Joint distributions of the two genes are shown.
b) Four variants of CTCF-based insulators and one tDNA-based insulator were tested for insulation activity. These insulators were "inactivated" by scrambling the CTCF binding motif or by removing the B-box, respectively. Only the transcriptionally-active insulator showed significant insulation, decreasing upstream expression while increasing downstream expression.

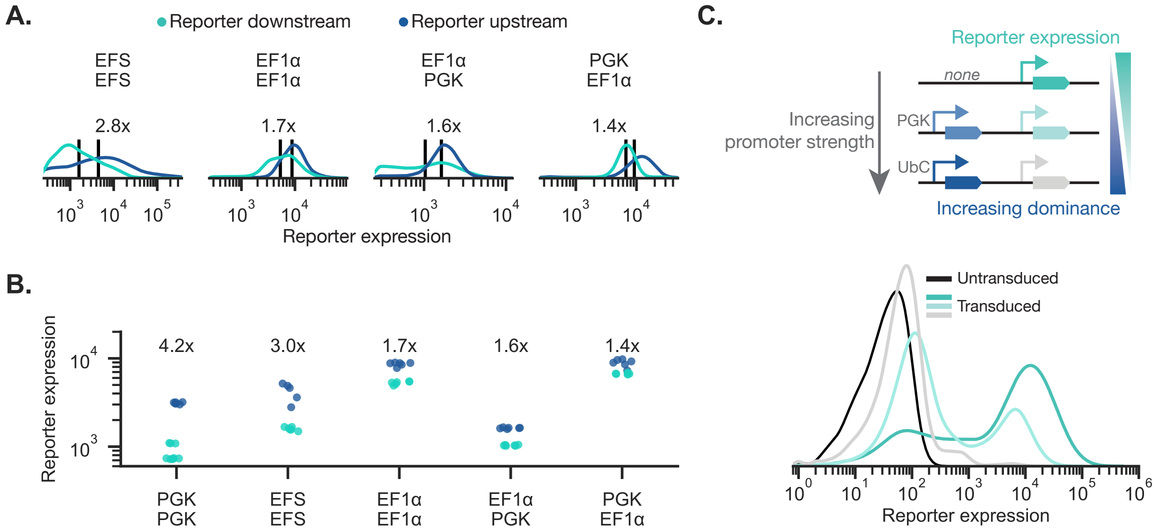


**Figure S4:** Two-gene constitutive gene pairs show upstream dominance.
a) Representative distributions of a reporter gene in the upstream and downstream positions for different combinations of constitutive promoters PiggyBac-integrated into HEK293T cells. All promoter pairs demonstrate upstream dominance.
b) The geometric mean of the reporter expression for each biological replicate (N=5), with the median upstream dominance fold change shown above.
c) Three lentiviruses with the same constant downstream reporter but different upstream genes were transduced into HEK293T cells. Stronger upstream expression reduces downstream reporter expression.

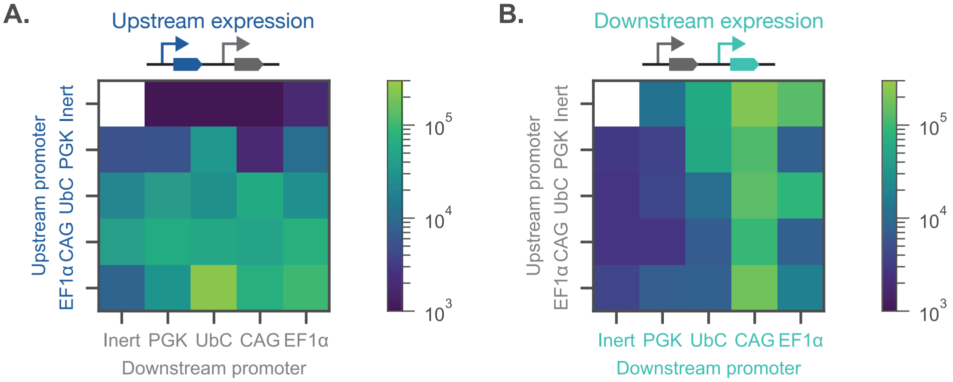


**Figure S5:** Expression patterns of tandem two-gene constructs PiggyBac-integrated into hiPSC lines.
A panel of constitutive promoters were placed in both the upstream and downstream positions and integrated into hiPSCs. Average of measurements from three biological replicates, (N=3).
a) Upstream expression for every tested promoter combination.
b) Downstream expression for every tested promoter combination.

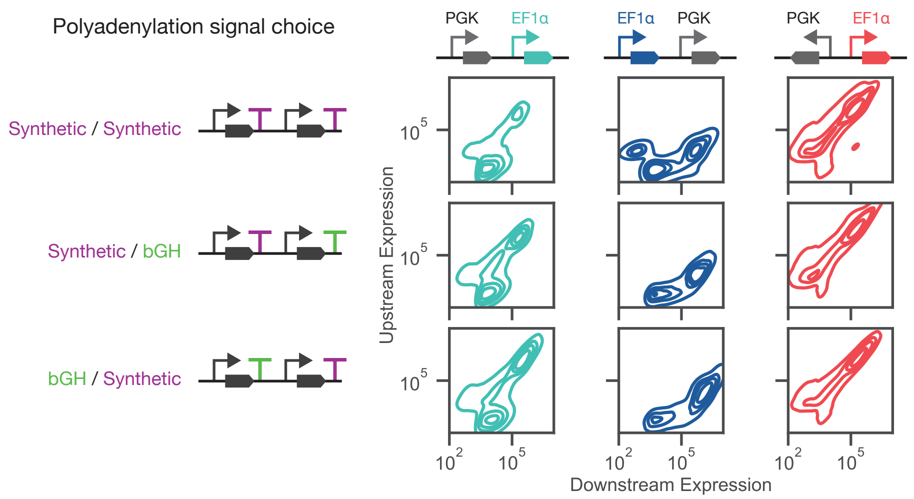


**Figure S6:** For three different combinations of polyadenylation signals, representative joint distributions for PiggyBac-integrated hiPSCs are shown. The PAS signal choice only minimally affects the resulting distributions for each syntax. Each column shows similar joint distributions.

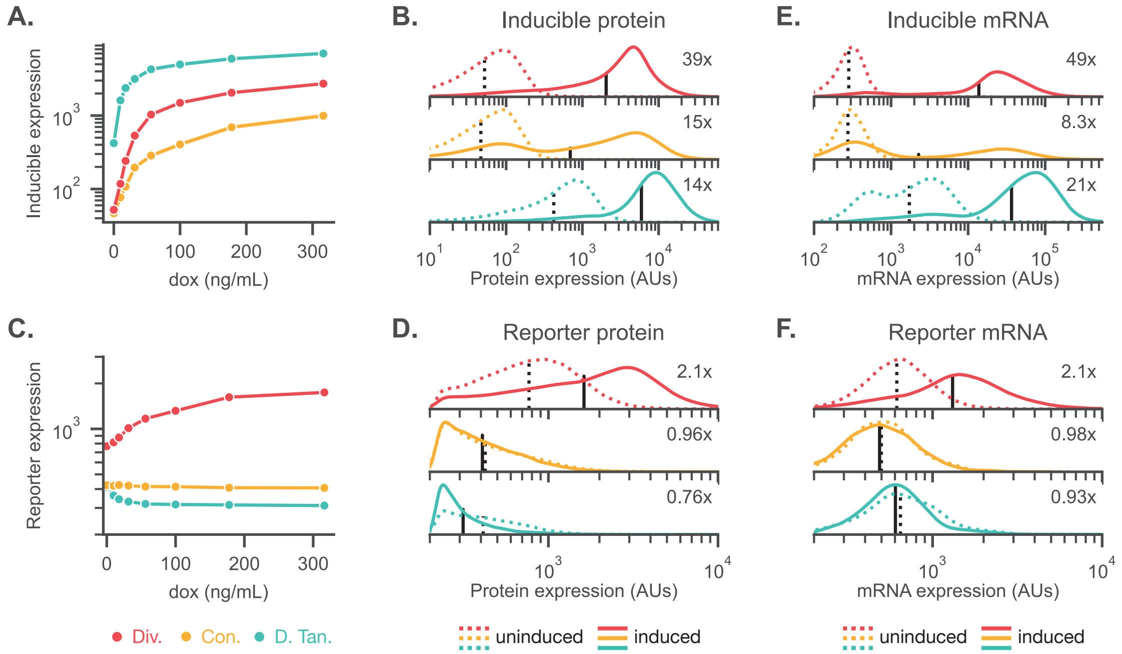


**Figure S7:** Circuit syntax affects both mRNA and protein levels.
a) The geometric mean of the un-normalized inducible gene expression is shown as a function of dox induction. The geometric mean and 95% confidence interval are calculated over three merged wells.
b) Representative protein distributions are shown for the inducible gene in both the uninduced (dashed) and induced (solid) cases.
c) The geometric mean of the un-normalized reporter expression is shown as a function of dox induction. The geometric mean and 95% confidence interval are calculated over three merged wells.
d) Representative protein distributions are shown for the reporter gene in both the uninduced (dashed) and induced (solid) cases.
e),f) Representative mRNA distributions are shown for the inducible and reporter genes in both the uninduced (dashed) and induced (solid) cases.

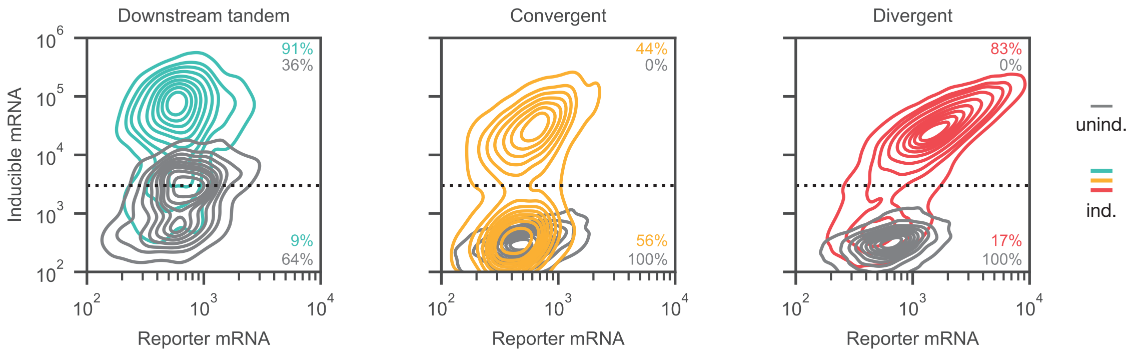


**Figure S8:** Joint mRNA distributions are shown for the monoclonal lines presented in fig. 2 for the uninduced (gray) and induced (colored) conditions.

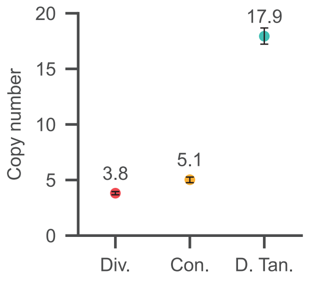


**Figure S9:** Measured copy number for the monoclonal PiggyBac lines presented in fig. 2. Copy number is measured via ddPCR probes that bind to the mRuby2 gene and normalized to the RPP30 gene. Error bars show the 95% confidence interval.

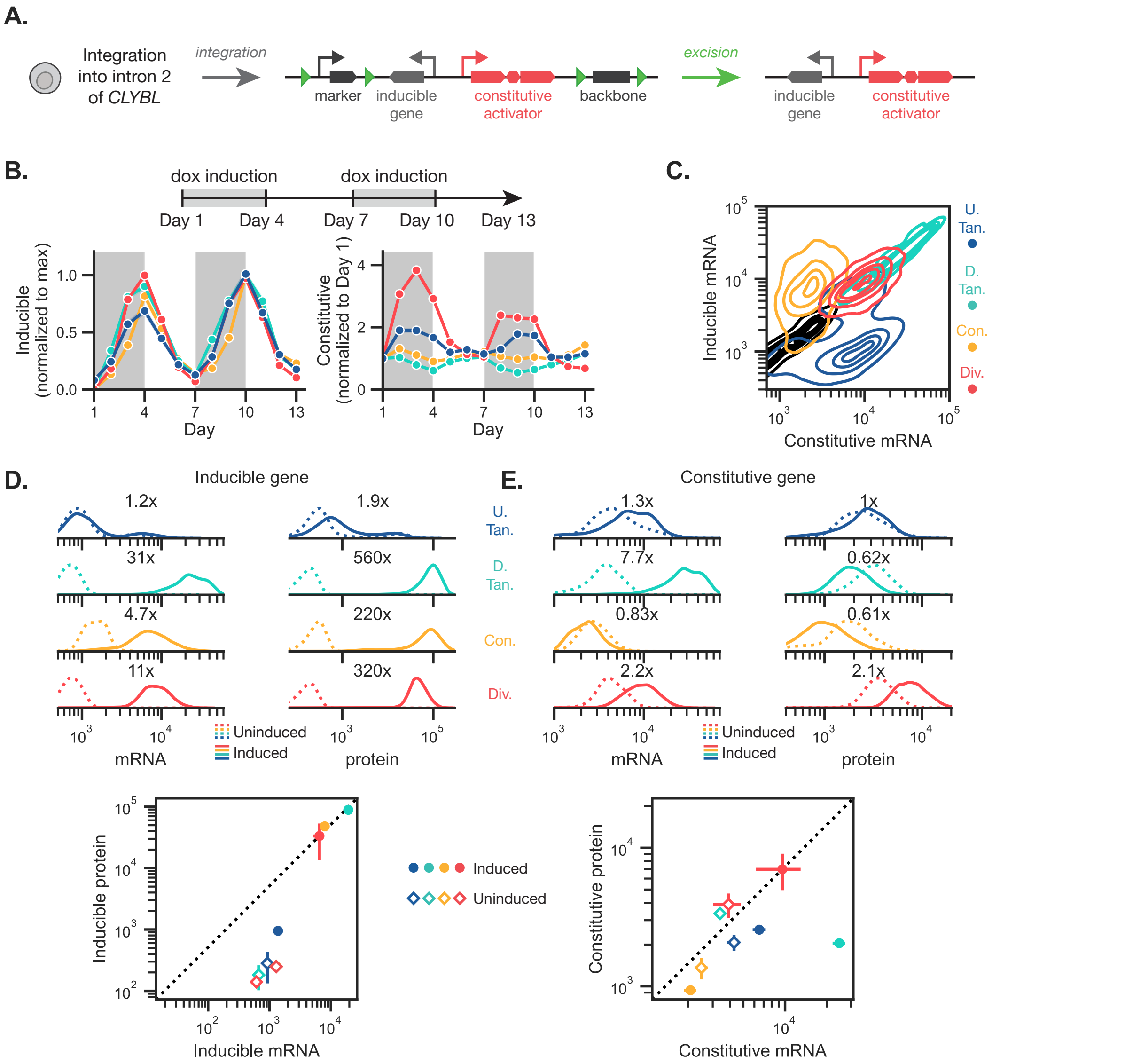


**Figure S10:** Generation and behavior of hiPSC lines with homozygous integration of gene syntax reporters
a) The hiPSC lines were created by integrating a plasmid donor sequentially into each allele of intron 2 of *CLYBL*. The plasmid backbone components and auxiliary elements of the STRAIGHT-IN platform are excised to leave the all-in-one circuit, consisting of an inducible gene expressed from the TRE promoter and a constitutive activator cassette expressed from the CAG promoter.
b) hiPSCs with the circuit integrated at a single allele were repeatedly induced over a period of two weeks. These cell lines display similar syntax-specific trends to the lines in fig. 2e, showing both upstream dominance and divergent amplification. The geometric mean and 95% confidence interval are calculated over three merged wells.
c) HCR RNA FISH combined with flow cytometry was performed on the homozygously integrated hiPSC lines. For a representative bioreplicate, the mRNA joint distributions are shown for each of the four syntaxes, alongside the parental line with no synthetic construct integrated. The downstream tandem syntax shows high correlation, potentially suggesting readthrough.
d) For a representative bioreplicate, the marginal distributions of the inducible mRNA and protein levels is shown. For $n = 3$ bioreplicates, the geometric mean of the mRNA and protein is shown, with the standard deviation as error bars.
e) For a representative bioreplicate, the marginal distributions of the inducible mRNA and protein levels is shown for the constitutive gene. For $n = 3$ bioreplicates, the geometric mean of the mRNA and protein is shown, with the standard deviation as error bars. Due to unproductive transcripts, the induced downstream tandem geometric mean is far from the diagonal.

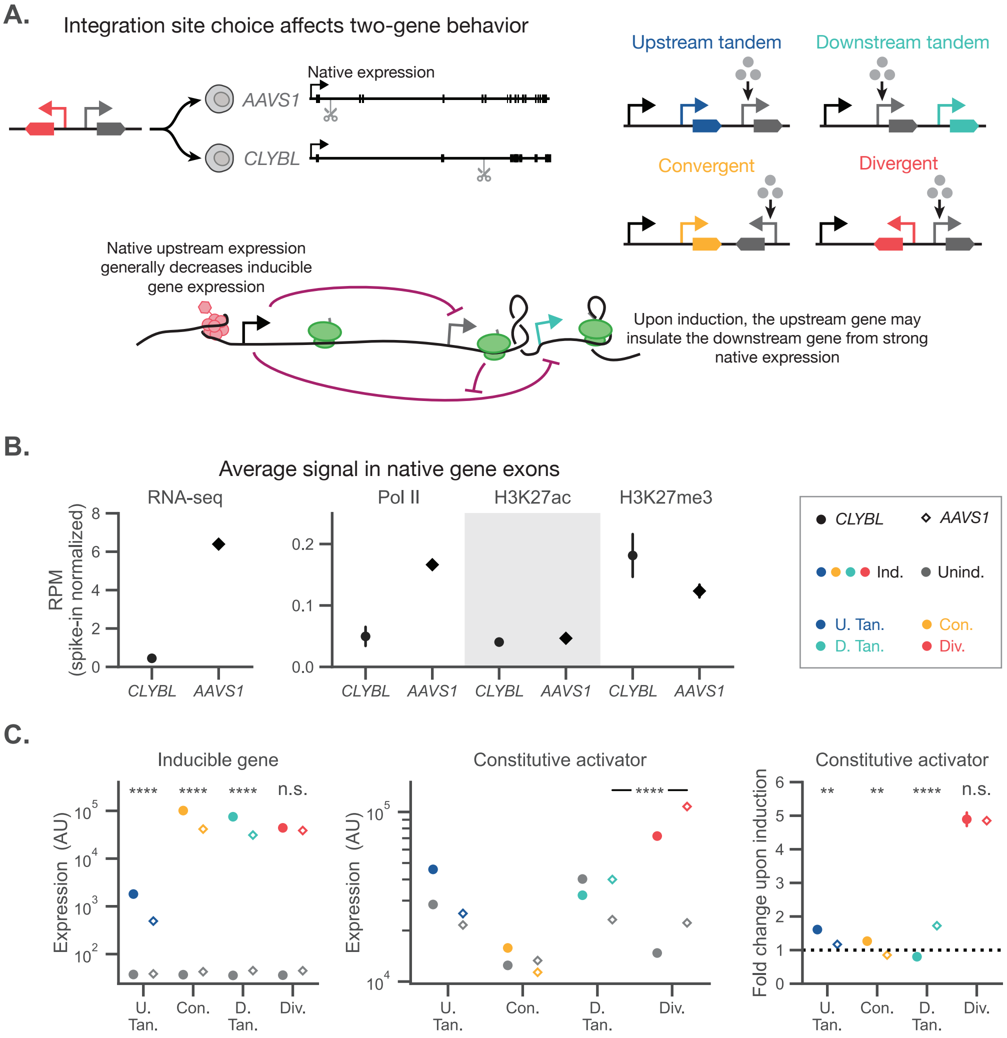


**Figure S11:** Safe harbor integration site choice affects two-gene expression levels.
a) The two safe harbors used in this work, *CLYBL* and *AAVS1*, are located within the introns of two genes. Specifically, this means that native expression from the promoter of these genes may affect the expression of our integrated circuits.
b) Using our genomics datasets, the average signal of RNA-seq reads, actively elongating RNA Pol II and two relevant histone marks are measured and averaged over the annotated exons of the native genes. We find that the *AAVS1* locus is more transcriptionally active and has fewer heterochromatin marks.
c) The expression of the inducible gene is compared for the uninduced and induced case. We find that the expression of the inducible gene is generally lower when integrated into the *AAVS1* locus. The fold change of the activator upon induction is also shown, and generally shows the same trend with the exception of the downstream tandem syntax.

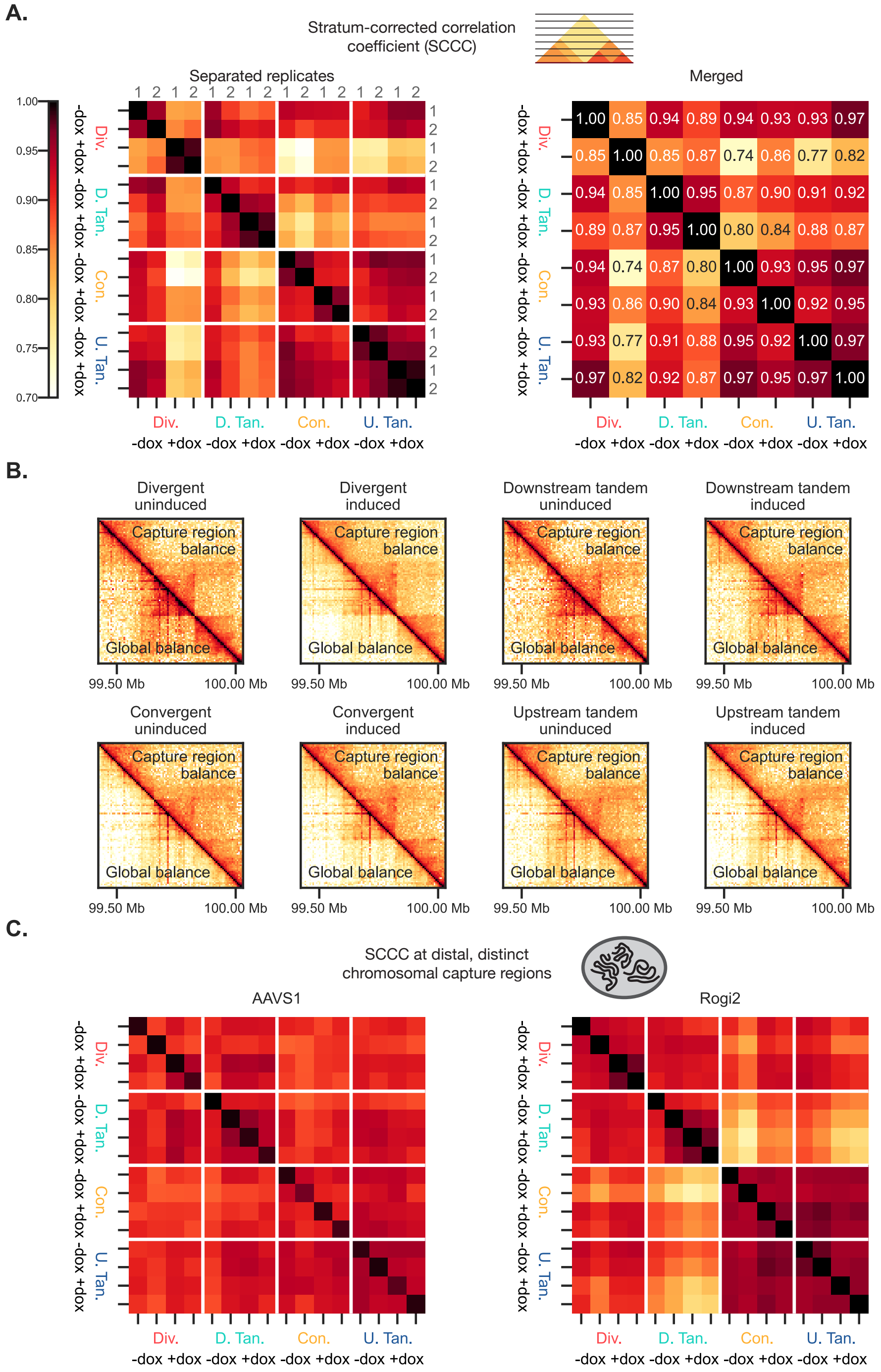


**Figure S12:** Region Capture Micro-C validation.
a) The stratum-corrected correlation coefficient (SCCC) measures the similarity between two interaction matrices while more heavily weighting short-distance genomic contacts. The SCCC is shown for each biological replicate and for the post-merged matrices.
b) The eight resulting merged RCMC genomic contacts were iteratively balanced (i.e., normalized such that every row and column sums to 1 and can be interpreted as a probability distribution), both across the entire genome and only within the capture region. Capture-region balancing provided cleaner contact probability distributions.
c) Two distal regions, *AAVS1*, and Rogi2, were largely unaffected by both integration of the synthetic construct at *CLYBL* and by dox induction, as measured with the SCCC.

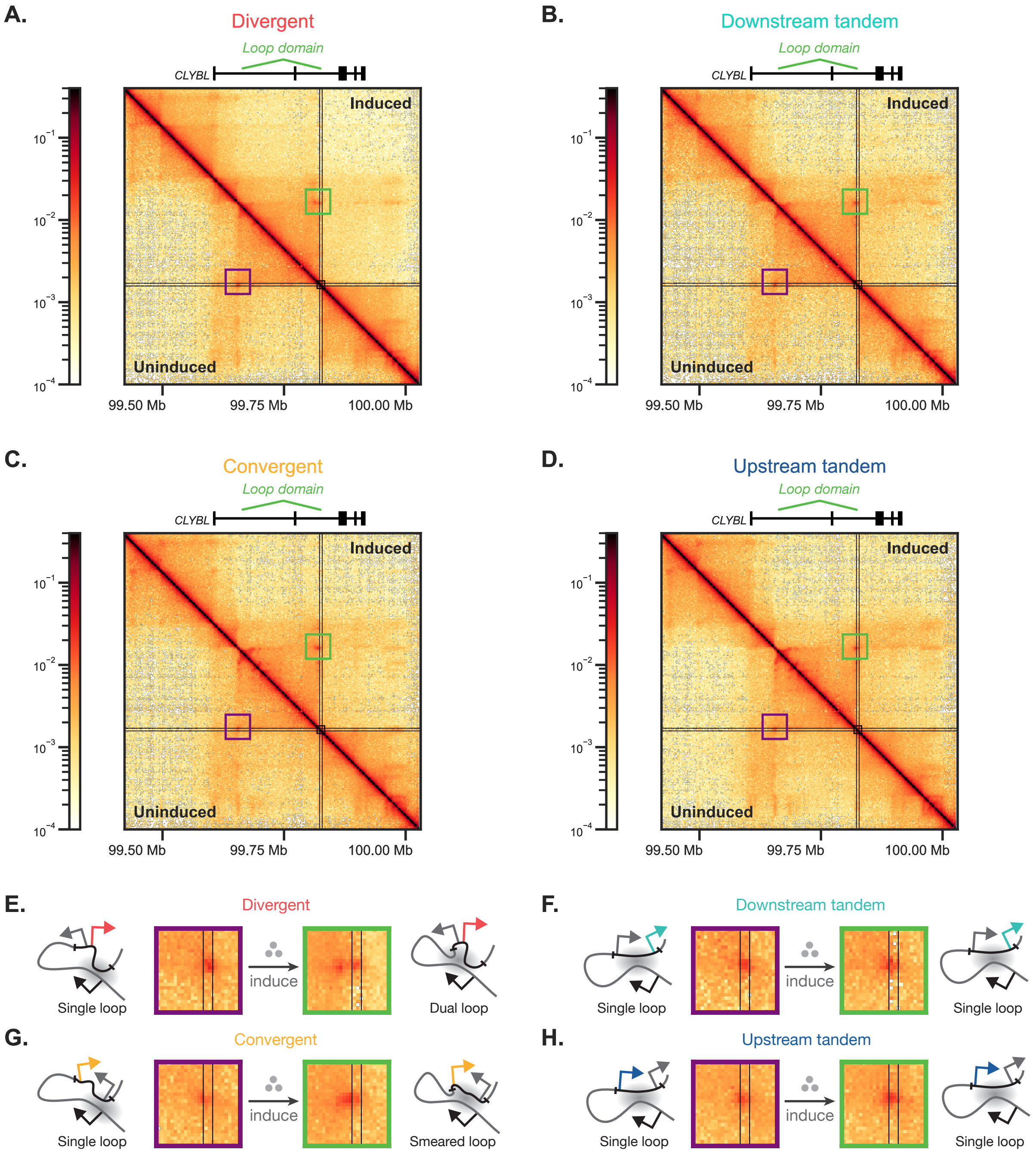


**Figure S13:** Transcriptional perturbation of a 200kb loop domain.
a),b),c),d) Within the capture region, one clear loop domain, identified by a “corner dot” is visible, representing a loop contact between the first intron of *CLYBL* and the integration location.
e),f),g),h) Upon induction, the loop domain is perturbed in both the divergent and convergent syntaxes. This perturbation does not occur in the tandem syntaxes, indicating syntax specificity.

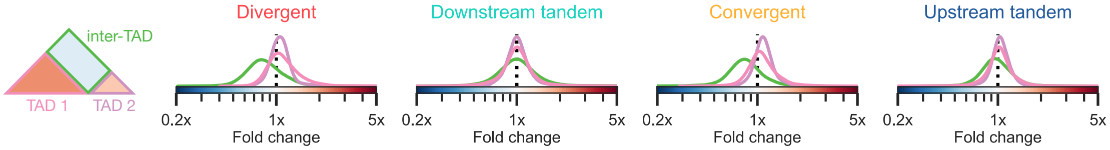


**Figure S14:** Region capture Micro-C TAD interaction distributions.
For the four integrated syntaxes, the change in the inter- and intra-TAD contacts upon induction are shown. The divergent and convergent cell lines show a strong decrease in inter-TAD interactions upon induction, whereas the tandem syntaxes do not show strong TAD interaction changes.

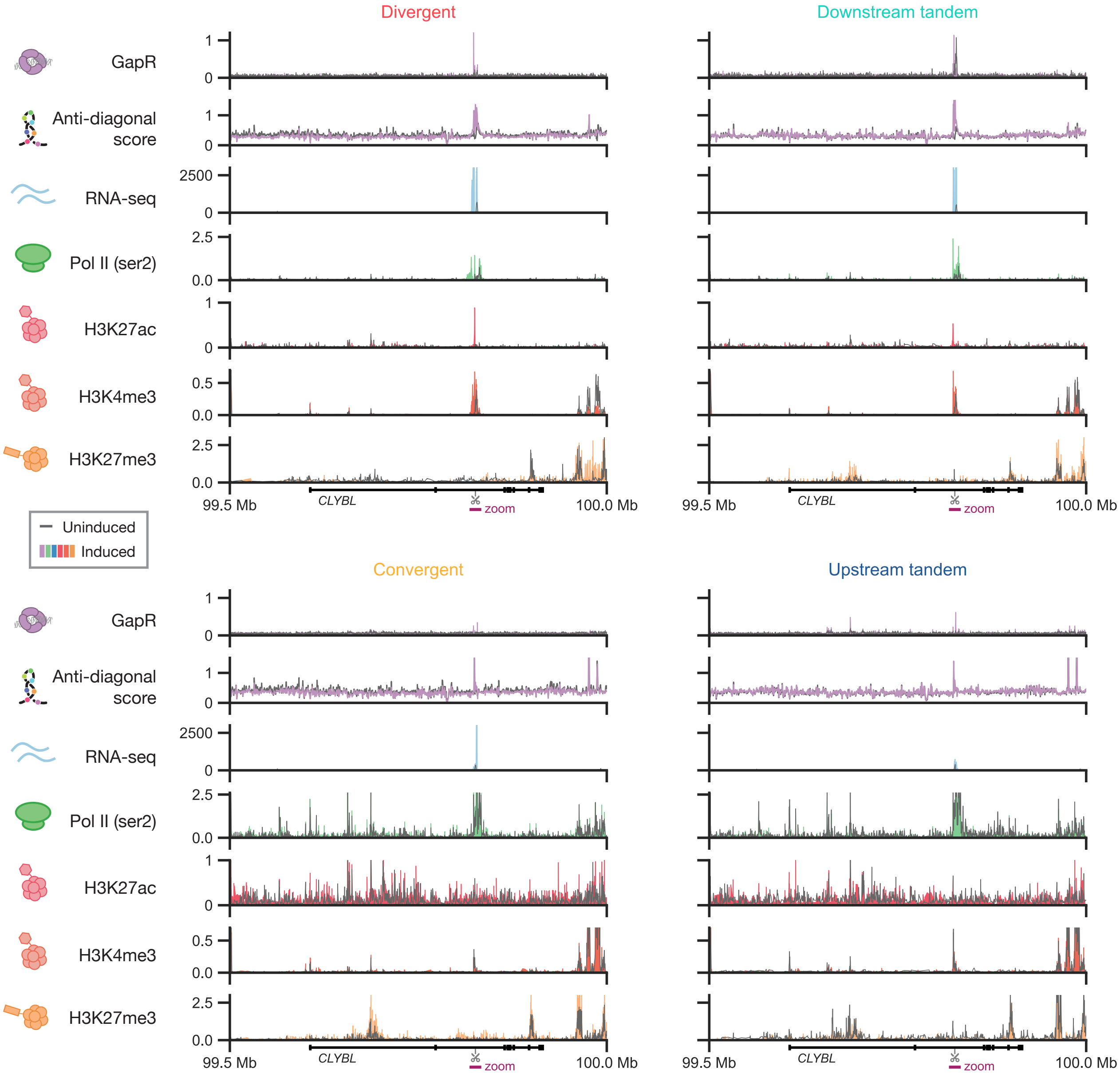


**Figure S15:** 1D chromatin state measured across the entire capture region.
For both the GapRUN and RCMC data (top) and the four CUT&Tag datasets (bottom), the signal across the entire capture region considered in fig. 3 is shown for both the uninduced (gray) and induced (colored) states. The 5kb region surrounding the integration region is shown in gray highlight. While the GapRUN data is not normalized, the RCMC data is internally normalized and the four CUT&Tag datasets are spike-in normalized, with the bottom panels run on a separate normalized

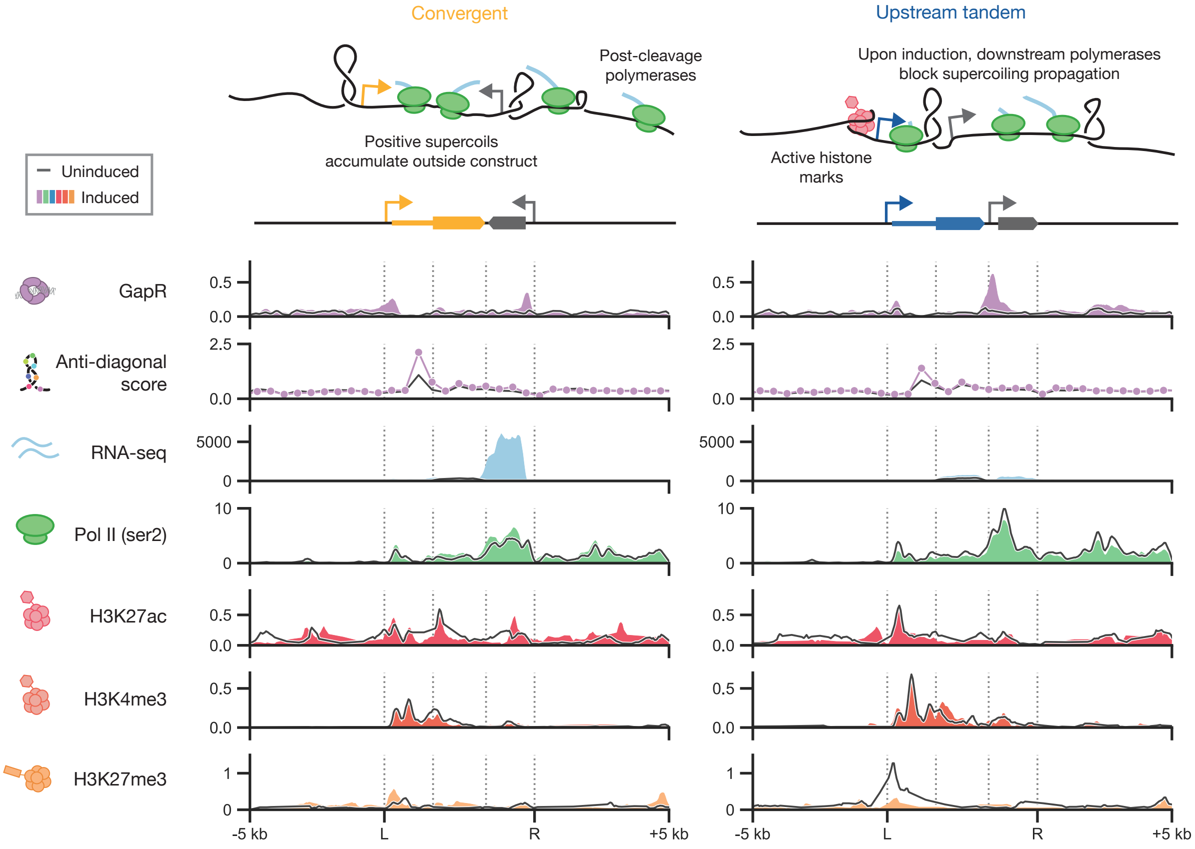


**Figure S16:** Transcription induces syntax-specific chromatin structures in the convergent and upstream tandem syntaxes. While both the convergent and upstream tandem syntaxes show induction-dependent supercoiling signals, only small changes in the histone marks and RNA Pol II occupancy are visible.

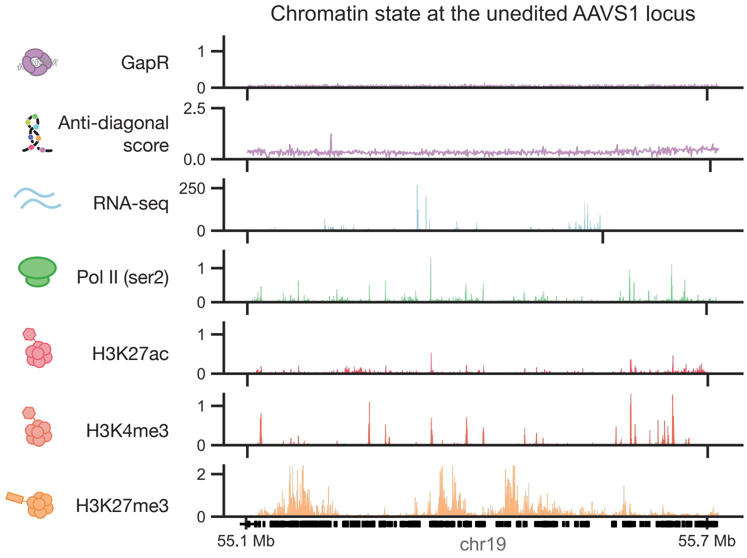


**Figure S17:** Chromatin state at the *AAVS1* locus.
As a comparison to the *CLYBL* locus where synthetic constructs were integrated, the chromatin state at the *AAVS1* locus was examined, for the cell line where the divergent construct was integrated at the *CLYBL* locus. No strong supercoiling signals were observed in any of the syntaxes, while background expression and histone marks were observed throughout the relatively gene-dense *AAVS1* locus.

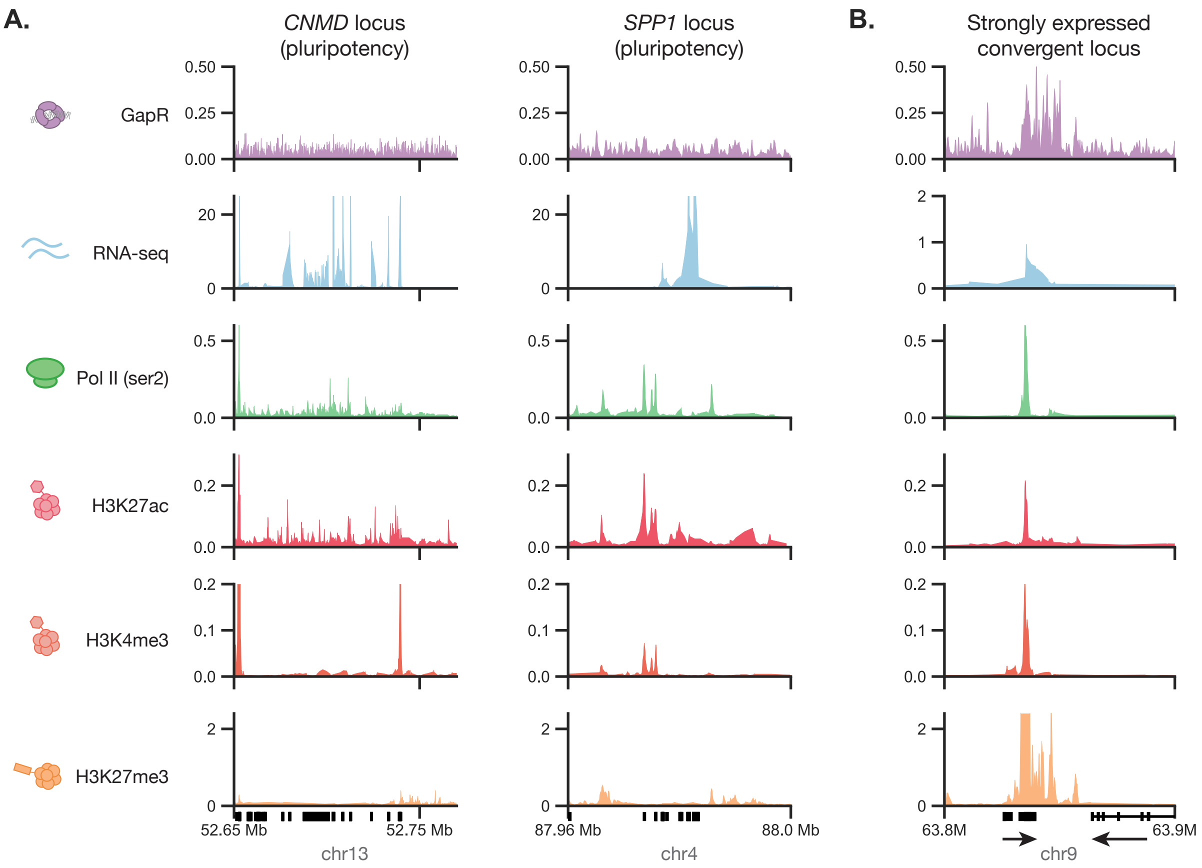


**Figure S18:** Chromatin state at distal regions.
a) Two known marker genes of pluripotency are shown. We observe high transcript counts at these loci. However, due to low expression of the surrounding genes, we see no supercoiling signal above background. b) Identifying native intergene spacers that have high supercoiling signal, we identify a convergent pair of transcripts on chr9, expressing a lncRNA (left) and FRG1JP (right). Strong supercoiling signal is present in the intergenic region.

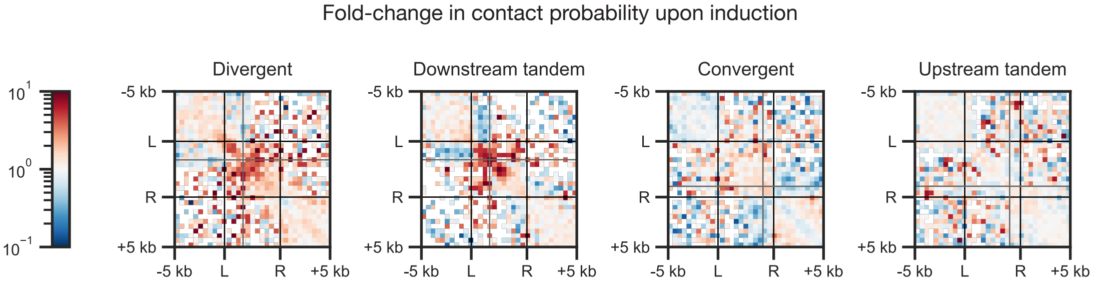


**Figure S19:** The fold change in contact probability is shown for the four syntaxes, in the local region around the integration region. The divergent and downstream tandem circuits show the strongest change in contact probability, though a diffuse increase in anti-diagonal contact probability is still visible in the convergent and upstream tandem syntaxes.

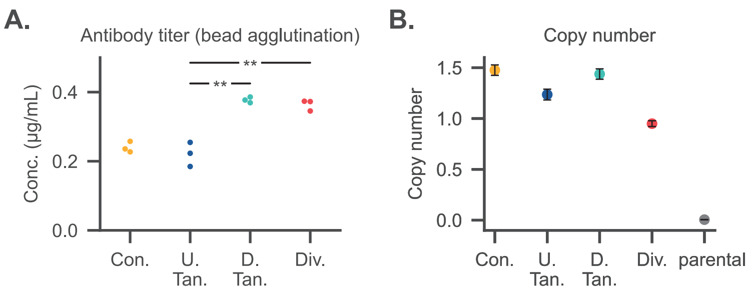


**Figure S20:** a) Antibody titer, as measured by a bead agglutination assay, for cell lines in fig. 5a expressing heavy and light chains with different syntaxes, three biological replicates (separately maintained passages of isolated lines). Statistics are two-sided student t-tests. ** : $p < 0.01$
b) Copy number of the lines presented in fig. 5a, as measured via ddPCR using probes that bind to the mRuby2 coding sequence present downstream of the light chain. Due to genome instability, fractional copy numbers above 1 may occur for HEK293T landing pads.

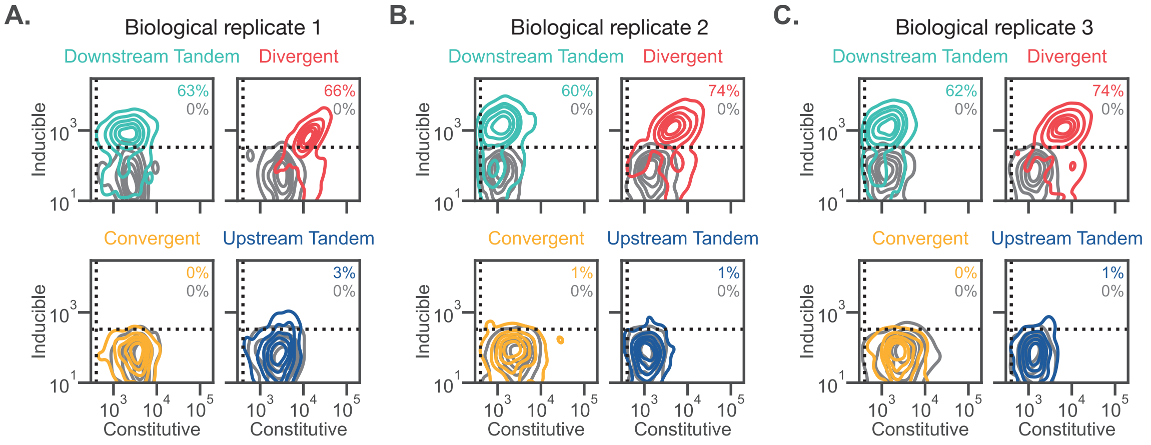


**Figure S21:** Joint distributions for three additional biological replicates of the lentiviral transductions in HEK293T cells presented in fig. 5b.

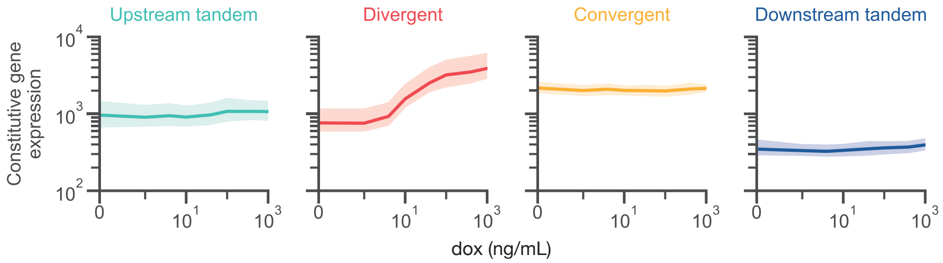


**Figure S22:** Constitutive gene expression as a function of inducer (dox) concentration for the lentiviral transductions of HEK293T cells presented in fig. 5b. Constitutive gene expression remains constant for the tandem and convergent syntaxes but increases with dox for the divergent syntax. Shading represents the 95% confidence interval across four biological replicates.

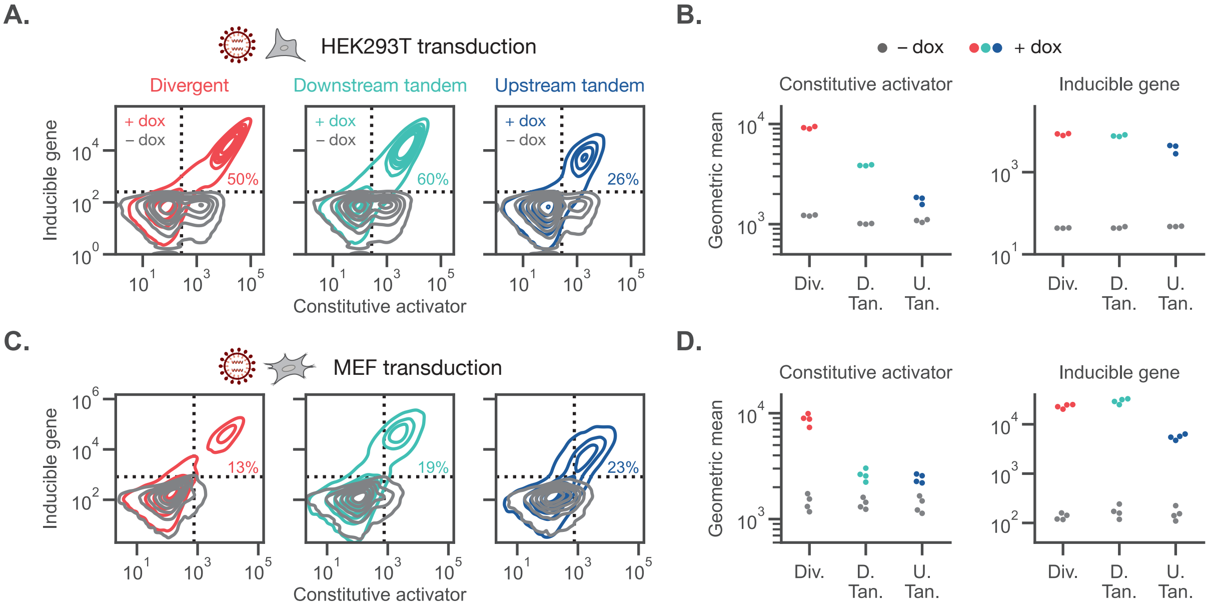


**Figure S23:** Performance of all-in-one inducible circuits across cell types.
a) Joint distributions of the constitutive activator and inducible gene are shown for the inducible all-in-one circuits in fig. 6a. Circuits were lentivirally transduced into HEK293T cells. Distributions show one representative biological replicate.
b) Geometric mean of constitutive activator and inducible gene expression for populations from a) gated on activator-positive cells. The dashed vertical line in a) shows this gate. Points depict three biological replicates.
c) Representative joint distributions of the constitutive activator and inducible gene are shown for the same circuits lentivirally transduced into primary mouse embryonic fibroblasts (MEFs).
d) Data from c) displayed as in b), N=4 biological replicates.

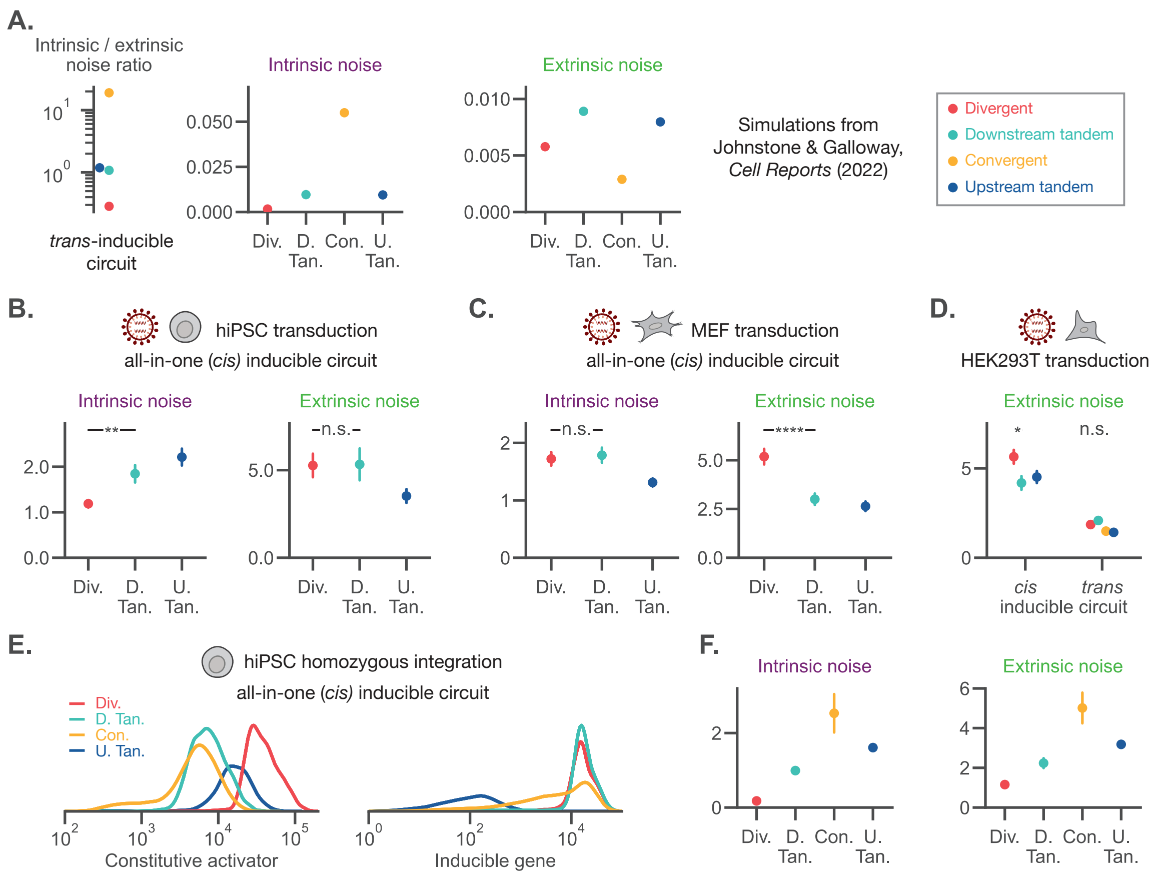


**Figure S24:** Characterization of noise in inducible circuits.

a) For populations of 2,000 simulations of *trans*-inducible circuit modeled in Johnstone & Galloway [7], the intrinsic noise, extrinsic noise, and the intrinsic-to-extrinsic noise ratio is calculated.

b),c) Noise analysis for lentiviral transduction of the *cis*-inducible circuit in human induced pluripotent stem cells (hiPSCs) or mouse embryonic fibroblasts (MEFs) induced with 300 ng/mL dox. Noise was calculated for the double-positive population. Points represent mean ± standard error for N=4 biological replicates.

d) The extrinsic noise component is shown comparing lentivirus transduction of either *cis* all-in-one designs or *trans* designs, as shown in fig. 5b. n=4 biological replicates for the *cis* design and n=3 biological replicates for the *trans* design.

e) For homozygously integrated constructs at the *CLYBL* locus, marginal distributions are shown for the fully induced (1000 ng/mL dox) syntaxes.

f) Performing noise decomposition, the marginal distributions shown are summarized by their noise contribution. The divergent syntax has the lowest intrinsic and extrinsic noise, whereas the convergent syntax has the greatest noise.

Statistics are two-sided student t-tests. n.s.: $p > 0.05$, * : $p < 0.05$, ** : $p < 0.01$

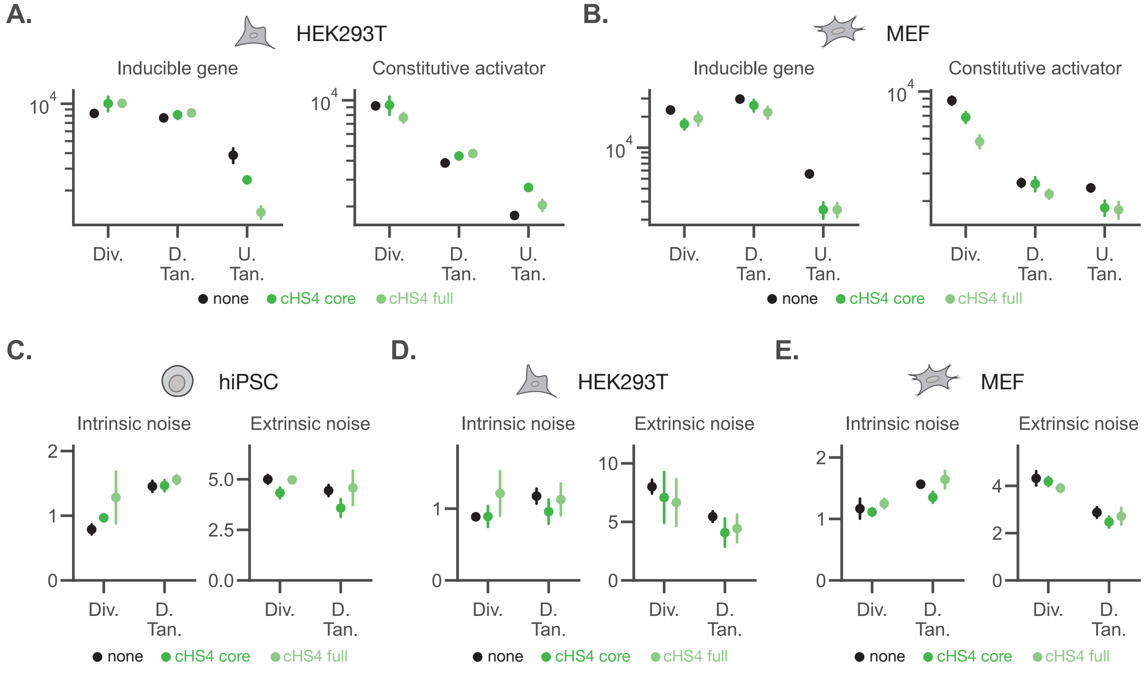


**Figure S25:** The cHS4 insulator does not mitigate the effects of syntax in lentiviral delivery of an inducible, all-in-one circuit.
a) The circuits in fig. 6f were lentivirally transduced into HEK293T cells. Geometric mean expression of the inducible gene and constitutive activator are shown for populations induced with 300 ng/mL dox. Points represent mean ± standard error for N=4 biological replicates.
b) The circuits in fig. 6f were also lentivirally transduced into mouse embryonic fibroblasts (MEFs), n=5 biological replicates.
c) Noise analysis for the circuits transduced into hiPSCs in fig. 6f. All pairwise comparisons between insulators for each syntax are not significant, two-sided student t-tests, $p > 0.05$.
d) Noise analysis for the circuits transduced into HEK293T cells in a). All pairwise comparisons between insulators for each syntax are not significant.
e) Noise analysis for the circuits transduced into MEFs in b). All pairwise comparisons between insulators for each syntax are not significant.

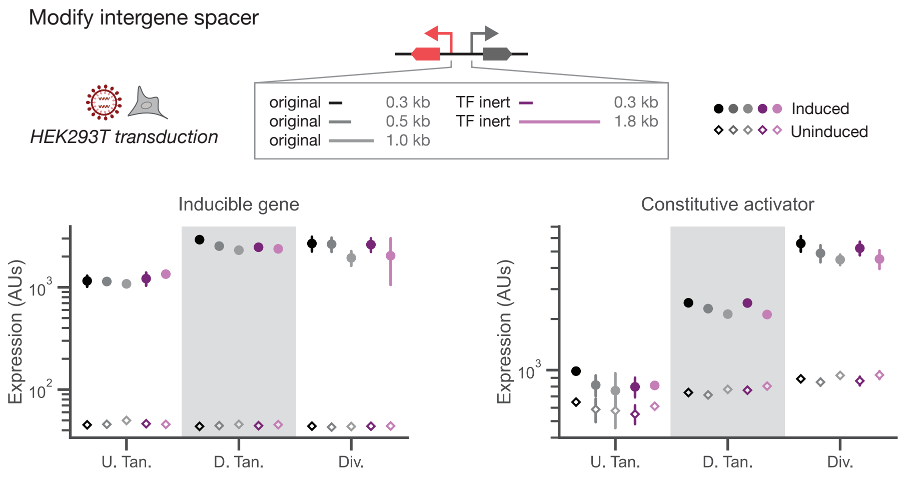


**Figure S26:** Intergene spacer identity only moderately affects lentivirally-induced circuits. Testing three spacers (“original”) that were derived from a frame-shifted, reverse-complement of a coding sequence and two insulators (“TF inert”) designed *de novo* to not contain mammalian TF binding sites, the geometric mean of expression over $n = 3$ biological replicates (separately produced lentivirus) is shown. Spacer identity only mildly changes expression levels. With the exception of the inducible expression for the longest TF-inert spacer, gene syntax determines the relative ordering of construct expression.

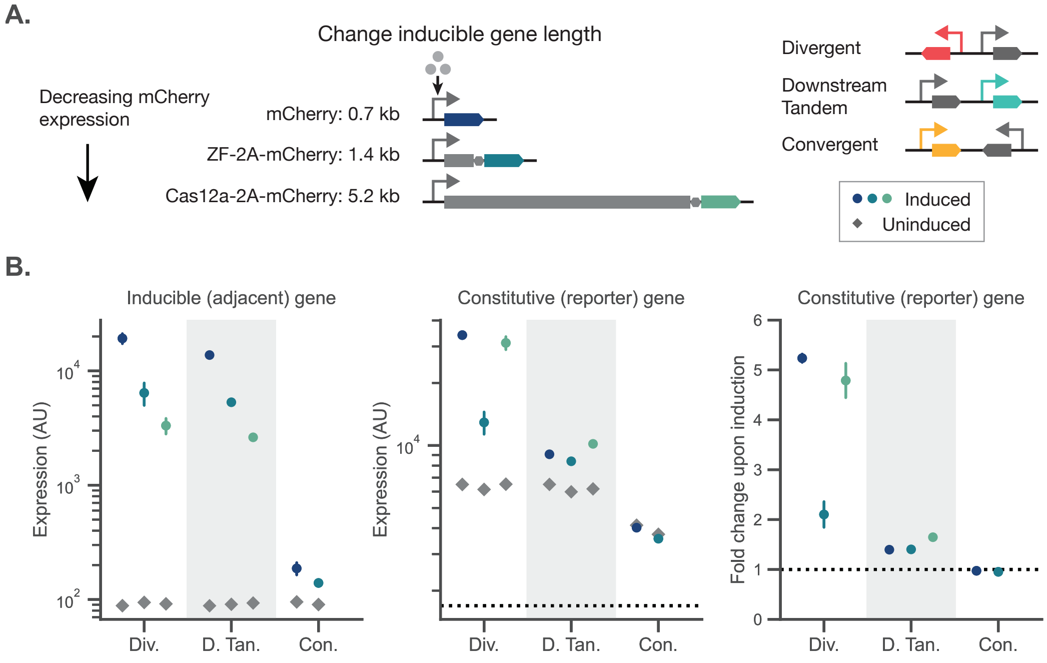


**Figure S27:** Inducible gene length affects expression non-monotonically.
a) To investigate the relationship between gene length and supercoiling-mediated biophysical coupling, we integrated three different inducible genes, all containing the same fluorescent protein mCherry, into the *CLYBL* safe-harbor locus. Cells expressing the constitutive reporter above a 95% percentile gate of the parental line were gated. Conditions with a small fraction of cells in this gate were excluded.
b) The expression of the inducible gene is shown as a function of circuit syntax and gene length. For the highly expressed genes, increasing the gene length decreases expression.
c) The expression of the constitutive gene is shown as a function of syntax and inducible gene length. We observe that, for all syntaxes, the constitutive gene is least affected by the intermediate (1.4 kb) gene length, suggesting a non-monotonic tradeoff between supercoiling generation, polymerase loading, and polymerase stalling. Error bars show the standard error for $n = 3$ bioreplicates (separate splits of singly-integrated hiPSCs).

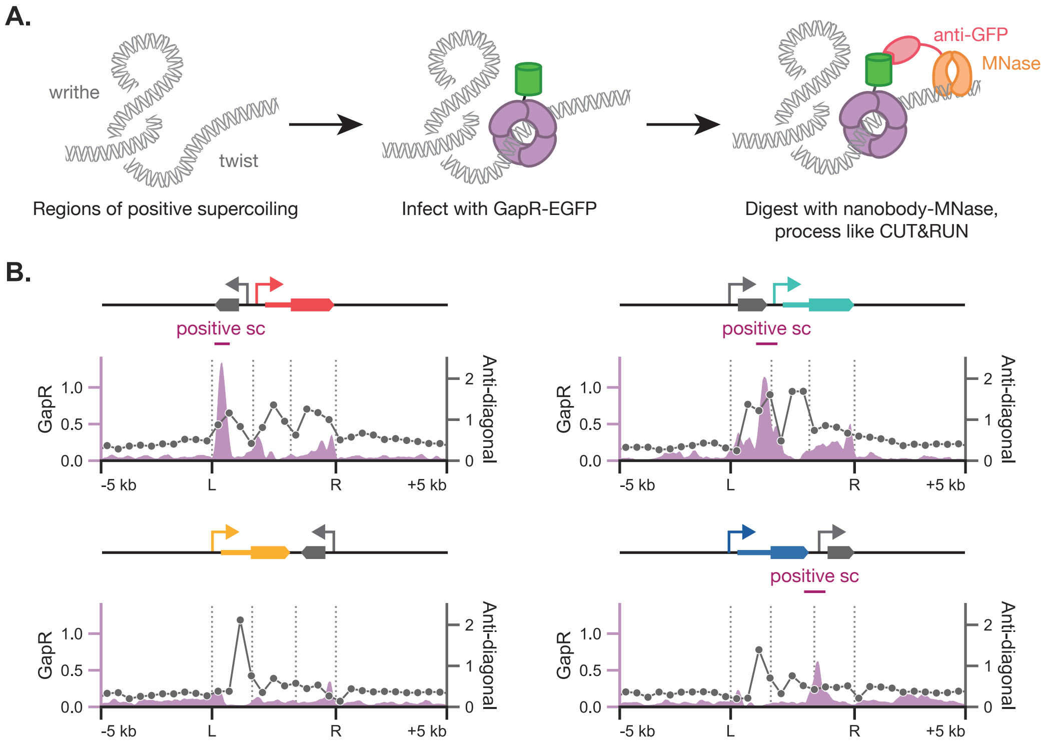


**Figure S28:** Measuring positive supercoiling with GapR

a) Positive supercoiling interconverts between twist, average rotation per base pair, and writhe, large-scale intertwined loops. Positive twist can be bound by a GapR-EGFP fusion protein that is integrated via lentivirus and expressed from the genome. Using a nanobody targeting GFP fused to MNase, the genome can be selectively digested where the GapR protein bound, resulting in fragments that can be processed as in CUT&RUN.

b) The GapR signal (solid purple) is compared to the anti-diagonal score (black line with purple markers). Regions of high anti-diagonal score overlapping with GapR signal suggests the presence of positively supercoiled plectonemes.

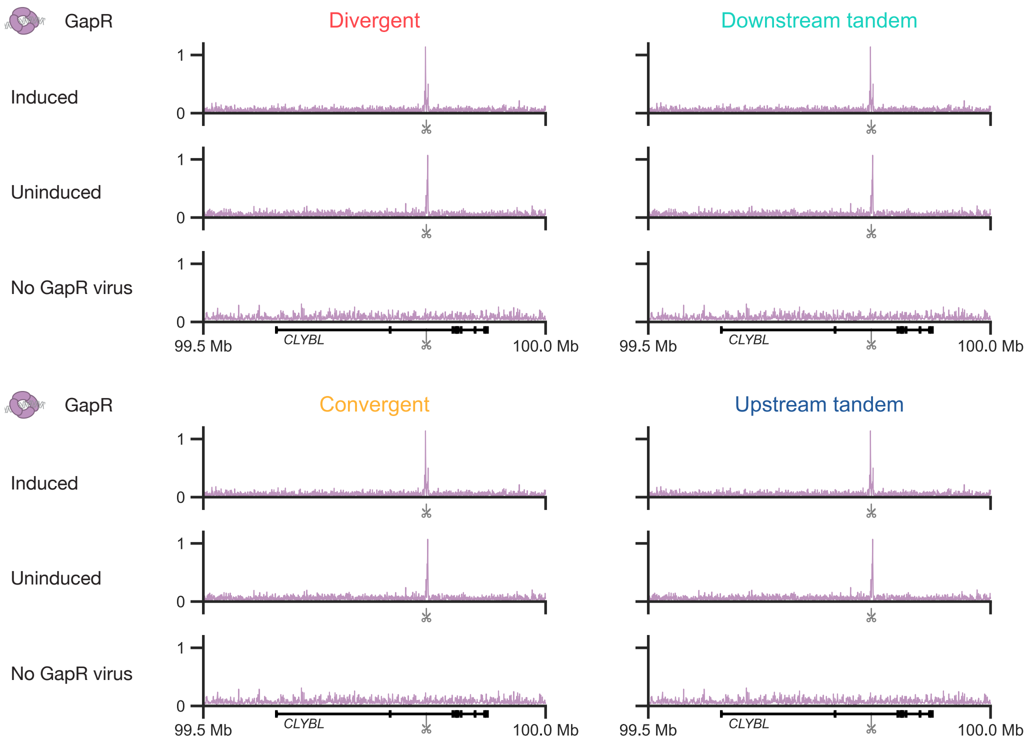


**Figure S29:** The positive twist signal is only present when the GapR virus is present. The GapR signal across the integration locus is shown for three cases: GapR-transduced and dox-induced, GapR-transduced and uninduced, and uninduced cells that were not transduced with the GapR virus. The untransduced, no-virus control cells only show background levels of supercoiling signal.
